# Turbulent adaptive landscape shaped size evolution in modern ocean giants

**DOI:** 10.1101/2022.09.07.506945

**Authors:** Gustavo Burin, Travis Park, Tamora D. James, Graham J. Slater, Natalie Cooper

**Author notes:** these authors contributed equally to this work (Gustavo Burin, Travis Park).

## Abstract

Adaptive landscapes are central to evolutionary theory, forming a conceptual bridge between micro- and macro-evolution^1–4^. Evolution by natural selection across an adaptive landscape should drive lineages towards fitness peaks, shaping the distribution of phenotypic variation within and among clades over evolutionary timescales^5^. Constant shifts in selection pressures mean the peaks themselves also evolve through time^4^, thus a key challenge is to identify these ‘ghosts of selection past’. Here, we characterise the global and local adaptive landscape for total length in cetaceans (whales and dolphins) across their ~ 53 million year evolutionary history, using 345 living and fossil taxa. We analyse shifts in long-term mean size^6^ and directional changes in average trait values^7^ using cutting-edge phylogenetic comparative methods. We demonstrate that the global macroevolutionary adaptive landscape of cetacean body size is relatively flat, with very few peak shifts after cetaceans colonised the oceans. Local peaks represent trends along branches linked to specific adaptations such as deep diving. These results contrast with previous studies using only extant taxa^8^, highlighting the vital role of fossil data for understanding macroevolutionary dynamics. Our results indicate that adaptive peaks are constantly changing and are associated with subzones of local adaptations, resembling turbulent waters with waves and ripples, creating moving targets for species adaptation. In addition, we identify limits in our ability to detect some evolutionary patterns and processes, and suggest multiple approaches are required to characterise complex hierarchical patterns of adaptation in deep-time.

## Introduction

The metaphor of the adaptive landscape provides a compelling framework for understanding patterns of phenotypic evolution. Originating with Sewall Wright’s^1^ visual explanation for changes in gene frequencies in populations under selection, modern theories of morphological evolutionary landscapes owe more to Simpson’s^2, 3^ reformulation of them in terms of zones in an adaptive grid^4^. Simpson considered adaptive zones to represent phenotypic solutions to functional problems, and the development of mathematical models for describing the evolution of phenotypes towards adaptive zones^9^ has led to a diversity of approaches for characterising phenotypic adaptive landscapes from phylogenetic comparative data^6, 10, 11^. Although these approaches assume that the adaptive landscape is composed of discrete peaks or zones, a fundamental insight from Simpson’s conceptual models was that adaptive zones are hierarchically organised (see also the distinction between global and local perspectives in 4). The location and breadth of zones and subzones in phenotypic space can themselves also evolve through time^4^, but whether phylogenetic comparative methods are able to detect such patterns has largely remained unexplored^12^.

Here, we combine a novel dataset of total length estimates with a time-scaled phylogeny, to quantify the global and local adaptive landscapes for cetacean (whales, dolphins, and relatives) body size over their entire ~ 53 million year evolutionary history. Cetacean body size is a powerful trait for such a study, spanning five orders of magnitude and including the largest animal to have ever lived, the 30 m long, 180,000 kg blue whale (*Balaenoptera musculus*). It has been previously suggested that body size variation in cetaceans is the outcome of distinct evolutionary trajectories or peaks on a rugged adaptive landscape that are associated with foraging behavior^8^ or physiological constraints on life in aquatic environments^13, 14^. However, the broader context for patterns of body size evolution in the clade and the underlying macroevolutionary adaptive landscape for cetacean body size have been neglected until now, particularly suffering from a lack of fossil taxa included in analyses (see 15, 16 for preliminary studies). This broader context is vital as it can shed light not only on clade-specific hypotheses in groups that are yet to be examined in detail, but also on the critical role that fossil data has in improving the accuracy of macroevolutionary studies^17–19^. Remarkably, we find that the global macroevolutionary adaptive landscape of cetacean body size is relatively flat, with only a handful of peak shifts occurring after the initial colonisation of the marine environment. Local peaks are more numerous, however, and manifest as trends along branches linked to more specific adaptations such as deep diving, macro-predation and bulk filter-feeding.

## Results

### Total length data imputation

By using phylogenetically-informed imputations and a range of proxy measurements (see Methods), along with total length information from the literature and museum specimens, we generated the largest total length dataset to date for Cetacea, comprising 345 taxa (89 living and 256 fossil; Figure 1). Where total lengths were known, imputed values showed a strong correlation with them, regardless of the amount of information used for estimation (see Supplementary Information; Figures S1-S4), meaning that total lengths could be estimated for even incomplete fossil taxa (Figures S1-S4), allowing their inclusion in downstream analyses. These additional data are critical to robust macroevolutionary inference - as in previous work^8, 20^, we recover a significantly more bottom-heavy disparity profile for extant cetaceans than would be expected under a constant-rates process (centre of gravity = 0.43, *P* = 0.005), consistent with early burst adaptive radiation models (Figure S5). However, this signal is entirely erased by the inclusion of fossil taxa, regardless of whether we include all extinct cetaceans or limit our sample to crown Neoceti only (centre of gravity = 0.29, *P* = 0.294).

**Figure 1.**
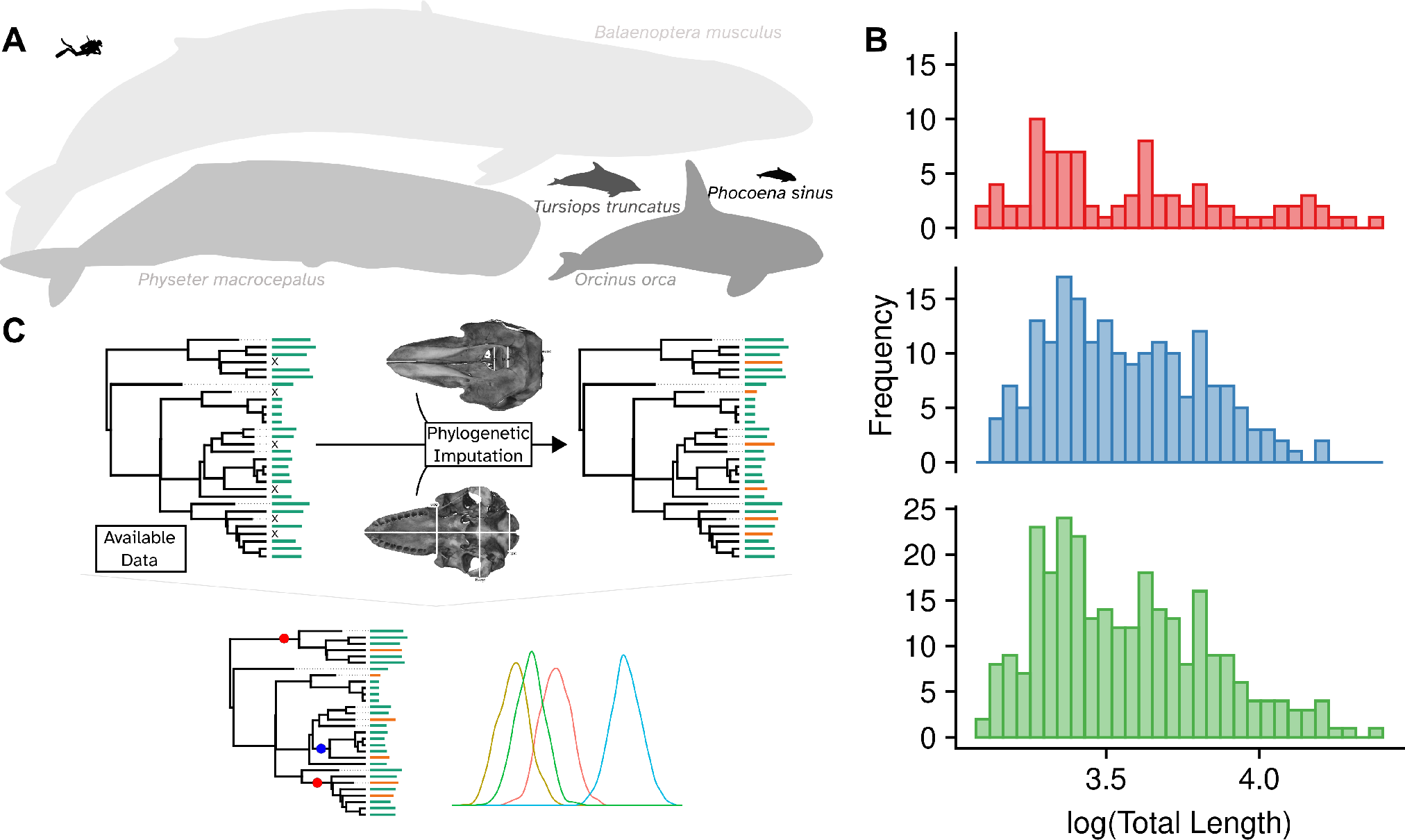
Total length variation in cetaceans and the analytical approach used in the analyses. A: Silhouettes of five species of cetacean highlighting the variation in total length within the group (diver silhouette to scale). B: Histograms showing the distribution of log_10_-transformed total length (m) for living (red bars/upper panel), extinct (blue bars/middle panel) and all cetacean species (green bars/bottom panel). C: Schematic representation of the analytical approach used in the paper. We used data from the literature, new measurements of museum specimens, and the phylogeny of^21^ to estimate total length values for species lacking these data. The imputed dataset and phylogeny were then analysed using `bayou`^6^ to detect shifts in the evolutionary regimes of total length, generating insights about the adaptive landscape of this trait.

### Exploring tempo and mode in body size evolution

The macroevolutionary adaptive landscape is often conceptualised as an *n*-dimensional topographic map, the ruggedness of which depicts the relationship between morphological form and fitness^4^. If topographic relief of the landscape is sufficient, selection should cause traits to evolve rapidly towards peaks and remain there until selection relaxes or the topography of landscape is altered, for example due to environmental disturbance or changes in the availability of resources. The topography of the adaptive landscape can be approximated as an Ornstein-Uhlenbeck process, where the positions of peaks on the landscape are represented as a vector of distinct long-term means, *θ* = (*θ*_1_,*θ*_2_, …, *θ_n_*), and the ruggedness of the landscape is represented by the magnitude of a parameter, *α*, that determines the strength of the pull towards the peaks as an exponential function of time^22^. As *α* → 0, the ruggedness of the adaptive landscape declines and evolution reverts to a random walk. Such a result would not necessarily mean that a trait does not experience strong selection over short (e.g. generational) timescales; rather, it would imply that there are no long-term active trends in trait evolution at the clade level.

We used reversible jump Markov Chain Monte Carlo methods^6^, to map the adaptive landscape for cetacean body sizes. An advantage of this approach is that does not require the user to specify the location of peaks *a priori*, rather they are estimated from the data. Estimated peaks differed by an order of magnitude, ranging from approximately 2m up to 12m, although the 95% credibility interval for the estimated peaks indicated that this variation could be even higher, ranging from 60cm to 50m (see Figure S6).

It has long been recognised that secondarily aquatic tetrapods tend to be larger than their terrestrial sister lineages^13^. Diverse explanations have been suggested for this pattern. Life in water frees the limbs from mechanical constraints associated with supporting body mass^23^, which should passively increase the upper limit on potential sizes without imposing a minimum size constraint. Conversely, heat is more readily lost in water than in air and, because surface area is reduced relative to volume in larger animals, this should place constraints on the lower size limits of fully aquatic taxa^24^. Our results indicate that the cetacean adaptive landscape for body size was profoundly shaped by the thermoregulatory lower limit on size in marine environments^13^, with a dramatic shift to a new adaptive peak associated with large (posterior mean = 12m total length) size during the major transitional stage of adapting to fully aquatic life in ambulocetids, from a small (posterior mean = 2m) ancestral size.

Body size variation is often viewed as a critical axis of trophic niche partitioning, enabling the coexistence of ecologically similar and phylogenetically related taxa^25^. Although this relationship is most typically studied in the context of macroecological questions^26^, where the size distribution of prey resources is discontinuous, size-based dietary niche partitioning could potentially yield novel peaks on the adaptive landscape for predator size. Indeed, among marine predators, mean, minimum, and maximum prey size all show strong positive correlations with predator size^27^, suggesting that this is a plausible axis of macroevolutionary diversification.

We found limited support for a trophically defined adaptive landscape for body size in cetaceans, *contra* previous work on extant taxa alone^8^. The cetacean body size adaptive landscape remains remarkably stable for ~ 20 million years after the initial colonisation of the marine realm. However, a dramatic decrease in long-term mean body size (from ~ 12.5m to ~ 2.9m) in the Late Oligocene along the branch leading to Delphinida (oceanic dolphins, porpoises, and river dolphins), a clade of predominantly piscivorous odontocetes, with a second shift to even smaller sizes (long-term mean ~ 2.1m) in the Early Miocene along the branch leading to total group Inioidea + Pontoporiidae does fit this model. Maneuverability is negatively correlated with body size in marine mammals^28^ and evolution about these smaller long-term means would provide a performance advantage for delphindans when foraging for small, agile prey in complex environments. It is tempting to speculate that the second decrease in long-term mean body size seen in the Lipotidae + Inioidea clade could be an adaptation to living in freshwater river systems, given their extant distributions. However, this modern distribution is convergently derived, and many fossil taxa in the clade are found in marine sediments^29^. Functional studies of *Inia* indicate that its flexible body enhances manoeuvrability at the expense of speed^28^. Thus, members of this lineage may have attained small body sizes to optimally maneuver in shallow coastal environments, in turn pre-adapting themselves for life in rivers. Delphinoids, in contrast, exhibit a suite of vertebral adaptations that result in a stiffer body^30^, which sacrifices some of the manoeuvrability attained by iniids and pontoporiids for increased speed and swimming efficiency^28^. This likely facilitated the diversification of pelagic piscivores during the Late Miocene, culminating in the dramatic, rapid radiation of oceanic dolphins that today comprise almost half of cetacean diversity^8, 21, 31^.

Aside from these major shifts, we detected few other peaks on the adaptive landscape that affect large clades or that are associated with substantial size change. Shifts within odontocetes represent increases in body size in relation to the Delphinida mean, and are detected for Kentriodontidae and closely-related stem delphinidans, and for *Orcaella* (Figure 2 and Figure S7). Within the baleen whales (Mysticeti), we only observed shifts within the crown group (Figure 2). These shifts indicate localised decreases in body size within balaenids and in ‘basal thalassotherians’ (*sensu*^32^), with other lineages of balaenopterids containing the remaining changes (Figure 2 and Figure S7). We found no evidence for shifts towards gigantic sizes in crown Balaenopteridae or Balaenidae, the largest extant cetaceans, but we did find a shift towards smaller size in the minke whale. Virtually all the shifts described above were retained across our sensitivity analyses (see Methods and Supplementary Information) regardless of the tree and prior parameters used (Supplementary Information; Figures S8-S58).

**Figure 2.**
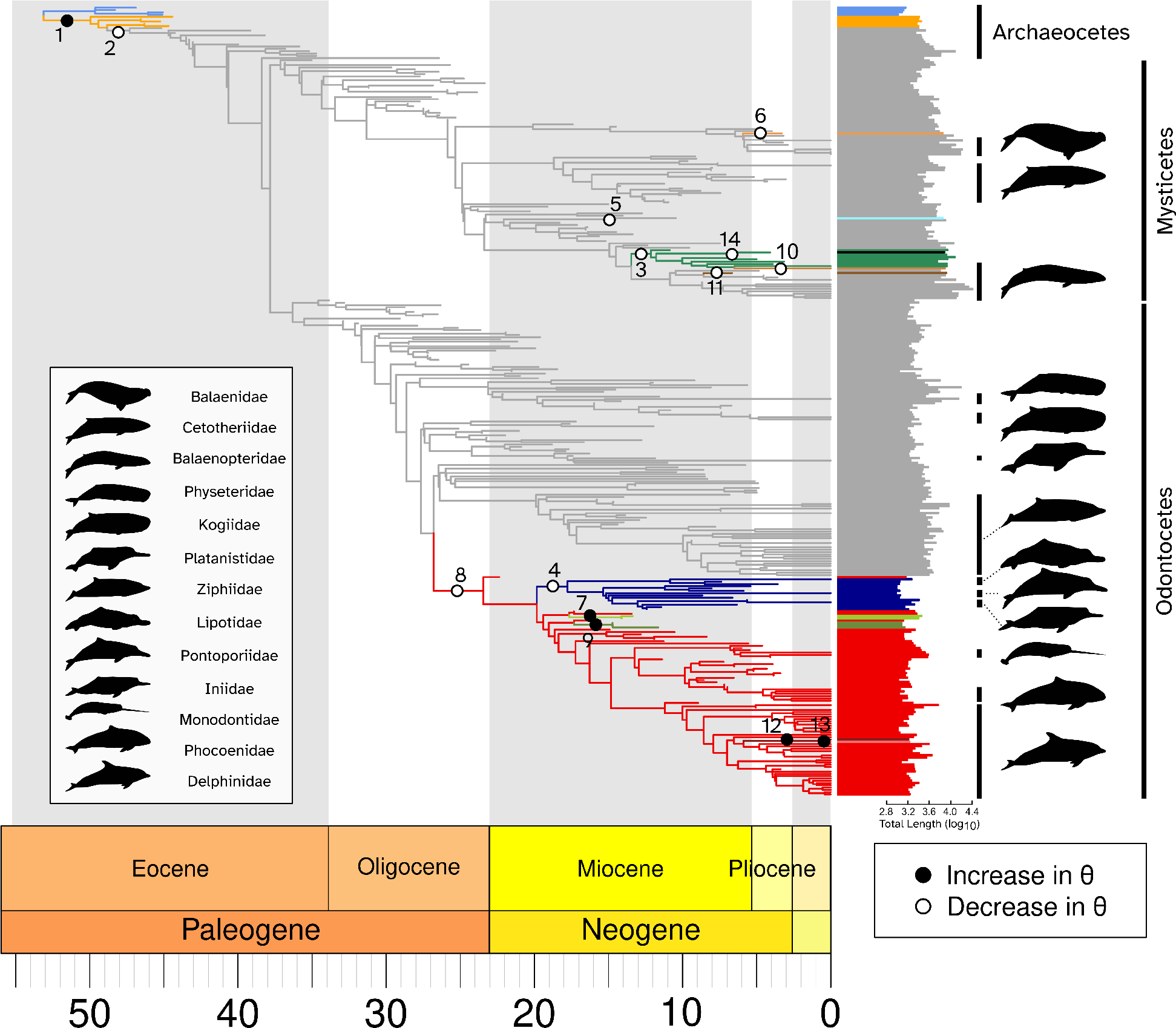
Evolutionary shifts in total length evolution of living and extinct cetaceans. Phylogeny of cetaceans showing the 14 shifts detected by bayou^6^. Solid black circles represent increases in average total length (*θ*) for a given regime; hollow circles represent decreases in average total length (*θ*) for a given regime. The bars to the right of the phylogeny show the log_10_-transformed total length values (m) for each species. The branches and their corresponding bars are coloured according to the regime they belong to. Silhouettes indicate cetacean families that have at least one living species.

Although the fit of Ornstein-Uhlenbeck models to comparative data is often interpreted in terms of estimates of *θ*, the location of the adaptive peak(s), the parameter *α* that describes the topography of the landscape and thus the historical strength of selection, is of equal importance^33^. Estimates from wild populations indicate that selection can drive incredibly rapid responses in heritable phenotypes within a single generation^34^, though this may represent short-term fluctuations in response to changes in local conditions^35^ that do not accumulate to yield large macroevolutionary divergences^12^. Still, for cetacean total length we estimate an extremely low rate of adaptation (posterior mean *α* = 0.0077, 95% Highest Posterior Density = 0.0001 –0.0141), indicating that it would take, on average, approximately 90 million years (or 1.6 times the age of the entire clade) to move halfway towards a new long-term mean body size after a peak shift. Such low rates of adaptation are, at face value, incompatible with the view that selection drives the evolution of body size across the adaptive landscape.

A flat adaptive landscape for cetacean body size implies that variance should have increased through time, yielding both larger and smaller taxa in spite of an inferred mean size at the upper end of the body size distribution. However, if our models are insufficient, then it is possible that our low parameter estimates reflect more complex underlying dynamics rather than weak selection. For example, the surprising lack of peak shifts identified in our bayou analyses, particularly in large-bodied extant taxa, could result from that model’s inability to accommodate multiple *α* values, which could cause us to miss nested shifts associated with periods of accelerated evolution towards a new local peak, or from small sample sizes (such as where shifts occur along terminal branches) that preclude parameter identifiability. To account for these possibilities, we fitted a simpler phenomenological model to our data in which phenotypic shifts along individual branches of phylogeny are treated as biased deviations from an otherwise unbiased random walk where rates are free to vary^7^. Any trends towards larger or smaller sizes can be thought of as evolutionary change that falls outside of the range of possibilities permitted by an otherwise unbiased process, though no explicit mechanistic explanation can be invoked.

We identified 18 branches along which evolution was biased away from an expected change of 0, none of which were found by `bayou`^6^ (Figure 3). Eleven of these shifts were towards larger sizes. Notably, some of these involved macroraptorial predators *Basilosaurus*, *Ankylorhiza tiedemani*, *Livyatan melvillei*, and the modern orca *Orcinus orca*. A massive (175%) increase in size was also detected along the branch leading to crown group balaenopterids (excluding minkes). Additional increases were detected in crown balaenids, and the deep-diving ziphiids. Seven trends towards smaller size were also recovered. For example, the ancestor of total group odontocetes underwent a substantial (35%) decrease in size relative to the neocete ancestor, while smaller decreases occured in the common ancestor of the pygmy sperm whale family Kogiidae, several odontocete terminal lineages, and a pair of nested trends towards smaller size in the toothed mysticete lineage Aetiocetidae.

**Figure 3.**
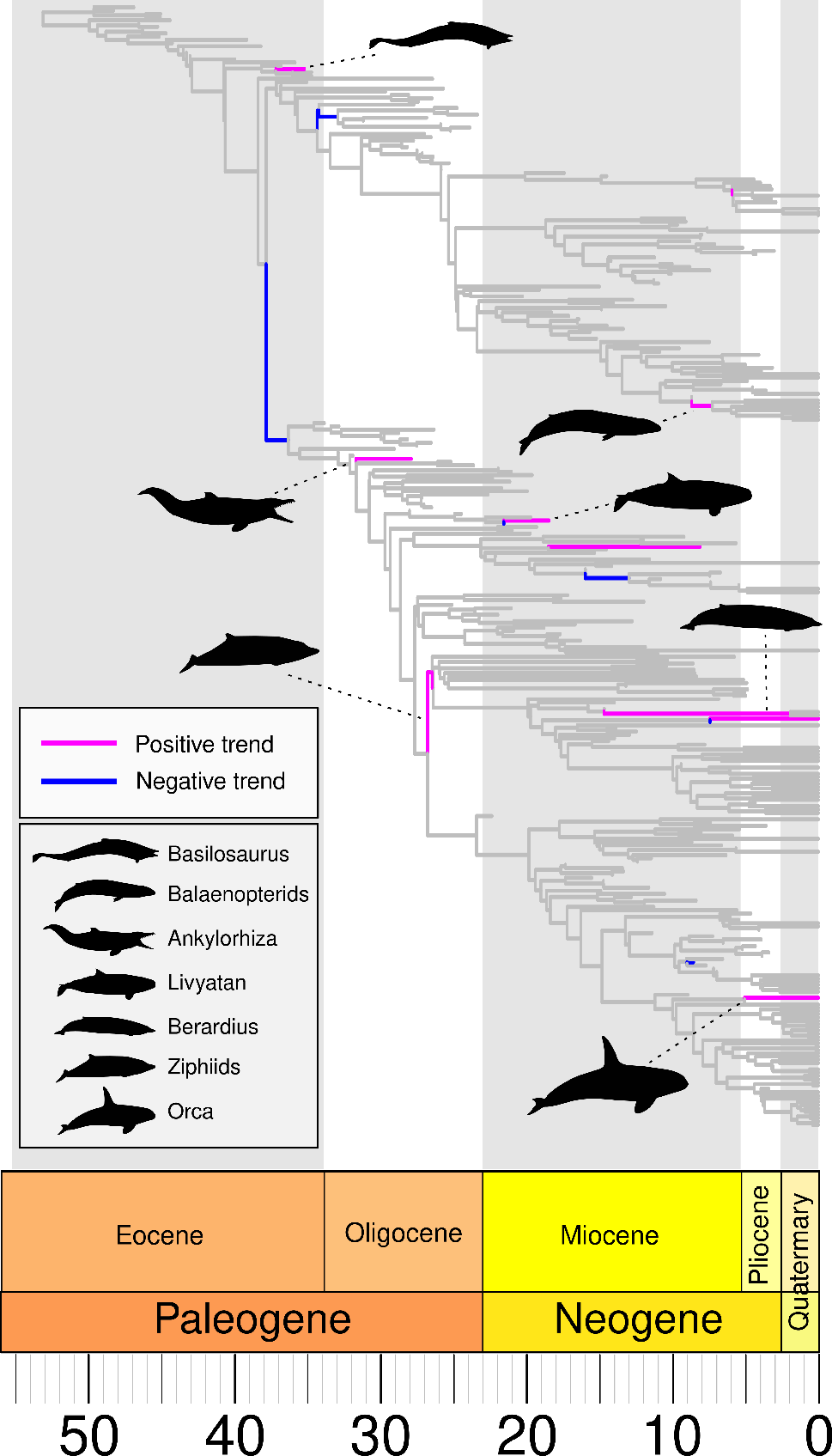
Evolutionary trends in total length evolution of living and extinct cetaceans. Phylogeny of cetaceans showing the branches with trends towards increasing or decreasing total length detected by the Fabric model^7^. Magenta lines represent branches with evidence of positive trends in average total length; blue lines represent branches with evidence of negative trends in average total length. Silhouettes indicate the main groups for which significant trends were found.

## Discussion

The evolution of morphological diversity demands explanation. Models of gradual evolution predict a simple increase in phenotypic variance through time, with more closely-related lineages exhibiting greater similarity to one another, on average, than they do to more distantly-related lineages^36, 37^. Where selection acts to drive lineages towards distinct phenotypes, it should mould and shape the distribution of phenotypic variation within and across clades, yielding patterns that deviate from the expectations of simple gradualistic models^5, 38^. Characterising these ‘ghosts of selection past’ is a central challenge of comparative evolutionary biology. Our analyses suggest fundamental limits in their ability to detect some evolutionary patterns and processes, and that a diversity of approaches are required to characterise these complex hierarchical patterns of phenotypic adaptation.

Comparative biologists increasingly model phenotypic macroevolutionary landscapes using the mathematics of the Ornstein-Uhlenbeck process, which connects microevolutionary processes^5^ to patterns of variance and covariance over phylogeny^39^. In this framework, peaks on the landscape act as phenotypic attractors while selection strength determines the steepness of the peaks. However, these models are data-hungry, and phylogenies with massive numbers of tips are required for accurate parameter estimation^40^. This, in turn, can result in a coarse, global view of the adaptive landscape, such as the identification of a single shift of large effect associated with the colonisation of the marine realm, or a shift to much smaller mean size associated with increased manouverability in piscivorous delphinidan odontocetes. What is missed, then, are more localised changes in the topography of the adaptive landscape, which may be particularly difficult to detect when different lineages experience change in different ways^4^.

These hidden dynamics are evident as multiple trends in body size evolution that appear on more local scales than the global peak shifts identified by our `bayou` analyses. In some cases, they occur along branches leading to individual species that can be characterised as possessing similar ecologies, such as trends towards larger size along branches leading to the macropredators (see also^41^). In other cases, trends are clade-dependent, such as a recent trend towards larger sizes in balaenopterids and, to a lesser extent, balaenids, associated with bulk feeding on dense prey patches^42^, and ziphiids, associated with deep diving behaviors^43^. Trends towards smaller sizes associated with the origin of odontocetes and kogiids may reflect similar responses to foraging on small, mobile prey, relative to ancestral states^44^.

The incongruence between a low number of global shifts and multiple conspicuous, more taxonomically-restricted local body size trends suggests a much more dynamic adaptive landscape than our analyses have found. How could this be the case? Body size is a complex and multifaceted trait, interconnected with almost every aspect of a species’ biology^45, 46^. This, in turn, means that body size is under multiple selection pressures at any given time, each one with its own adaptive mean, resulting in a locally rugged adaptive landscape (i.e. the principle of frustration *sensu*^47^). We argue that the close proximity of these long-term means facilitates movement between peaks on the adaptive landscape in relatively small increments that are not identified as discrete shifts in our `bayou` analyses but which appear as trends in the analysis using the Fabric model^7^. Therefore, we describe the adaptive landscape of body size in cetaceans over macroevolutionary timescales by comparing it to an ocean of turbulent waters; the few shifts we identified are analogous to the large waves, the smaller peaks that represent unique local conditions of adaptation^4^ or are ephemeral in nature, more akin to ripples on the surface of the ocean that change rapidly across both space and time, rather than to the more stable topography of terrestrial landscapes. These dynamics likely further increase the difficulty of peaks being detected as discrete shifts in comparative analyses, highlighting the need to use multiple approaches (incorporating fossils,^48^) to fully characterise adaptive landscapes in deep time, especially in traits as labile and integrated as body size.

## Methods

### Total length data

We collated total length (m) for living species primarily from stranding records in the literature (n = 1,327 individuals), and the strandings database at the Natural History Museum London (n = 1,659 individuals). If no other data were available we used data from aboriginal subsistence harvests (n = 80 individuals). We only included values within the range of adult lengths for each species, and, where possible, we recorded a minimum of 10 total length values for each species. This resulted in a total length dataset of 2,986 individuals of living species. We collated data on total lengths of fossils from the literature; some were true total length measurements, others were based on estimates (a total of n = 273). 87 species in our phylogeny (see below), two living and 85 fossil, had no available total length data. To estimate total length for these taxa we collated data for nine proxy measurements^16^ from 913 adult specimens (451 extinct and 462 living) from the published literature, or by measuring specimens using either Mitutoyo Absolute Digimatic CD-8"AX calipers, large calipers, or tape measures depending on the size of the feature. The proxy measurements were (1) width of antorbital notches, (2) bizygomatic width, (3) exoccipital width,(4) occipital condyle breadth, (5) condylobasal length, (6) maximum narial fossa width, (7) maximum nasal width, (8) atlas articular facet width, and (9) humerus length. We obtained a minimum of two proxy measurements from each specimen, as few fossils possessed all features. Prior to the analyses we removed species with uncertain taxonomy and those not present in the phylogeny.

### Phylogeny

We used the “Safe” analysis phylogeny from^21^ and removed all taxa without any total length proxy measurements from the phylogeny leaving 345 taxa. 60 of these remaining taxa were singletons, forming terminals with branch lengths of zero. Because our analyses required branch lengths, we added 0.001 MY to all branches to remove the zero-length branches. To ensure this did not substantially influence our results we repeated all downstream analyses (and sensitivity analyses; see Supplementary Information) removing the species with zero length branches from the phylogeny (n = 285 taxa; see Supplementary Information; Figures S26-S30).

### Total length data imputation

To impute total lengths for the 87 species without available total length data, we used a phylogenetically-informed imputation approach. These methods use the eigenvectors of the phylogenetic distance matrix to estimate trait values for species without the focal variable^49^,^50^ while accounting for intraspecific variation^49^,^50^. We used the phylogenetic imputation algorithm in the **R** package *Rphylopars*^50^, using a Brownian Motion model for trait evolution, and including specimens with at least one proxy measurement to incorporate intraspecific variation. All measurements were log_10_-transformed prior to the imputation. We also performed several sensitivity analyses to assess the robustness of our imputed total length data (see Supplementary Information). The final total length dataset contained 345 taxa; 89 living and 256 fossil.

### Exploring tempo and mode in body size evolution

We performed disparity through time analyses^51^ using the dtt function in the **R** package geiger^52^, with 10,000 simulated datasets used to generate a null, constant-rates distribution. The centre of gravity was computed as the weighted sum of the product of standardised average subclade disparity and relative node age, with significance determined as the proportion of simulated centres of gravity that were lower than the empirical value^53^.

We used *bayou*^6^ to detect evolutionary shifts in total length evolution across cetaceans. *bayou* uses reversible-jump MCMC algorithms to fit multi-peak OU models of trait evolution, identifying the number, magnitude and location of shifts in *θ*, the evolutionary regime mean of total length. We used a cumulative Poisson distribution as the prior distribution for the number of shifts, setting the prior on the expected number of shifts (*λ*) as 2.5% of the total number of branches in the corresponding tree, i.e. *λ* = 15. We allowed an unlimited number of shifts to occur on each branch, while setting the prior probability for each branch to contain a shift to be proportional to the branch’s length. For all other parameters, we used the recommended default distributions found in (https://github.com/uyedaj/bayou/blob/master/tutorial.md). We ran the MCMC chains for 1 million generations, sampling every 1,000 generations, yielding a posterior sample of 700 after discarding the first 30% as burn-in. We assessed the convergence of each chain using Gelman and Rubin’s R statistic, and confirmed that all parameters had an effective sample size greater than 200.

To assess the relative importance of the shifts we followed Uyeda *et al*.’s^6^ criteria: we selected shifts that had posterior probabilities greater than 0.1 (corresponding to approximately an eightfold increase in the posterior probability compared to the prior probability), with only three of these shifts leading to singleton taxa. For the selected shifts, we calculated the posterior-to-prior ratio as a measure of how much information our data is providing on the location and magnitude of changes in evolutionary regime. All `bayou` analyses were performed in R^54^.

### Sensitivity analyses

To assess the effect of prior parameterisation on the *bayou* results, we performed multiple sensitivity analyses using different values for the average expected shifts in the prior distribution (see Supplementary Information; Figures S5-S8). We also repeated all the *bayou* analyses above using five additional tree topologies (see Supplementary Information; Figures S9-S25), one excluding Archaeoceti (*No Archaeoceti*; n = 322), one including only fossil taxa (*No Extant*; n = 256), one including only living taxa (*Extant*; n = 89), one for Mysticeti only (*Baleen*; n = 105), and one for Odontoceti only (*Toothed*; n = 217), and for each of these topologies excluding taxa on zero length branches (*No Archaeoceti* (n = 267), *No Extant* (n = 207), *Baleen* (n = 230 83), *Toothed* (n = 184). Supplementary Information; Figures S31-S44. Note that the *Extant* tree does not contain any taxa with zero-length branches).

### Evaluating per-branch trends in body size evolution

We used the `Fabric` model^7^ as implemented in `BayesTraits` v4.0^55^ to evaluate the presence, location, and strength of linear trends in cetacean total length evolution. This model assumes that evolution follows a random walk in which the variance parameter (= evolutionary rate) can vary across branches of the phylogeny. However, it also uses a reversible jump MCMC to sample trend parameters along branches that result in biases towards larger (+ve) or smaller (-ve) trait values, accepting these if they increase the probability of the model.

We ran three independent chains with 5 million steps of burn-in and 20 million steps of sampling. After visually checking that all three chains converged on the same target distribution and that effective sample sizes were larger than 200, we used the Fabric post-processing software^7^ to summarise the output of each chain, ultimately retaining all branch trends that were significant in all three chains. The processed output file can be downloaded from the NHM Data Portal^56^.

## Acknowledgements

We thank Roberto Portela Miguez and Richard Sabin for collections access. We also thank Josef Uyeda, Fábio Machado and 348 Daniel Caetano for their helpful feedback and discussions about bayou results. This work was funded by Leverhulme Trust 349 Research Project grant (RPG-2019-323).

## Author contributions statement

All authors conceived the ideas. T.P. assembled the dataset. G.B. performed the analyses. All authors drafted and reviewed the manuscript.

## Data accessibility statement

Data are available from the NHM Data Portal^56^ and code is available from GitHub (https://github.com/gburin/bodysize-355evolution-cetacea) and will be deposited at Zenodo upon acceptance.

