## Supplementary Information for "Turbulent adaptive landscape shaped size evolution in modern ocean giants"

### Supplementary Information from: *Turbulent adaptive landscape shaped size evolution in modern ocean giants*

**these authors contributed equally*

3.5 Supplementary Results: Removing taxa with zero-length branches 38

**4 Supplementary References** **63**

### 1 Comparison of imputed and empirical total length data

#### 1.1 Supplementary Methods

To assess the quality of total length estimates using the phylogenetic imputation approach, we generated datasets with different percentages of missing total length data for species with available empirical total length data. We then compared the values estimated by phylogenetic imputation with the empirical values to assess the accuracy of the approach.

To create the test datasets we randomly removed data for 10% (n = 25), 20% (n = 51), 50% (n = 129), of all species with existing total length data, leaving 233, 207 and 129 species respectively. Note that we removed total length data for all specimens belonging to these species, so in some cases the percentage of missing total length data *records* is larger than the percentage of missing *species*. We also generated a dataset with empirical total length data from extant species only (n = 89).

For all four datasets, we replicated the same protocol as in the main analysis. Total length values and proxy measurements were $\log_{10}$-transformed, and we used the phylogenetic imputation algorithm in the **R** package *Rphylopars* ([1]), using a Brownian Motion model for trait evolution, and including specimens with at least one proxy measurement to incorporate intraspecific variation.

We assessed the accuracy of estimated values by plotting the imputed values against the empirical average values for the species whose total length data was removed from each dataset (Figures S1 to S4).

We also calculated the Normalized Root Mean Squared Error for the main dataset (NRMSE;[2]), which represents the mean squared errors for the imputed values standardized by the range of empirical values (equation 1).

$$NRMSE = \frac{\sqrt{\frac{\sum(X_i^* - X_i)^2}{n}}}{max(X) - min(X)} \tag{1}$$

Where $X$ represents the variable "total length", $X_i^*$ is the imputed value of total length for species $i$, $X_i$ is the empirical value of total length for species $i$, and $n$ is the number of taxa. Mean squared errors were calculated for estimates using all possible combinations of proxy measurements used for the imputation (ranging from using just one measurement up to all nine).

#### 1.2 Supplementary Results

Figures S1 to S4 show the association between imputed and empirical average total length values. Imputed total length data were strongly correlated with the empirical values (Figures S1 to S4), regardless of the amount of information used for the estimation of missing values.

Normalized Root Mean Square Errors (NRMSE) values for each dataset with different numbers of proxy variables removed are shown in Table S1. NRMSE was low, and within acceptable values found in other papers (3–5).

Together these results suggest that our phylogenetically-informed imputations provide robust estimates of total length.

Table S1: *Normalized Root Mean Square Errors (NRMSE) values calculated for each manipulated dataset. Each column represents the NRMSE value calculated over all possible combinations of measurements after removing a given number of measurements prior to the imputation. Mean* $\pm$ *standard deviation of combined NRMSE values* $= 0.2091 \pm 0.0959$.

| Removed variables | 1 | 2 | 3 | 4 | 5 | 6 | 7 | 8 |
|---|---|---|---|---|---|---|---|---|
| NRMSE | 0.087 | 0.176 | 0.259 | 0.321 | 0.322 | 0.259 | 0.167 | 0.082 |

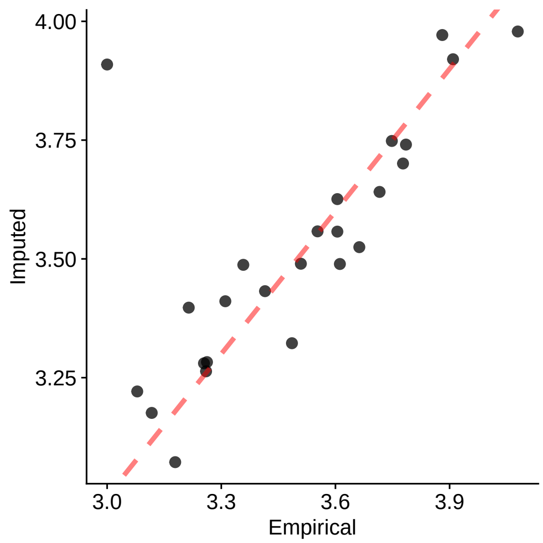

Figure S1: Relationship between imputed and empirical average total length values after randomly dropping 10% of species with total length data. Each point represents a species with total length data removed prior to the imputation (n = 25). The red dotted line is the 1:1 line.

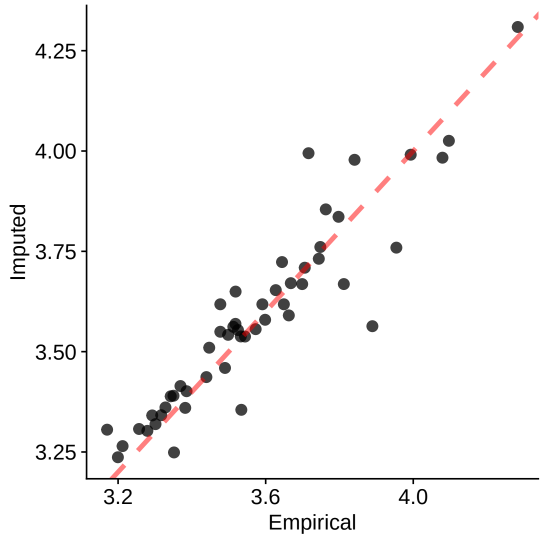

Figure S2: Relationship between imputed and empirical average total length values after randomly dropping 20% of species with total length data. Each point represents a species with total length data removed prior to the imputation (n = 51). The red dotted line is the 1:1 line.

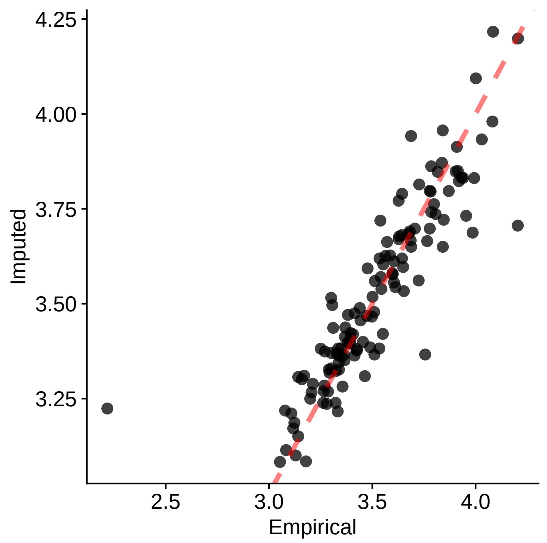

Figure S3: Relationship between imputed and empirical average total length values after randomly dropping 50% of species with total length data. Each point represents a species with total length data removed prior to the imputation (n = 129). The red dotted line is the 1:1 line.

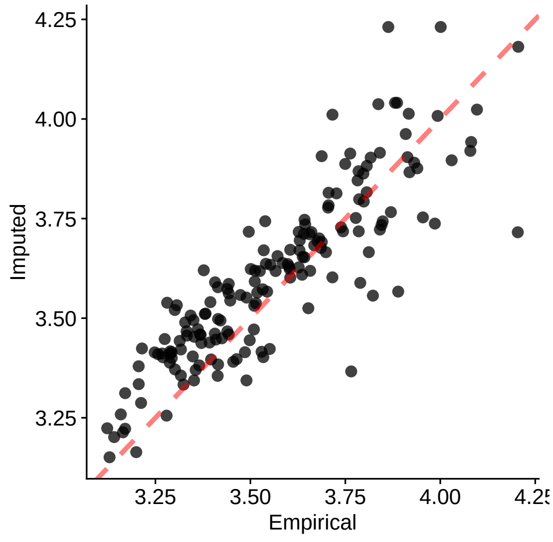

Figure S4: Relationship between imputed and empirical average total length values after removing all fossil species with total length data. Each point represents a species with total length data removed prior to the imputation (n = 170). The red dotted line is the 1:1 line.

#### 2 Disparity through time

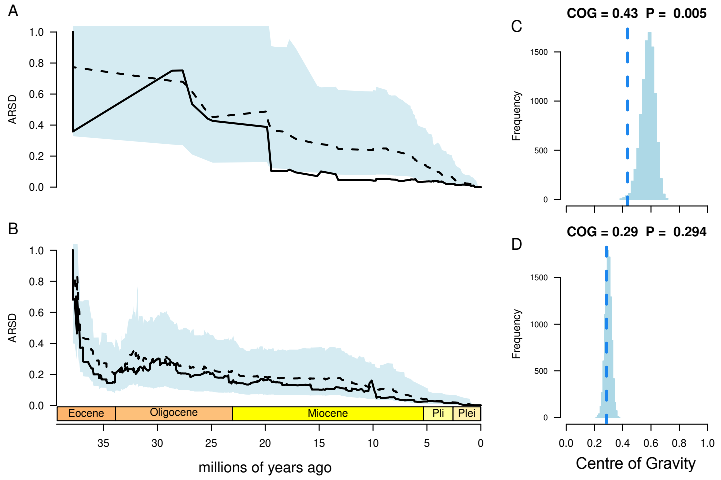

Figure S5: The disparity through time profile for extant cetaceans (A) is consistent with an adaptive radiation, with the average relative subclade disparity (ARSD) dropping substantially below constant rates expectation in the Early Miocene. This is reflected in significantly low centre of gravity (C). Adding fossil data (here, crown neocetes B, though similar results obtain for total group Cetacea) results in a subclade disparity profile that is more consistent with, and not signficantly different from (D) constant rates expectation.

#### 2.1 Supplementary Results: Root Theta

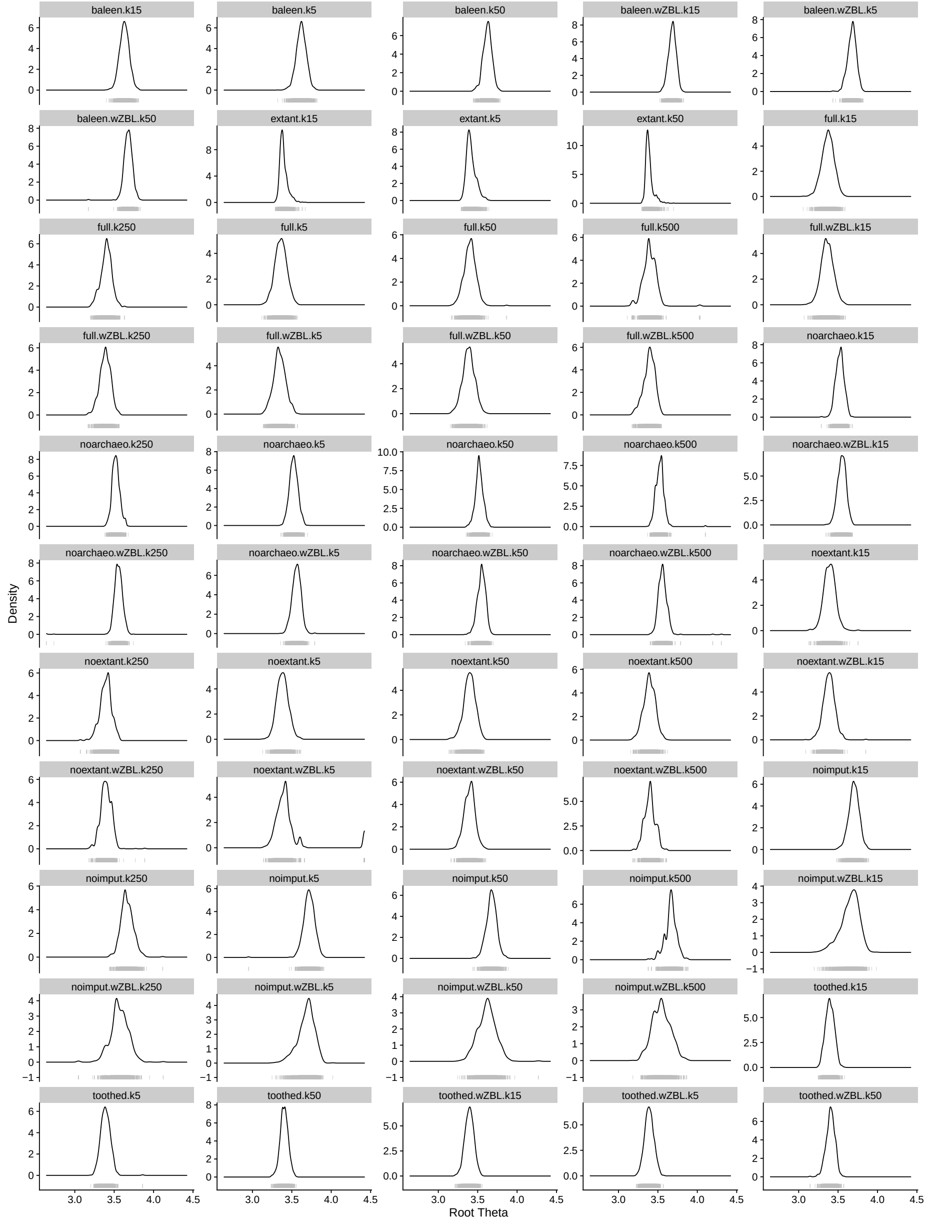

Figure S6: Posterior distribution of estimated $\theta$ values at the root for each scenario of the `bayou` analyses. In each panel, the grey dashes below the x axis represent the raw values of the posterior, that are summarised as density curves in the plots.

### 3 *bayou* sensitivity analyses

#### 3.1 Supplementary Methods

##### 3.1.1 Phylogenies and priors

To test the sensitivity of our results to both tree topology and the prior parameter for expected number of shifts ($\lambda$), we replicated the *bayou* protocol described in the main text for the original tree (*Full* tree) and six additional trees, with five different values of $\lambda$.

The six trees were as follows:

1. ***Full***. The most comprehensive phylogeny possible that included all species with total length data (n = 345). This is the the "Safe" analysis phylogeny from[6] with species without total length data (empirical or imputed) removed, and is the tree used for the analyses in the main text.

2. ***No Archaeoceti***. The *Full* tree after dropping all species that are classed as archaeocetes (n = 322).

3. ***No Extant***. The *Full* tree for fossil species only (n = 256).

4. ***No Imput***. The *Full* tree after dropping all species that lacked direct total length data (n = 258).

5. ***Extant***. The *Full* tree for living species only (n = 89). Note that this tree does not contain any taxa with zero-length branches (see below).

6. ***Baleen***. The *Full* tree for baleen whales (Mysticeti) only (n = 105).

7. ***Toothed***. The *Full* tree for toothed whales (Odontoceti) only (n = 217).

For each of these seven phylogenies in turn, we replicated the same *bayou* analysis pipeline described in the main text using five different values for the prior parameter on the expected number of shifts ($\lambda$), namely 5, 15, 50, 250 and 500 (the last two values were not used for the *Extant*, *Baleen* or *Toothed* trees because there were too few branches in those trees). Note that the prior on the expected number of shifts for the main analyses was set as 2.5% of the total number of branches in the corresponding tree, i.e. $\lambda = 15$. The results were evaluated the same way for all analyses; we considered only the shifts with posterior probability greater than 0.1 and also assessed the posterior-to-prior ratio for the selected shifts.

##### 3.1.2 Removing taxa with zero-length branches

As described in the main analyses, 60 of the taxa in the *Full* tree were singletons, forming terminals with branch lengths of zero. Because our analyses required branch lengths, we added 0.001 MY to all branches to remove the zero-length branches, and used these branch lengths in our main analyses. To ensure this did not substantially influence our results we repeated all our analyses removing the species with zero length branches from the phylogeny, leaving n = 285 taxa in the *Full* tree. We repeated this for the other four trees containing singletons: *No Archaeoceti* (n = 267), *No Extant* (n = 207), *No Imput* (n = 216), *Baleen* (n = 83), *Toothed* (n = 184). Note that the *Extant* tree does not contain any taxa with zero-length branches.

#### 3.2 Supplementary Results: Summary

Overall, our results using different trees and values of $\lambda$ are congruent with the main results (Figures S8 to S56).

Using all taxa, the shifts observed in the main scenario (*Full* tree, $\lambda = 15$, Figure 1) were retained, regardless of the tree and $\lambda$ value used, provided that the tree allowed for a given shift to exist (i.e. we cannot observe the shifts for archaeocetes in the analyses that exclude this group; Figures S8 to S32). Our analyses removing taxa with zero length branches were also consistent with the main results (Figures S33 to S56).

The results for different values of $\lambda$ highlight the sensitivity of the method to this parameter, with the average posterior number of shifts converging towards the average prior number, i.e. as $\lambda$ increases, more shifts are detected (leading to many more triangles in the plots; see Figures S8 to S56).

Although some of the scenarios we tested are very unlikely to represent the true evolutionary history of the group (such as the ones using 250 and 500 shifts; e.g. Figure S11), the shifts that group large numbers of lineages (such as shifts 1, 2 and 8 in Figure 1) are consistent across our sensitivity analyses regardless of the tree and prior parameters used (Figures S8 to S56). Lastly, the results from the analysis excluding species without direct total length data also show similar patterns to the main analyses, highlighting that is it unlikely that these species are somehow biasing our results. This provides confidence that these represent true shifts in mean total length regime ($\theta$).

#### 3.3 Supplementary Results: Phenogram

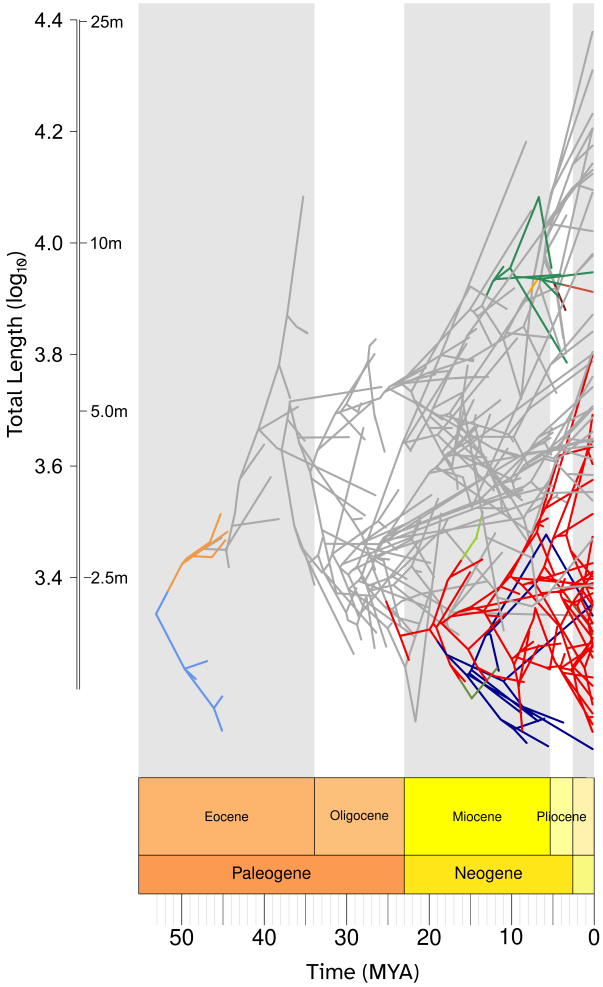

Figure S7: Phenogram of total length evolution in cetaceans. The tips, branches and nodes of the phylogeny are re-positioned according to the trait value, and are coloured according to the regime shifts in Figure 2). This figure shows that there is little overlap in total length between the major regimes, and that the trait space is homogeneously occupied, which highlights the flat adaptive landscape found for cetaceans.

#### 3.4 Supplementary Results: Phylogenies and priors

##### 3.4.1 *Full* tree

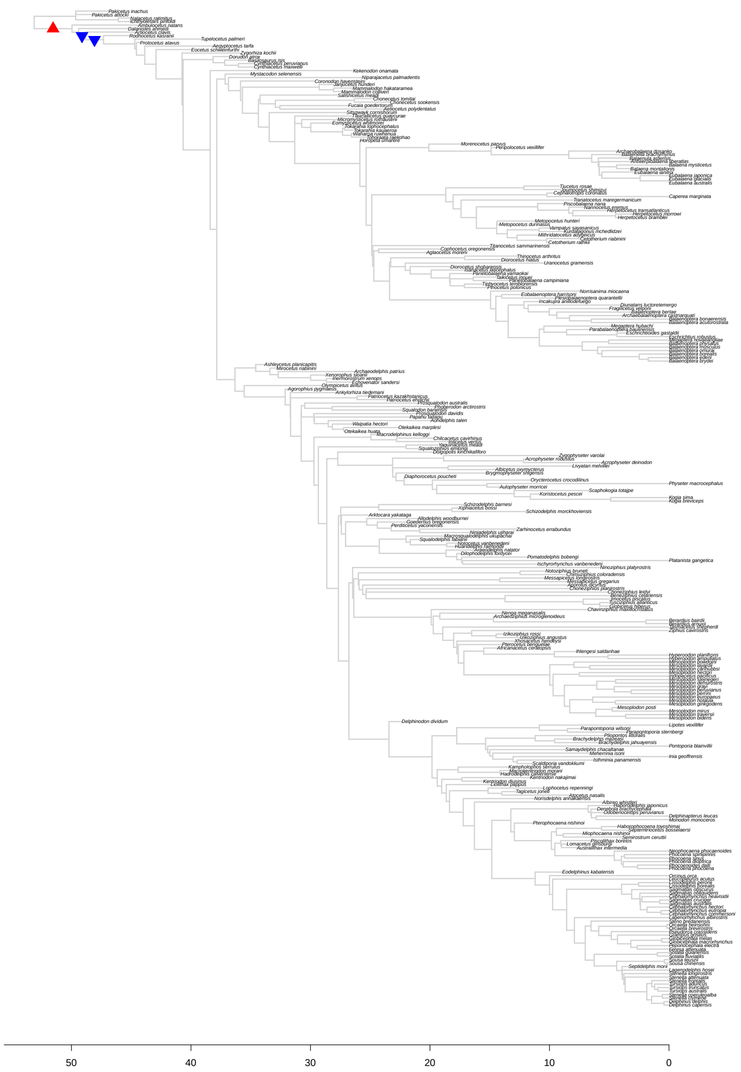

Figure S8: Results for *bayou* fit for the *Full* tree setting the average number of shifts in the prior distribution ($\lambda$) to 5. The triangles represent the position and direction of the shifts with posterior probability higher than 0.1, with upward triangles (in red) indicating increases in $\theta$, and downward triangles (in blue) indicating decreases in $\theta$.

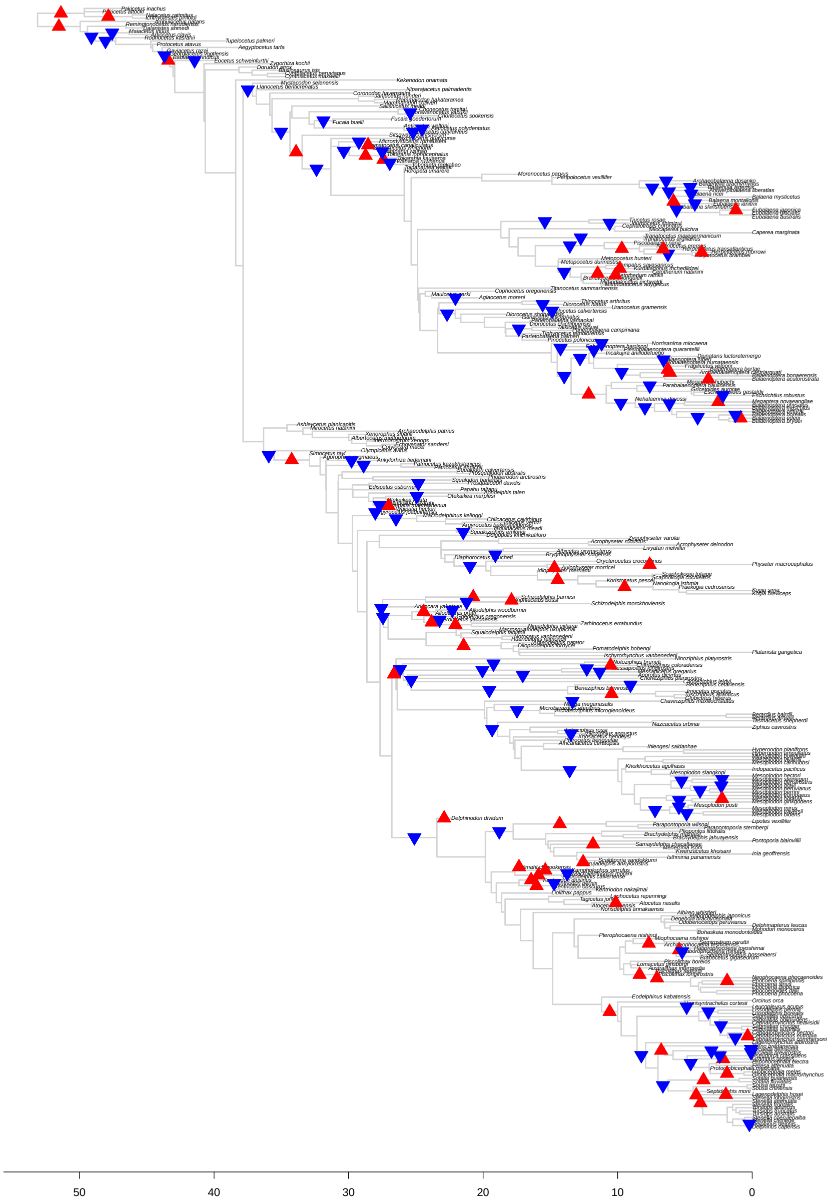

Figure S9: Results for *bayou* fit for the *Full* tree setting the average number of shifts in the prior distribution ($\lambda$) to 50. The triangles represent the position and direction of the shifts with posterior probability higher than 0.1, with upward triangles (in red) indicating increases in $\theta$, and downward triangles (in blue) indicating decreases in $\theta$.

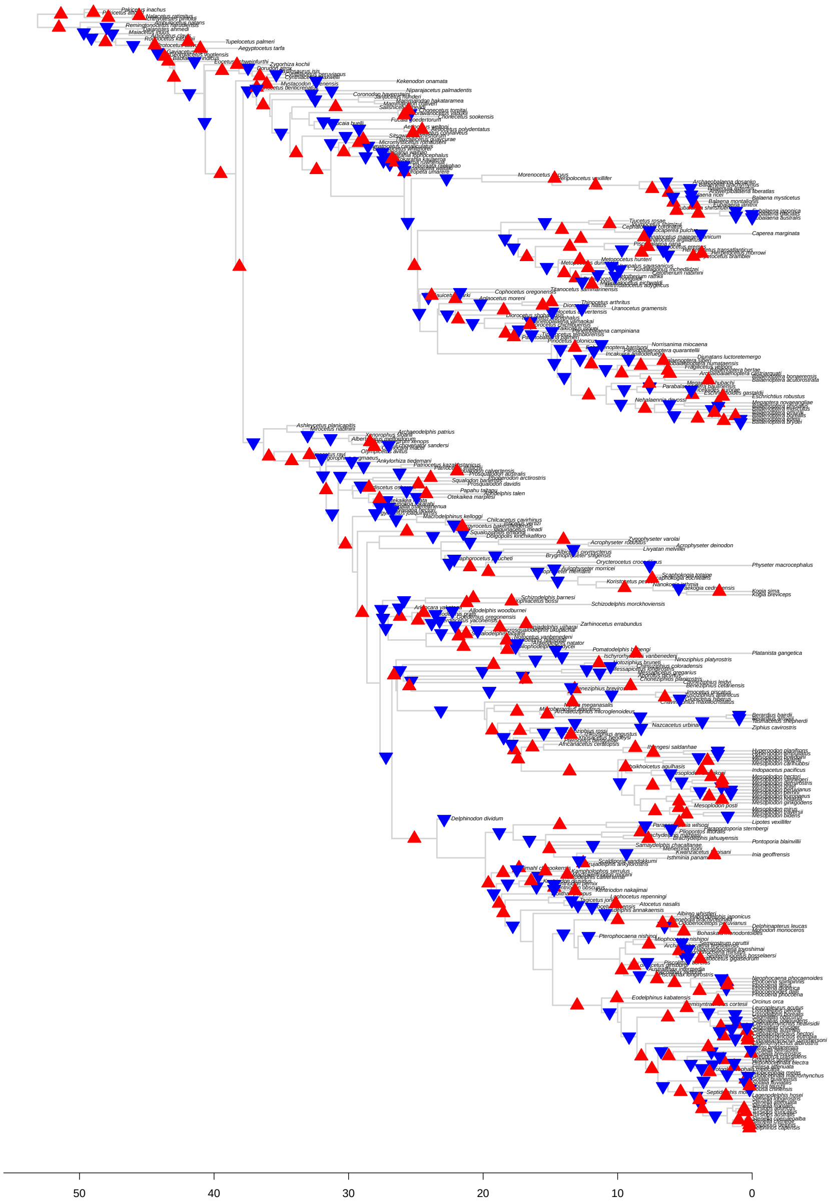

Figure S10: Results for *bayou* fit for the *Full* tree setting the average number of shifts in the prior distribution ($\lambda$) to 250. The triangles represent the position and direction of the shifts with posterior probability higher than 0.1, with upward triangles (in red) indicating increases in $\theta$, and downward triangles (in blue) indicating decreases in $\theta$.

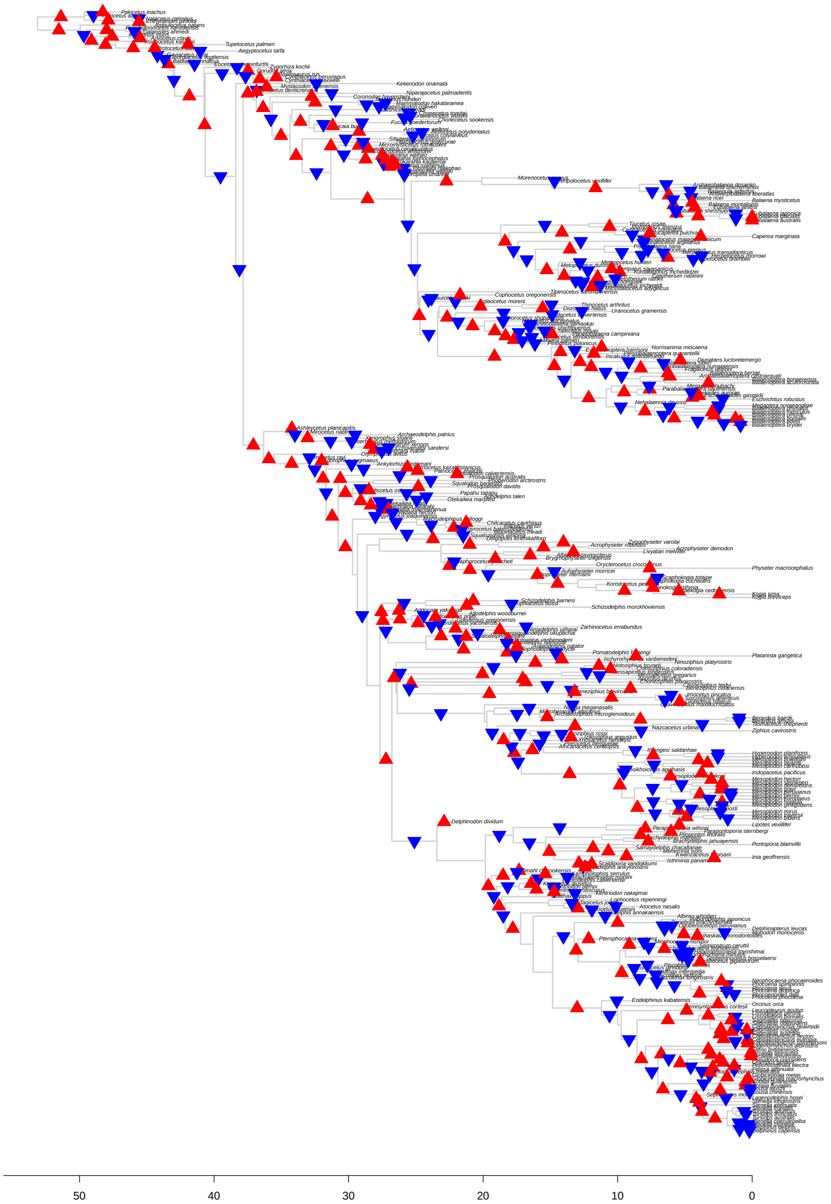

Figure S11: Results for *bayou* fit for the *Full* tree setting the average number of shifts in the prior distribution ($\lambda$) to 500. The triangles represent the position and direction of the shifts with posterior probability higher than 0.1, with upward triangles (in red) indicating increases in $\theta$, and downward triangles (in blue) indicating decreases in $\theta$.

##### 3.4.2 *No Archaeoceti* tree

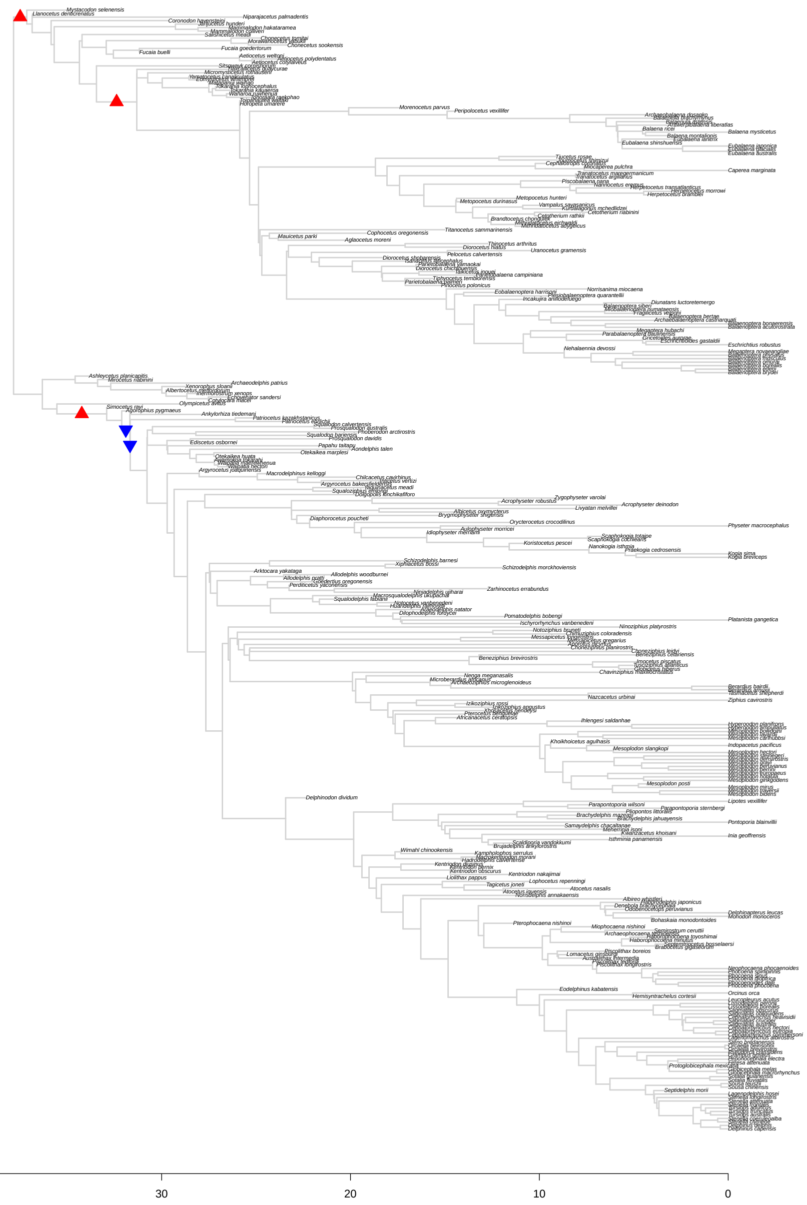

Figure S12: Results for *bayou* fit for the *No Archaeoceti* tree setting the average number of shifts in the prior distribution ($\lambda$) to 5. The triangles represent the position and direction of the shifts with posterior probability higher than 0.1, with upward triangles (in red) indicating increases in $\theta$, and downward triangles (in blue) indicating decreases in $\theta$.

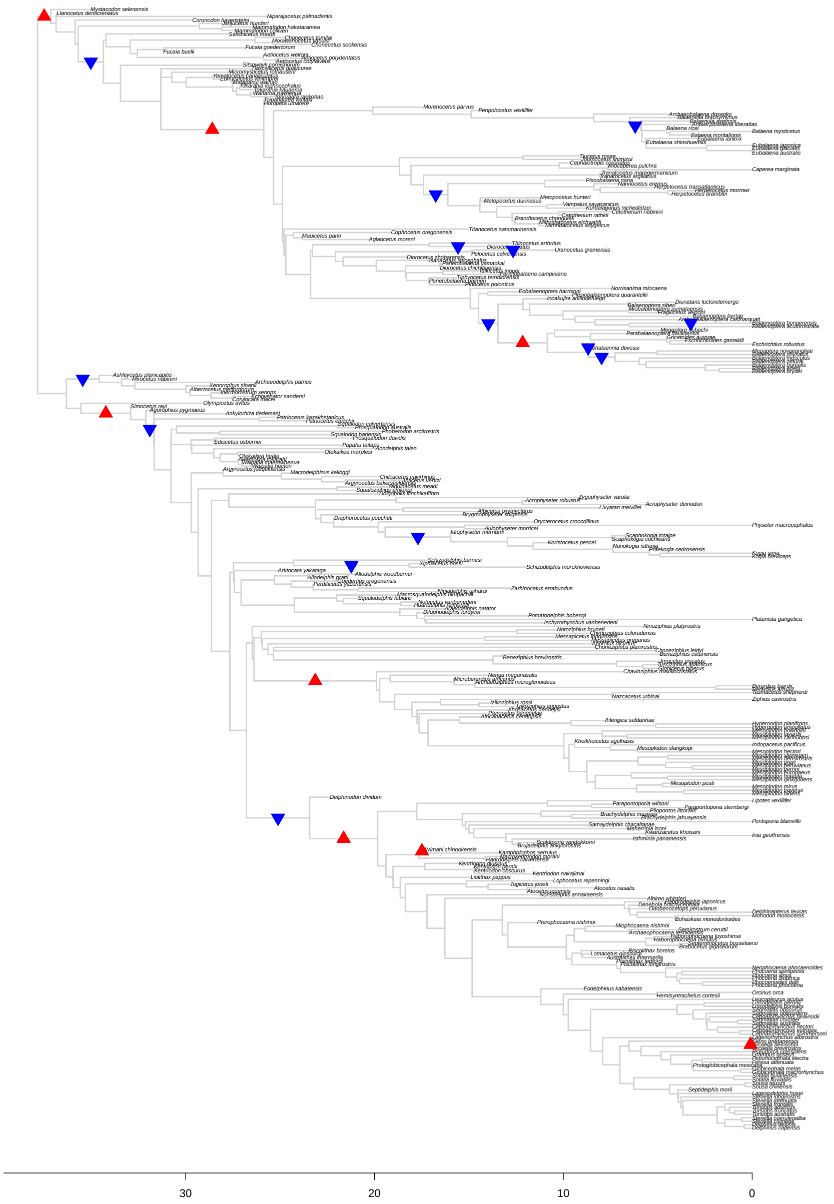

Figure S13: Results for *bayou* fit for the *No Archaeoceti* tree setting the average number of shifts in the prior distribution ($\lambda$) to 15. The triangles represent the position and direction of the shifts with posterior probability higher than 0.1, with upward triangles (in red) indicating increases in $\theta$, and downward triangles (in blue) indicating decreases in $\theta$.

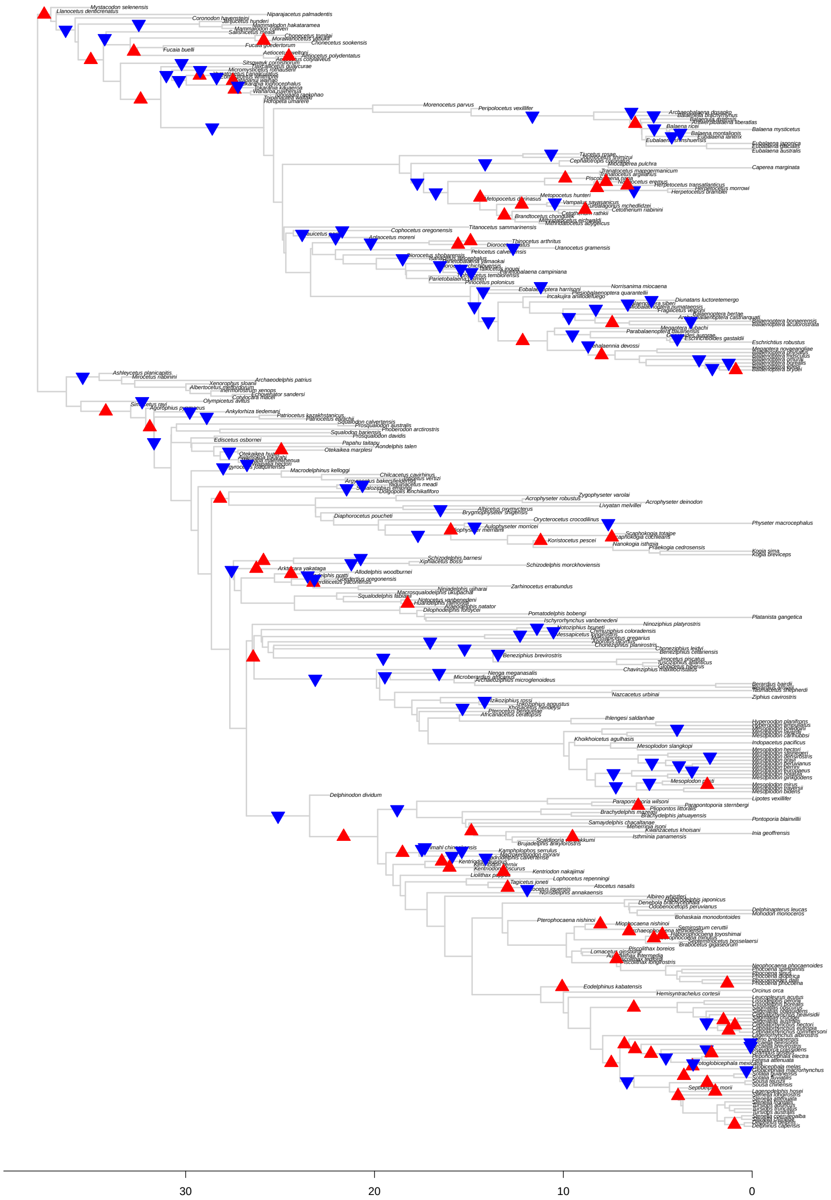

Figure S14: Results for *bayou* fit for the *No Archaeoceti* tree setting the average number of shifts in the prior distribution ($\lambda$) to 50. The triangles represent the position and direction of the shifts with posterior probability higher than 0.1, with upward triangles (in red) indicating increases in $\theta$, and downward triangles (in blue) indicating decreases in $\theta$.

##### 3.4.3 *No Extant* tree

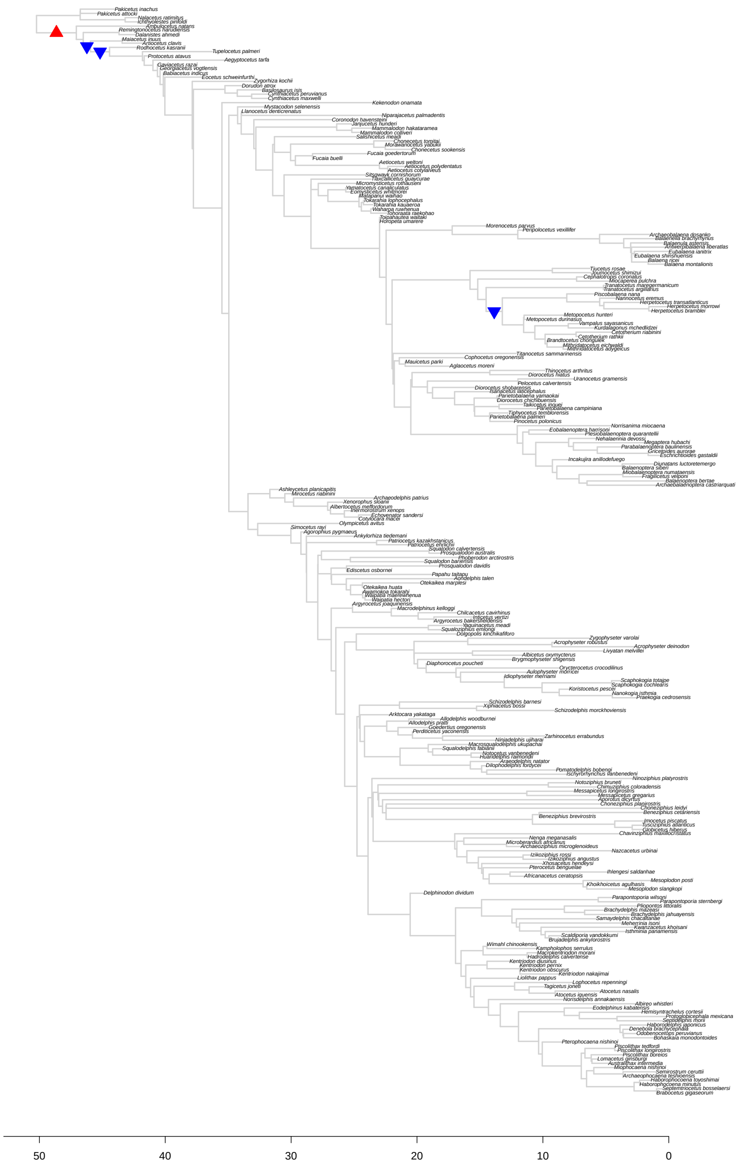

Figure S15: Results for *bayou* fit for the *No Extant* tree setting the average number of shifts in the prior distribution ($\lambda$) to 5. The triangles represent the position and direction of the shifts with posterior probability higher than 0.1, with upward triangles (in red) indicating increases in $\theta$, and downward triangles (in blue) indicating decreases in $\theta$.

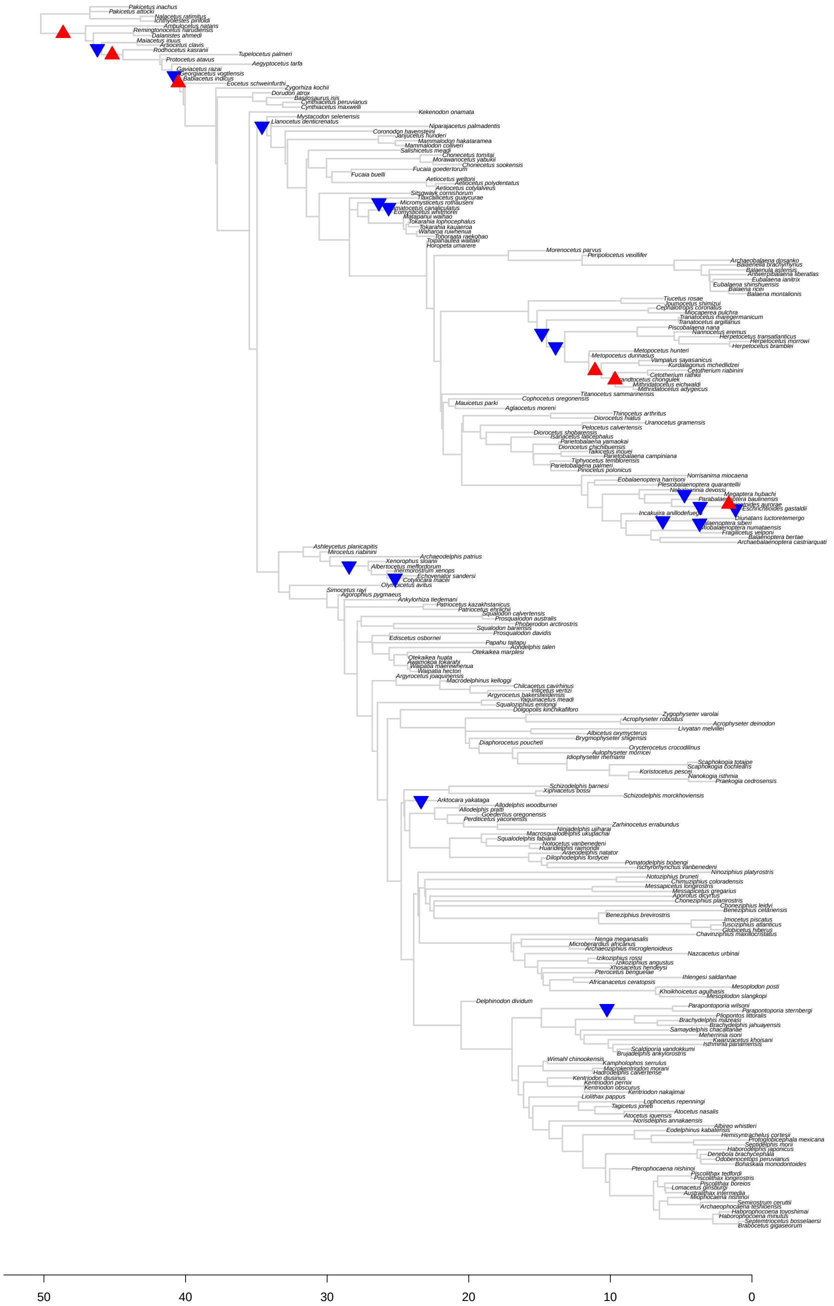

Figure S16: Results for *bayou* fit for the *No Extant* tree setting the average number of shifts in the prior distribution ($\lambda$) to 15. The triangles represent the position and direction of the shifts with posterior probability higher than 0.1, with upward triangles (in red) indicating increases in $\theta$, and downward triangles (in blue) indicating decreases in $\theta$.

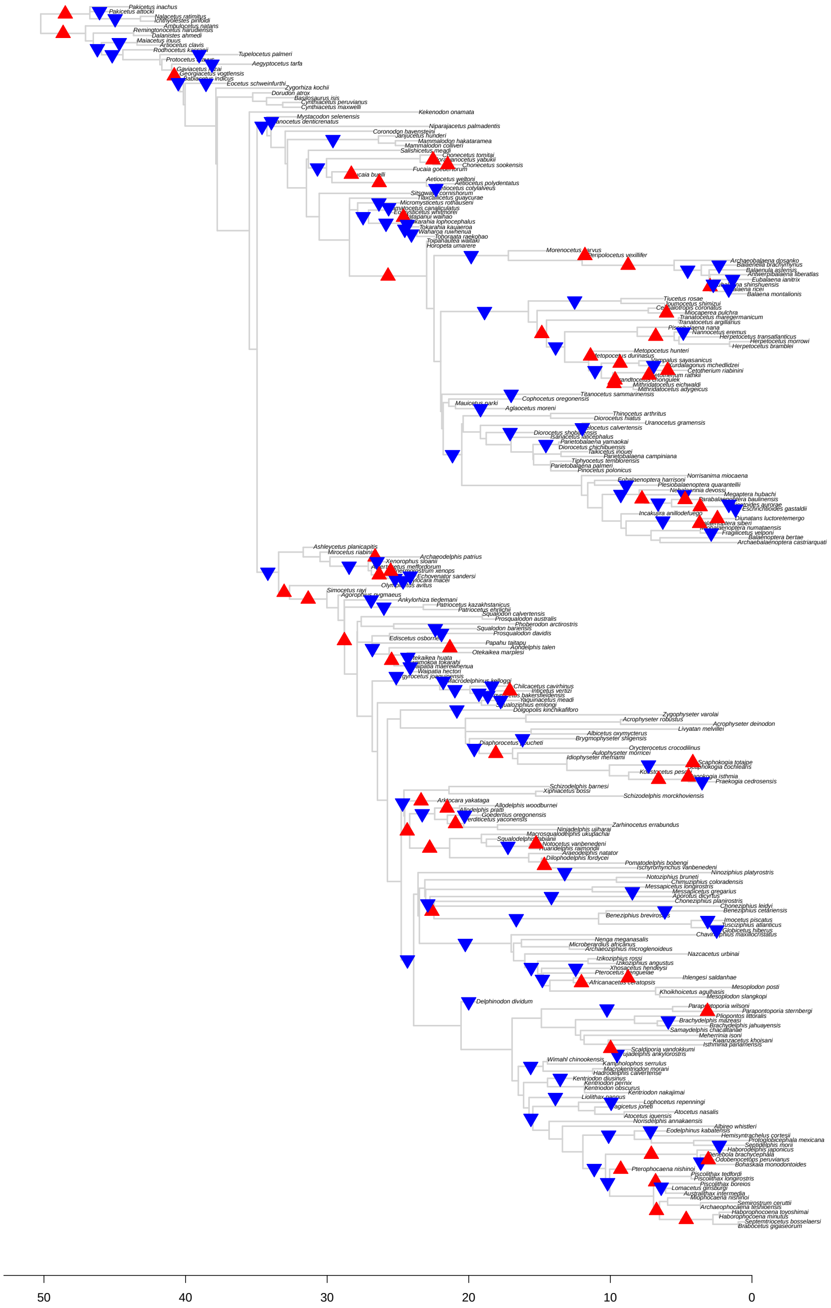

Figure S17: Results for *bayou* fit for the *No Extant* tree setting the average number of shifts in the prior distribution ($\lambda$) to 50. The triangles represent the position and direction of the shifts with posterior probability higher than 0.1, with upward triangles (in red) indicating increases in $\theta$, and downward triangles (in blue) indicating decreases in $\theta$.

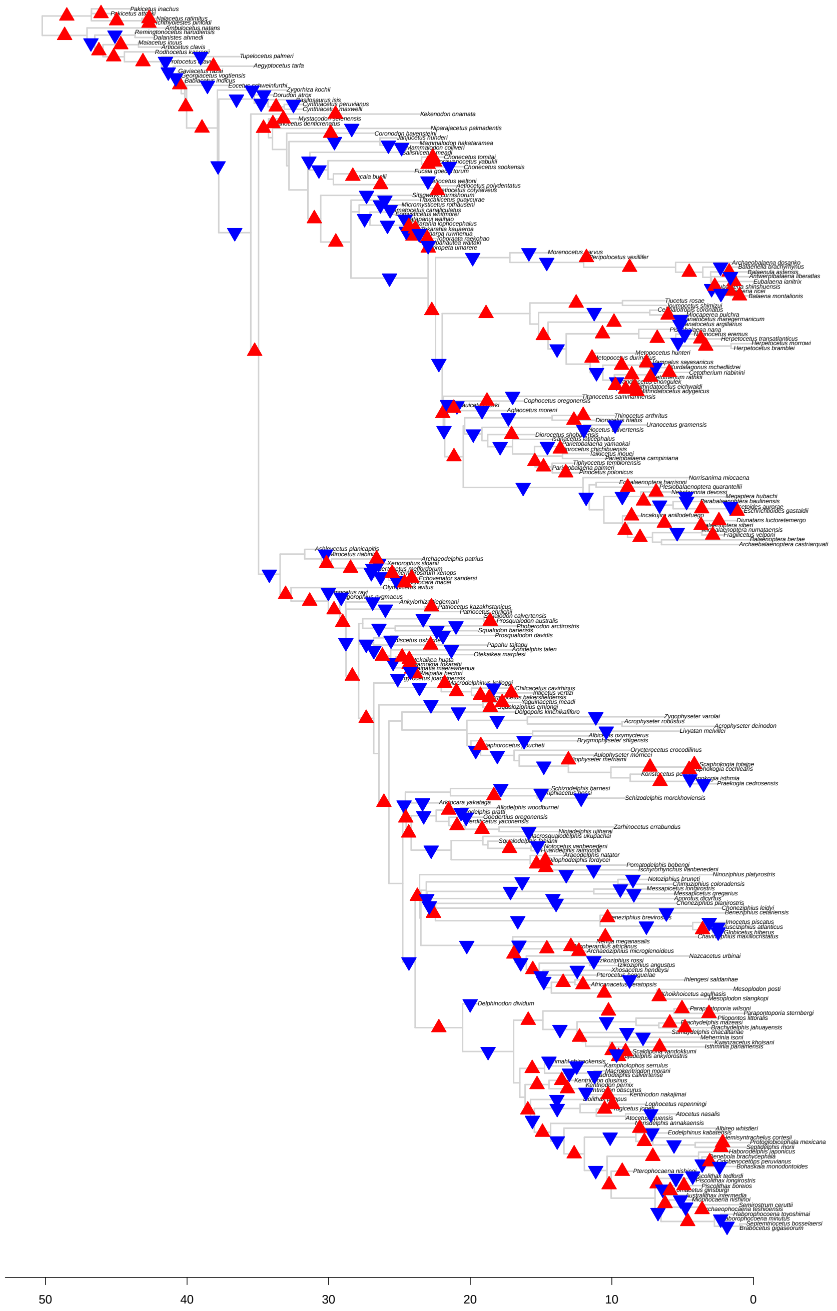

Figure S18: Results for *bayou* fit for the *No Extant* tree setting the average number of shifts in the prior distribution ($\lambda$) to 250. The triangles represent the position and direction of the shifts with posterior probability higher than 0.1, with upward triangles (in red) indicating increases in $\theta$, and downward triangles (in blue) indicating decreases in $\theta$.

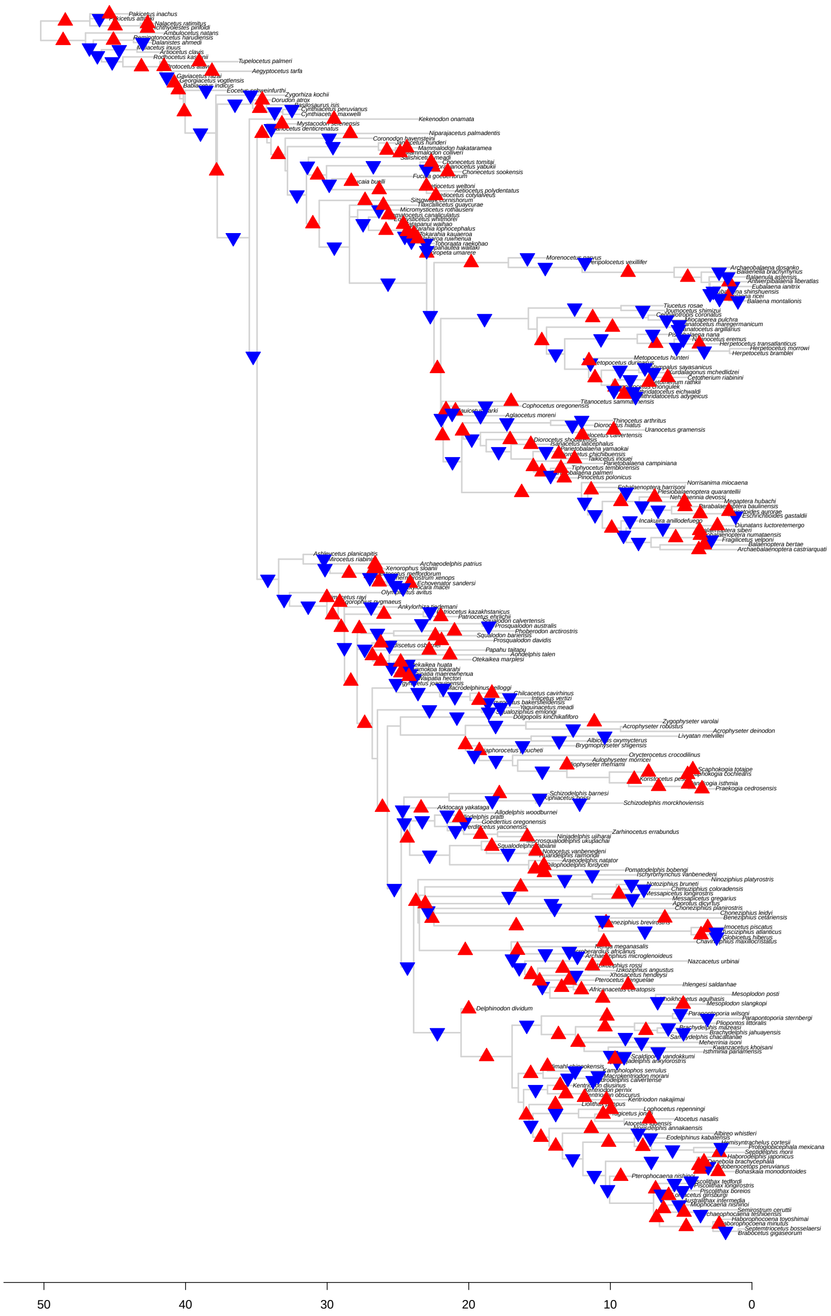

Figure S19: Results for *bayou* fit for the *No Extant* tree setting the average number of shifts in the prior distribution ($\lambda$) to 500. The triangles represent the position and direction of the shifts with posterior probability higher than 0.1, with upward triangles (in red) indicating increases in $\theta$, and downward triangles (in blue) indicating decreases in $\theta$.

##### 3.4.4 *No Imput* tree

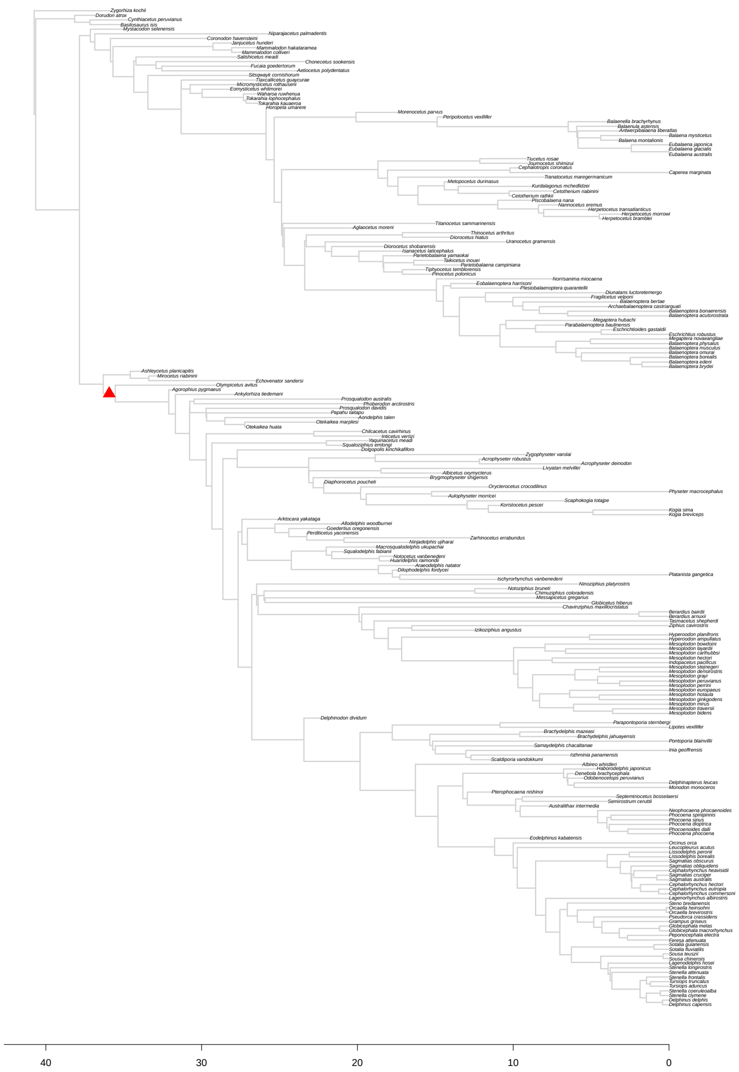

Figure S20: Results for *bayou* fit for the *No Imput* tree setting the average number of shifts in the prior distribution ($\lambda$) to 5. The triangles represent the position and direction of the shifts with posterior probability higher than 0.1, with upward triangles (in red) indicating increases in $\theta$, and downward triangles (in blue) indicating decreases in $\theta$.

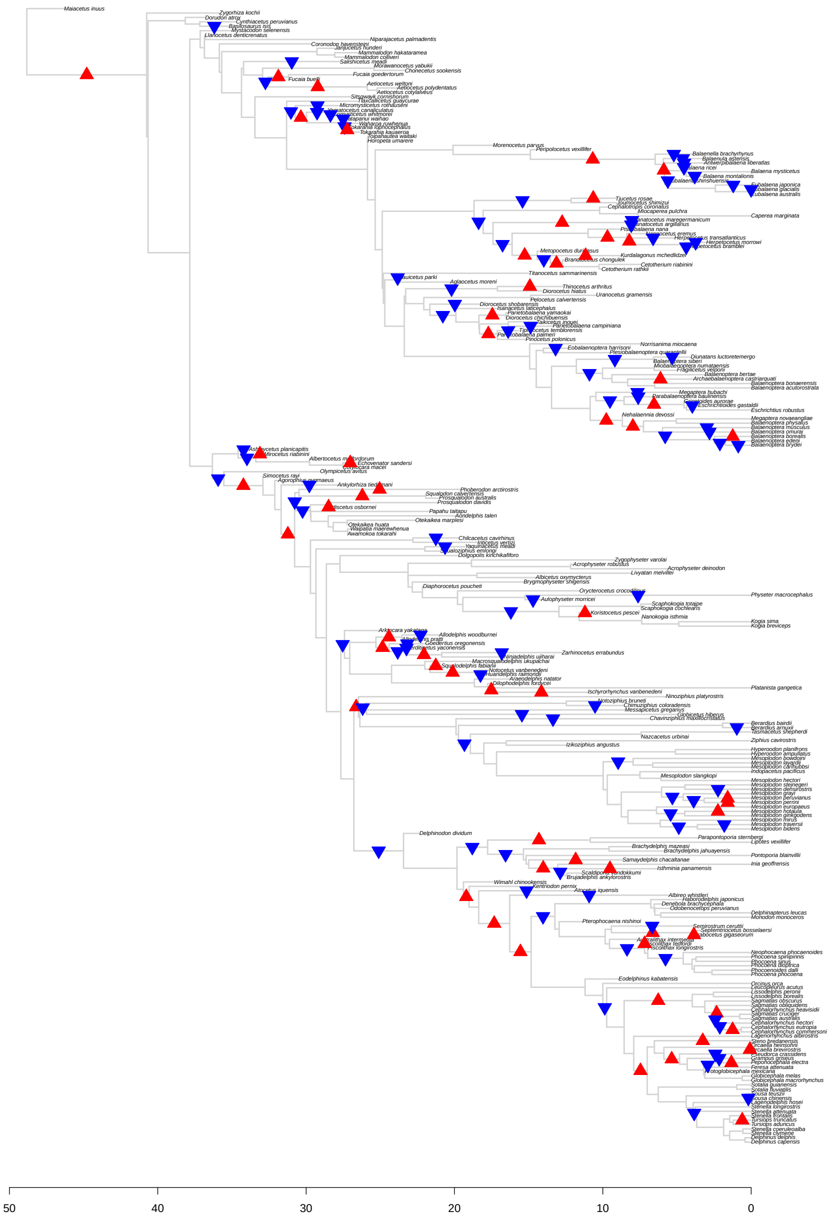

Figure S21: Results for *bayou* fit for the *No Imput* tree setting the average number of shifts in the prior distribution ($\lambda$) to 50. The triangles represent the position and direction of the shifts with posterior probability higher than 0.1, with upward triangles (in red) indicating increases in $\theta$, and downward triangles (in blue) indicating decreases in $\theta$.

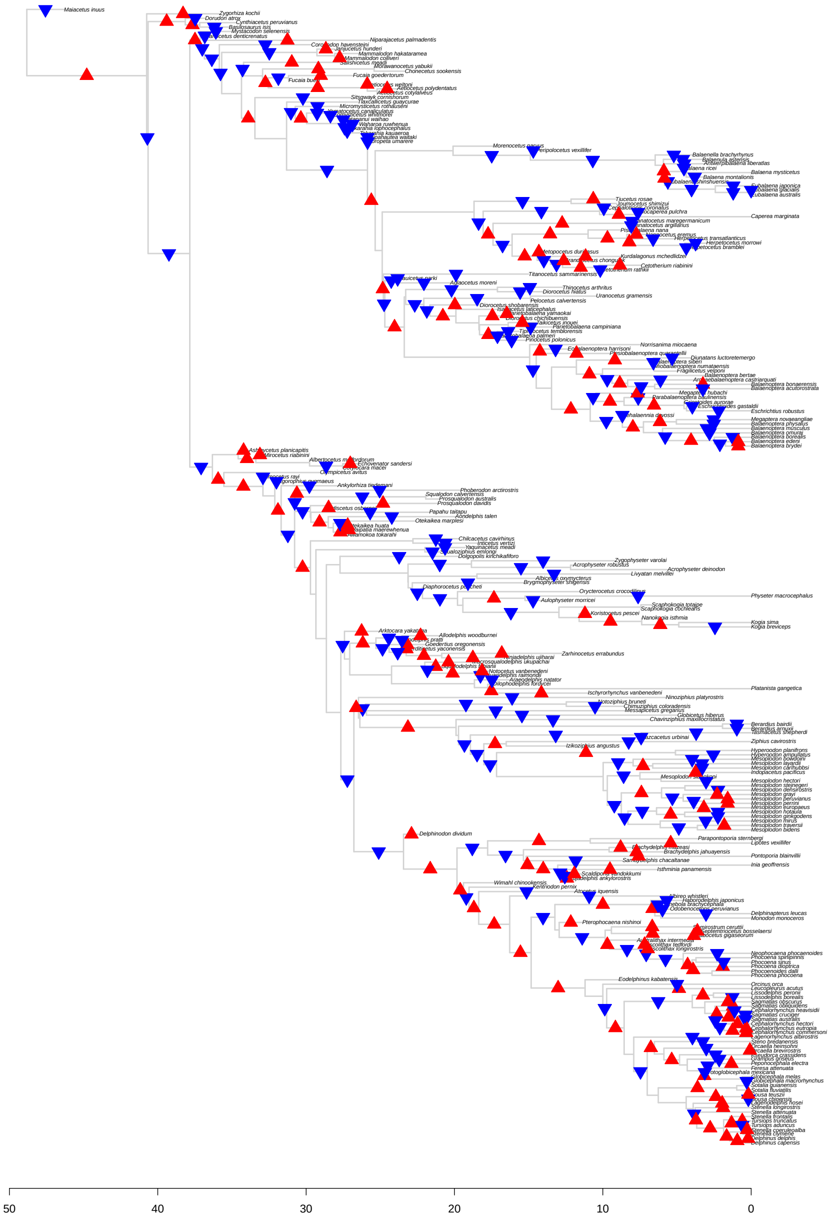

Figure S22: Results for *bayou* fit for the *No Imput* tree setting the average number of shifts in the prior distribution ($\lambda$) to 250. The triangles represent the position and direction of the shifts with posterior probability higher than 0.1, with upward triangles (in red) indicating increases in $\theta$, and downward triangles (in blue) indicating decreases in $\theta$.

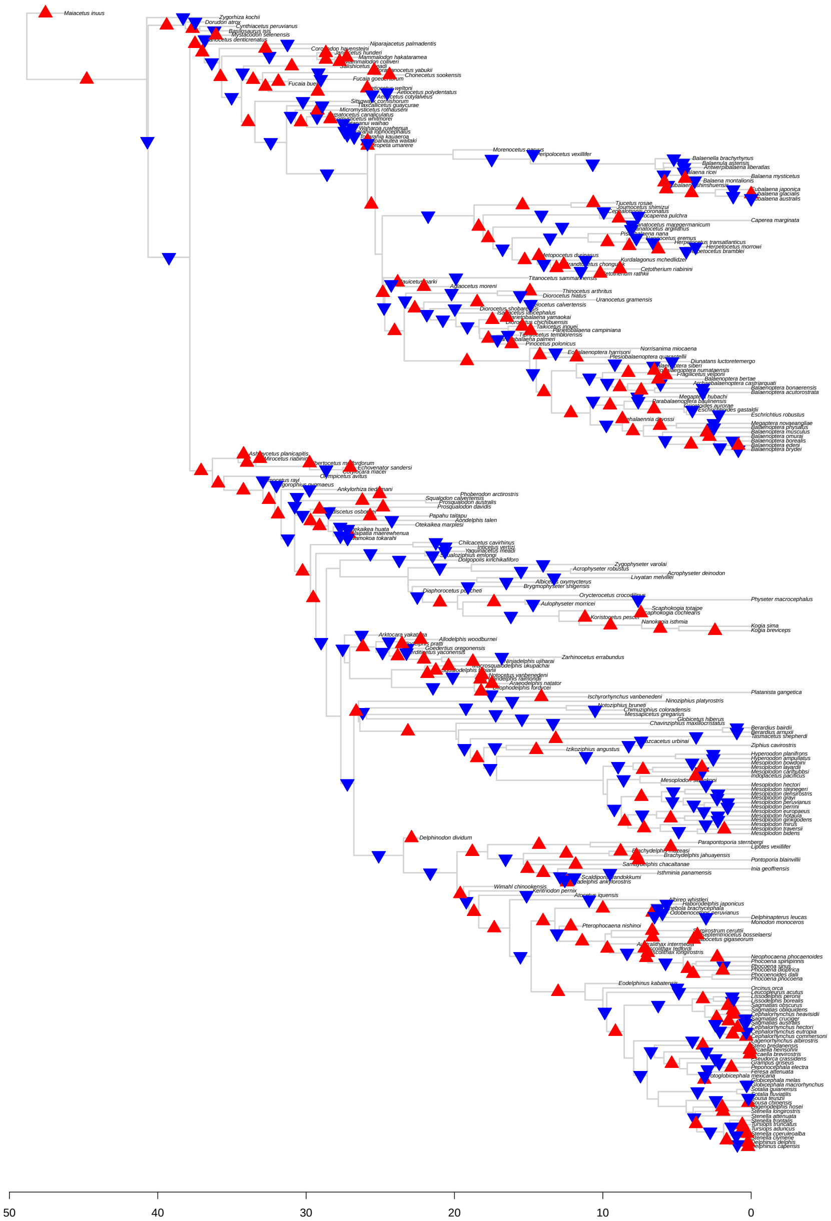

Figure S23: Results for *bayou* fit for the *No Imput* tree setting the average number of shifts in the prior distribution ($\lambda$) to 500. The triangles represent the position and direction of the shifts with posterior probability higher than 0.1, with upward triangles (in red) indicating increases in $\theta$, and downward triangles (in blue) indicating decreases in $\theta$.

##### 3.4.5 *Extant* tree

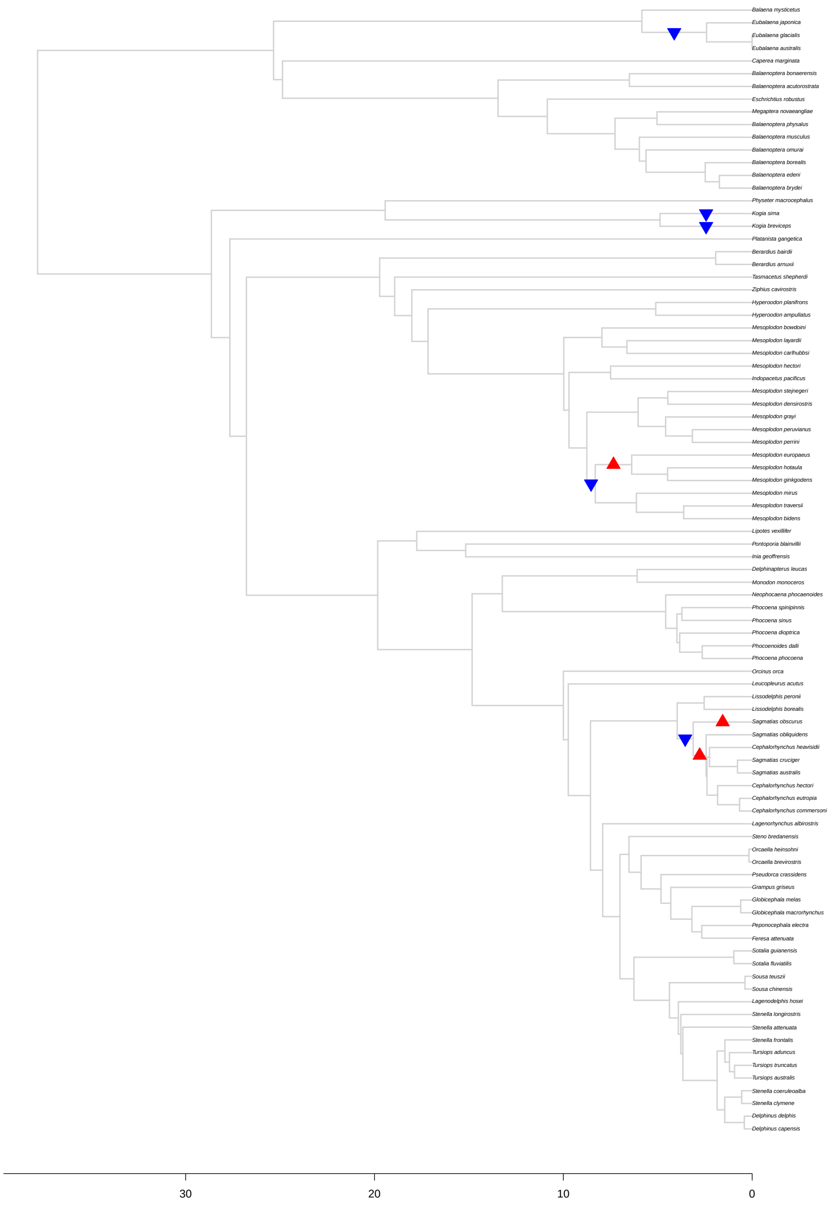

Figure S24: Results for *bayou* fit for the *Extant* tree setting the average number of shifts in the prior distribution ($\lambda$) to 5. The triangles represent the position and direction of the shifts with posterior probability higher than 0.1, with upward triangles (in red) indicating increases in $\theta$, and downward triangles (in blue) indicating decreases in $\theta$..

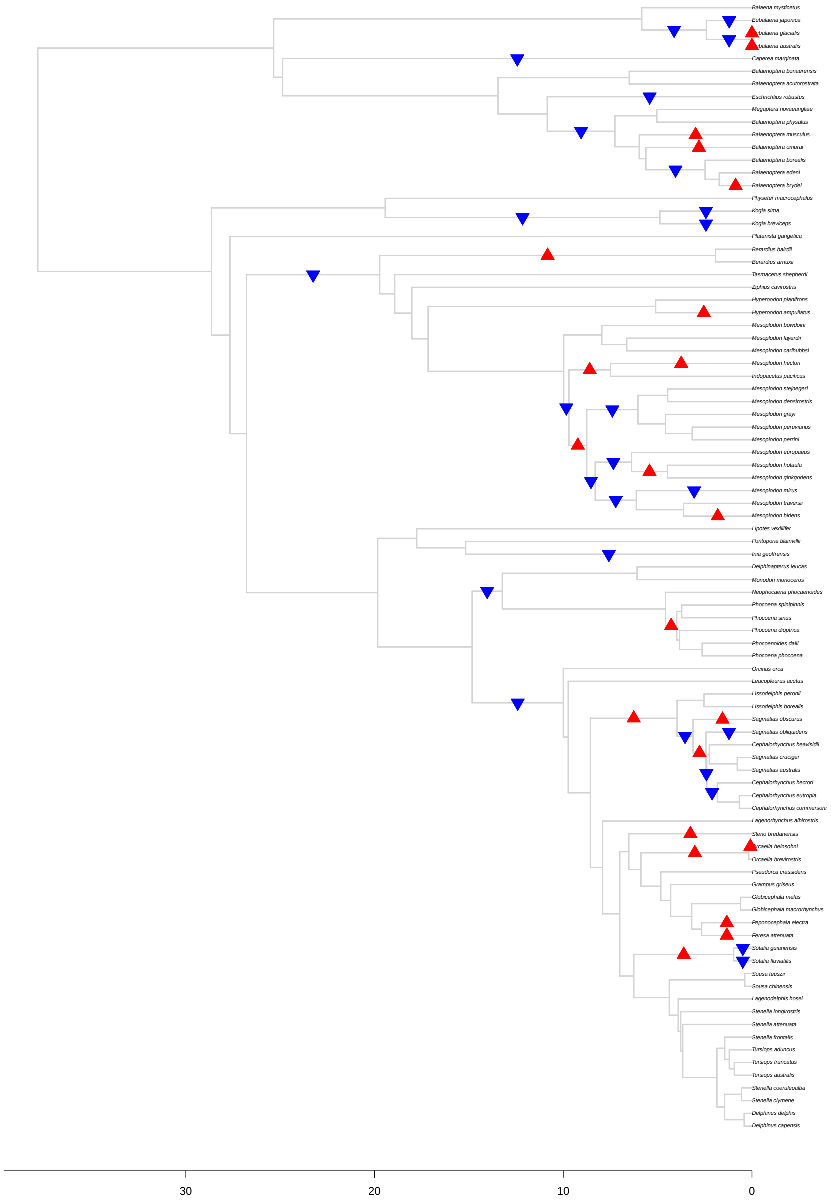

Figure S25: Results for *bayou* fit for the *Extant* tree setting the average number of shifts in the prior distribution ($\lambda$) to 15. The triangles represent the position and direction of the shifts with posterior probability higher than 0.1, with upward triangles (in red) indicating increases in $\theta$, and downward triangles (in blue) indicating decreases in $\theta$.

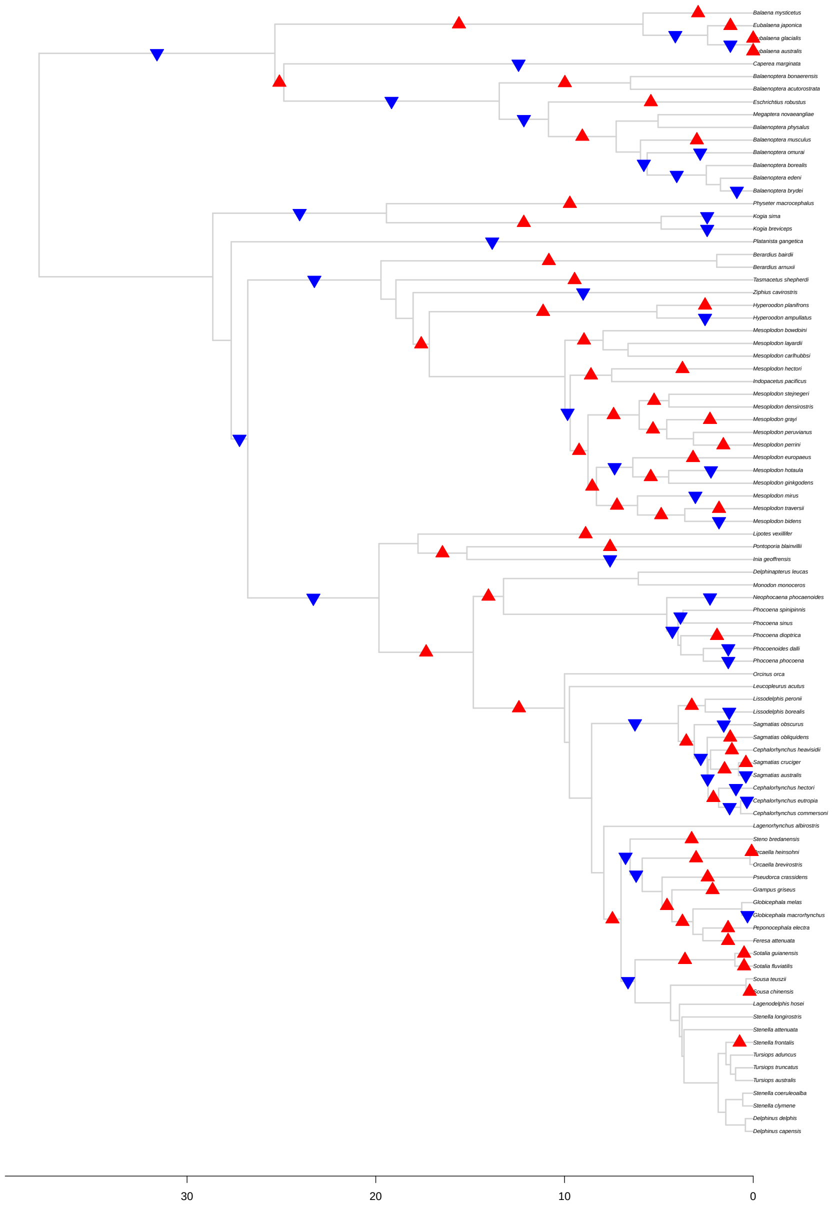

Figure S26: Results for *bayou* fit for the *Extant* tree setting the average number of shifts in the prior distribution ($\lambda$) to 50. The triangles represent the position and direction of the shifts with posterior probability higher than 0.1, with upward triangles (in red) indicating increases in $\theta$, and downward triangles (in blue) indicating decreases in $\theta$.

##### 3.4.6 *Baleen* tree

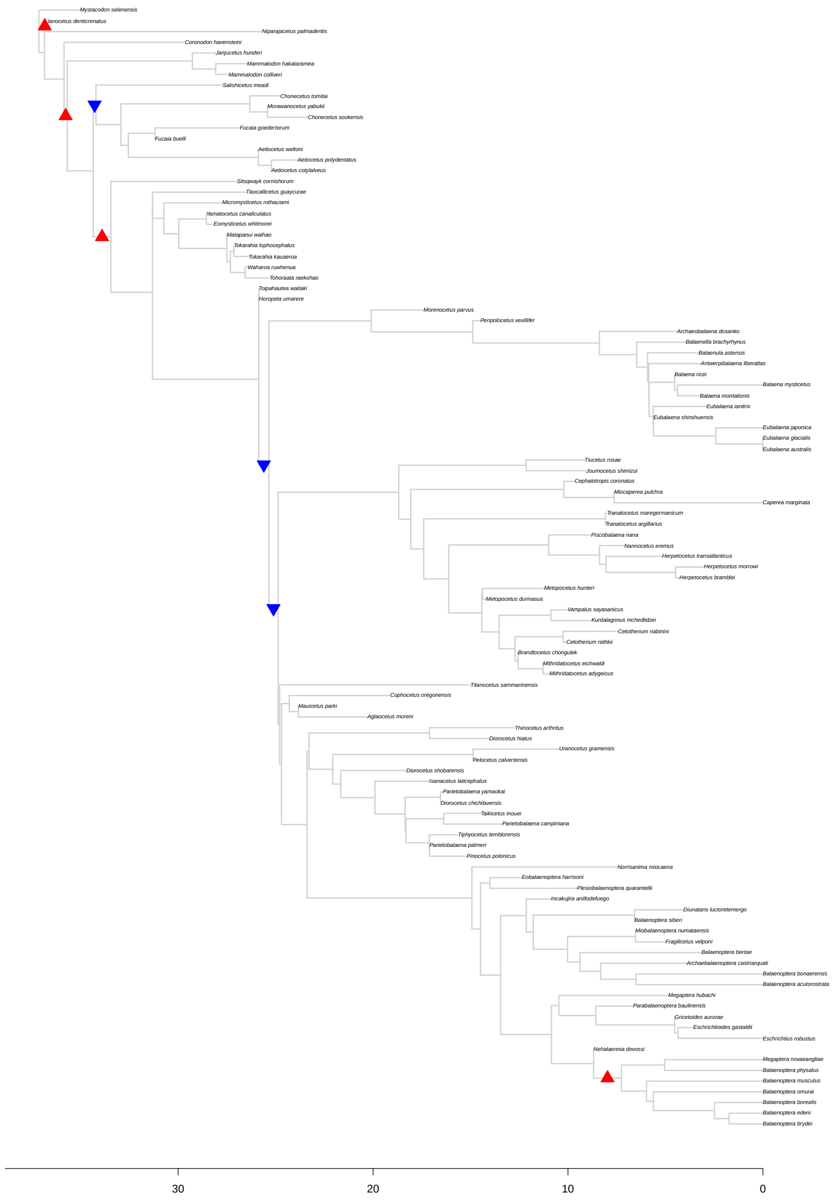

Figure S27: Results for *bayou* fit for the *Baleen* tree setting the average number of shifts in the prior distribution ($\lambda$) to 5. The triangles represent the position and direction of the shifts with posterior probability higher than 0.1, with upward triangles (in red) indicating increases in $\theta$, and downward triangles (in blue) indicating decreases in $\theta$.

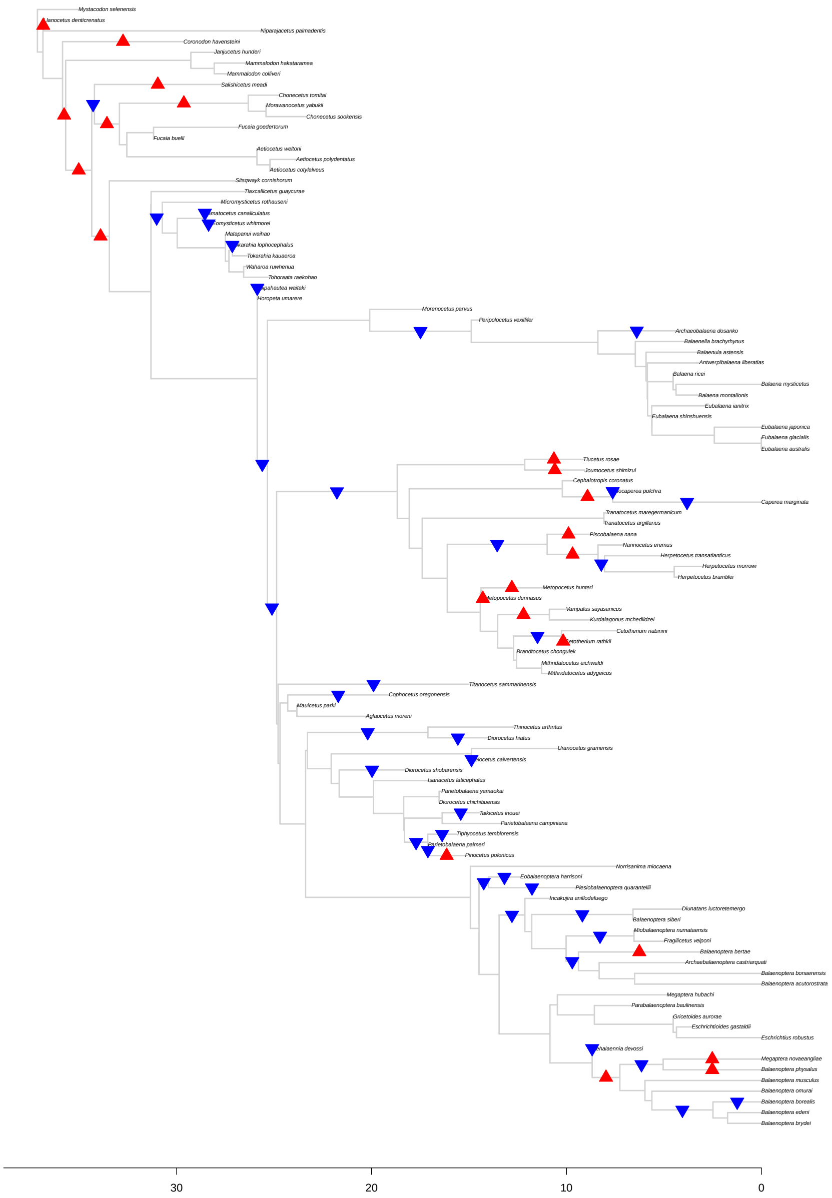

Figure S28: Results for *bayou* fit for the *Baleen* tree setting the average number of shifts in the prior distribution ($\lambda$) to 15. The triangles represent the position and direction of the shifts with posterior probability higher than 0.1, with upward triangles (in red) indicating increases in $\theta$, and downward triangles (in blue) indicating decreases in $\theta$.

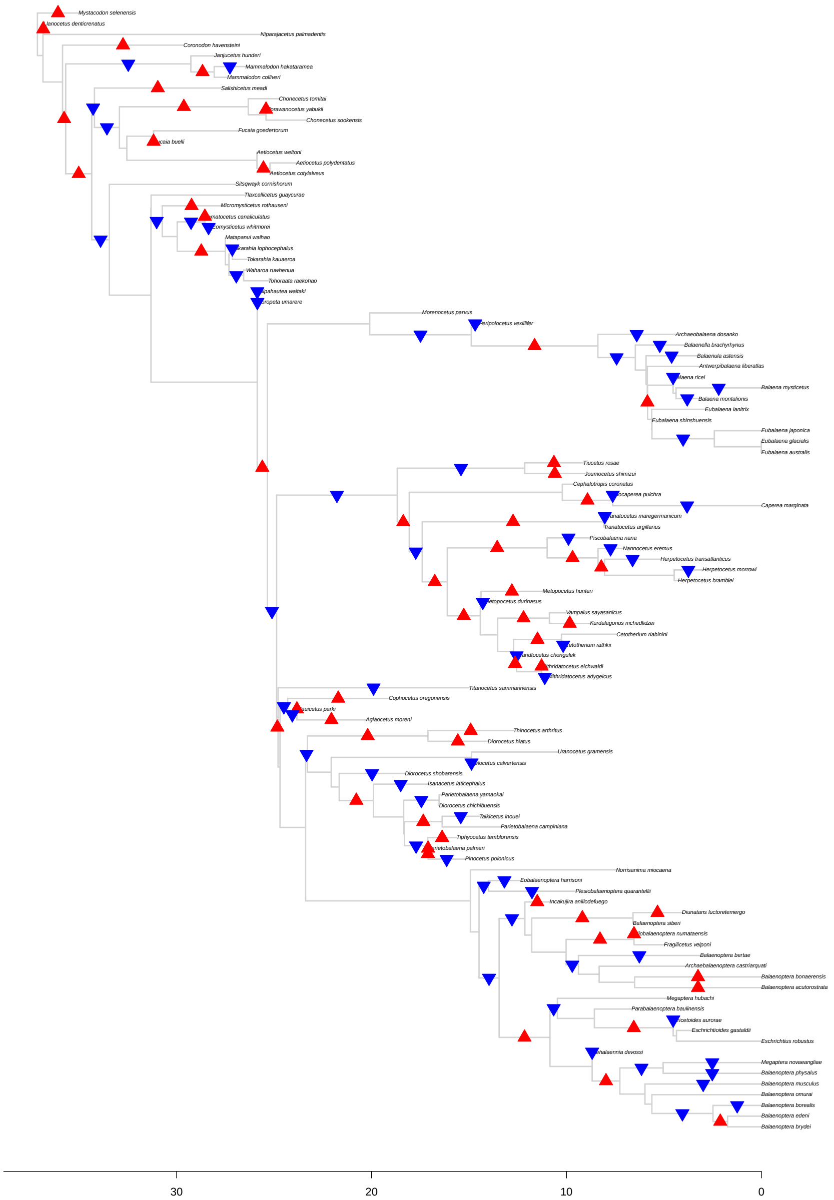

Figure S29: Results for *bayou* fit for the *Baleen* tree setting the average number of shifts in the prior distribution ($\lambda$) to 50. The triangles represent the position and direction of the shifts with posterior probability higher than 0.1, with upward triangles (in red) indicating increases in $\theta$, and downward triangles (in blue) indicating decreases in $\theta$.

##### 3.4.7 *Toothed* tree

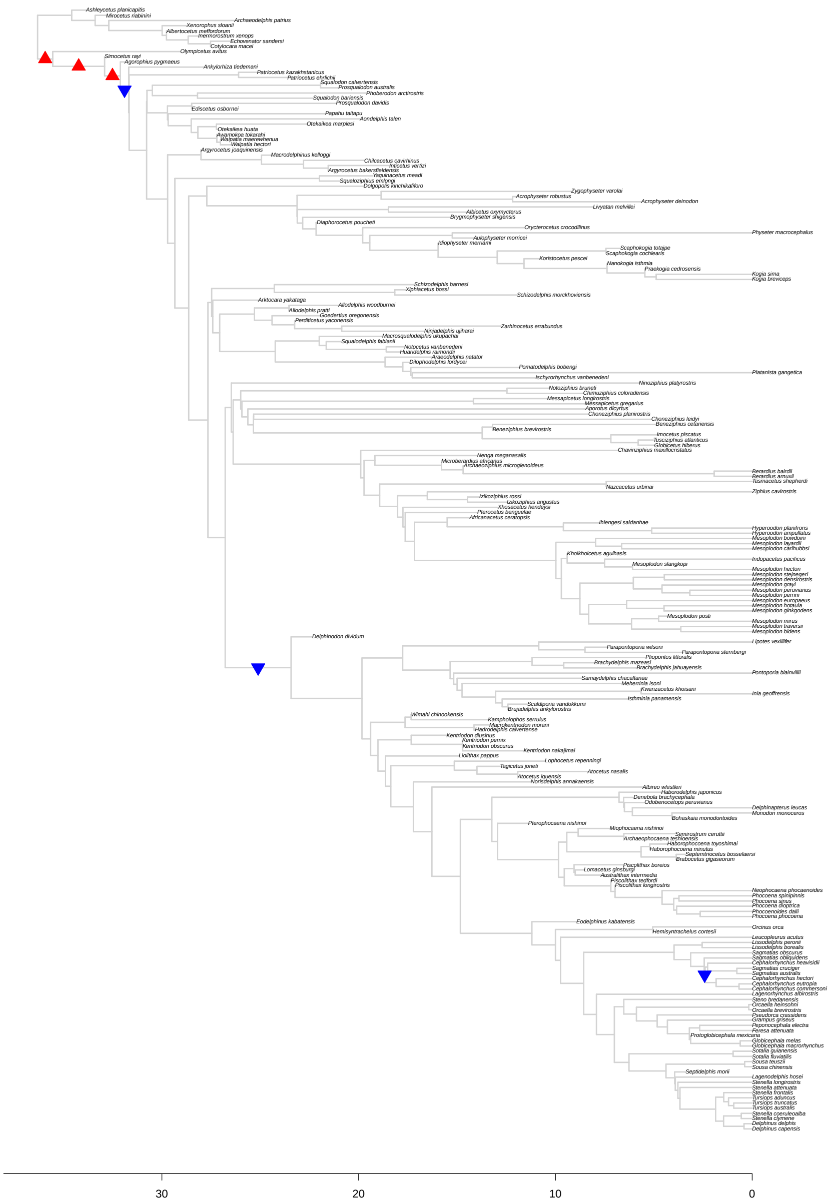

Figure S30: Results for *bayou* fit for the *Toothed* tree setting the average number of shifts in the prior distribution ($\lambda$) to 5. The triangles represent the position and direction of the shifts with posterior probability higher than 0.1, with upward triangles (in red) indicating increases in $\theta$, and downward triangles (in blue) indicating decreases in $\theta$.

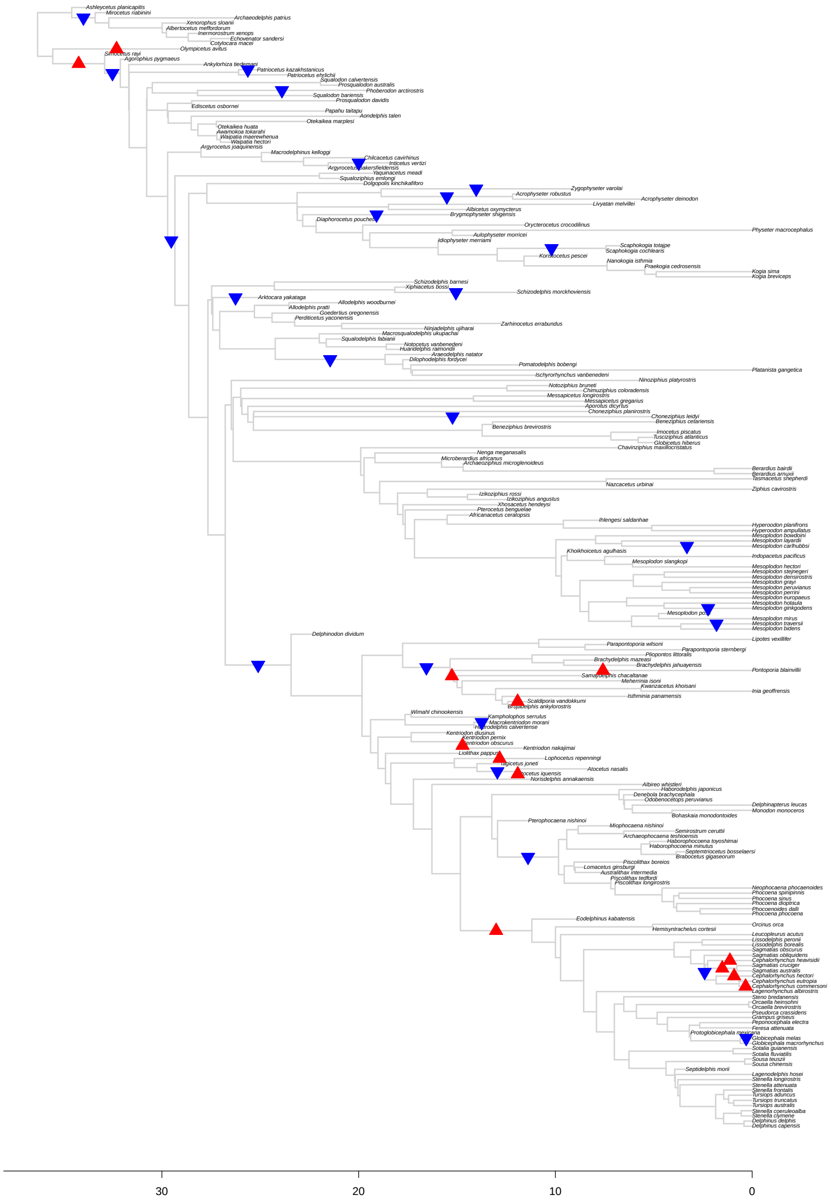

Figure S31: Results for *bayou* fit for the *Toothed* tree setting the average number of shifts in the prior distribution ($\lambda$) to 15. The triangles represent the position and direction of the shifts with posterior probability higher than 0.1, with upward triangles (in red) indicating increases in $\theta$, and downward triangles (in blue) indicating decreases in $\theta$.

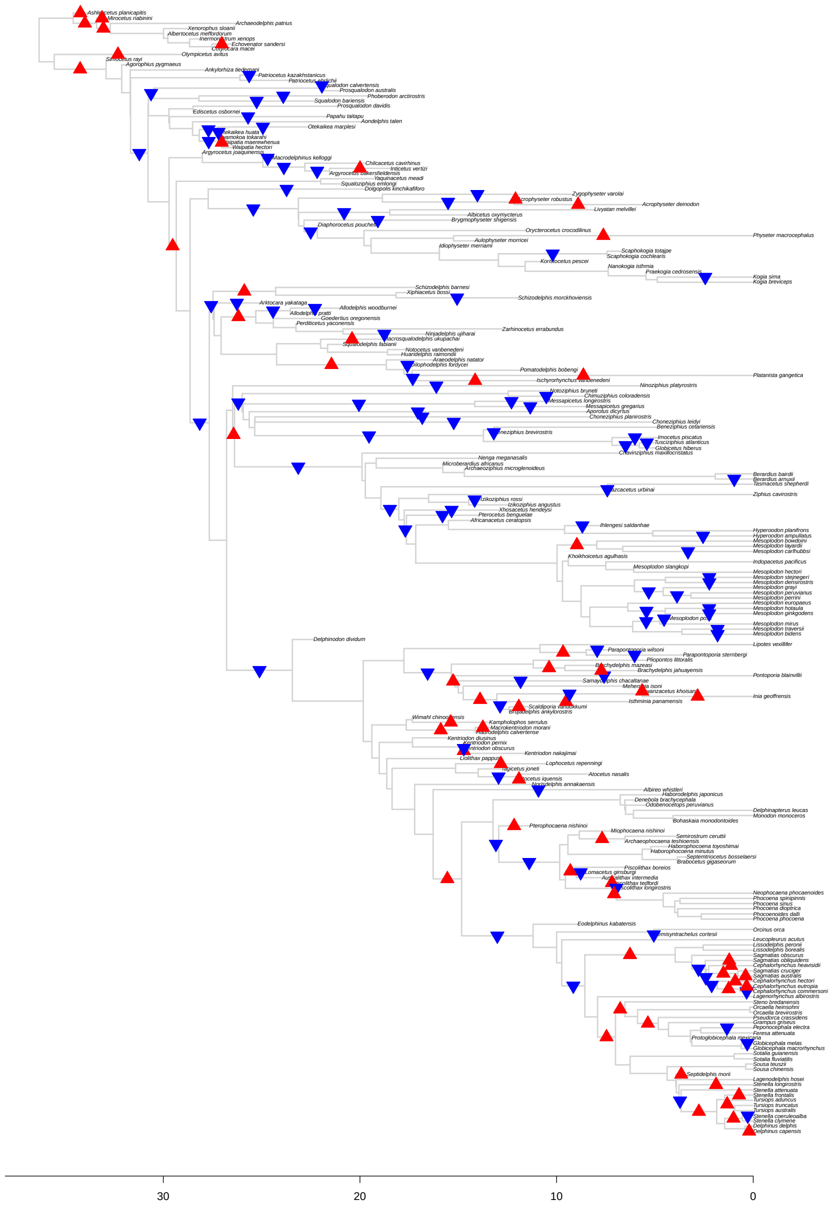

Figure S32: Results for *bayou* fit for the *Toothed* tree setting the average number of shifts in the prior distribution ($\lambda$) to 50. The triangles represent the position and direction of the shifts with posterior probability higher than 0.1, with upward triangles (in red) indicating increases in $\theta$, and downward triangles (in blue) indicating decreases in $\theta$.

#### 3.5 Supplementary Results: Removing taxa with zero-length branches

##### 3.5.1 *Full* tree

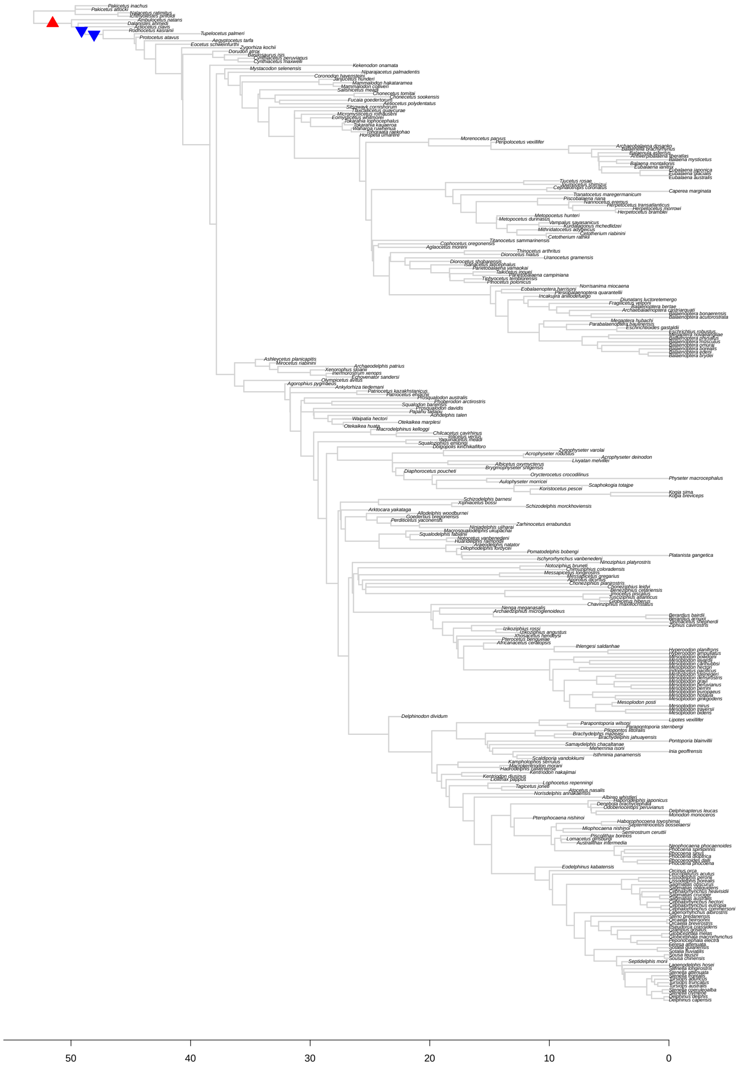

Figure S33: Results for *bayou* fit for the *Full* tree, excluding taxa on zero-length branches, setting the average number of shifts in the prior distribution ($\lambda$) to 5. The triangles represent the position and direction of the shifts with posterior probability higher than 0.1, with upward triangles (in red) indicating increases in $\theta$, and downward triangles (in blue) indicating decreases in $\theta$.

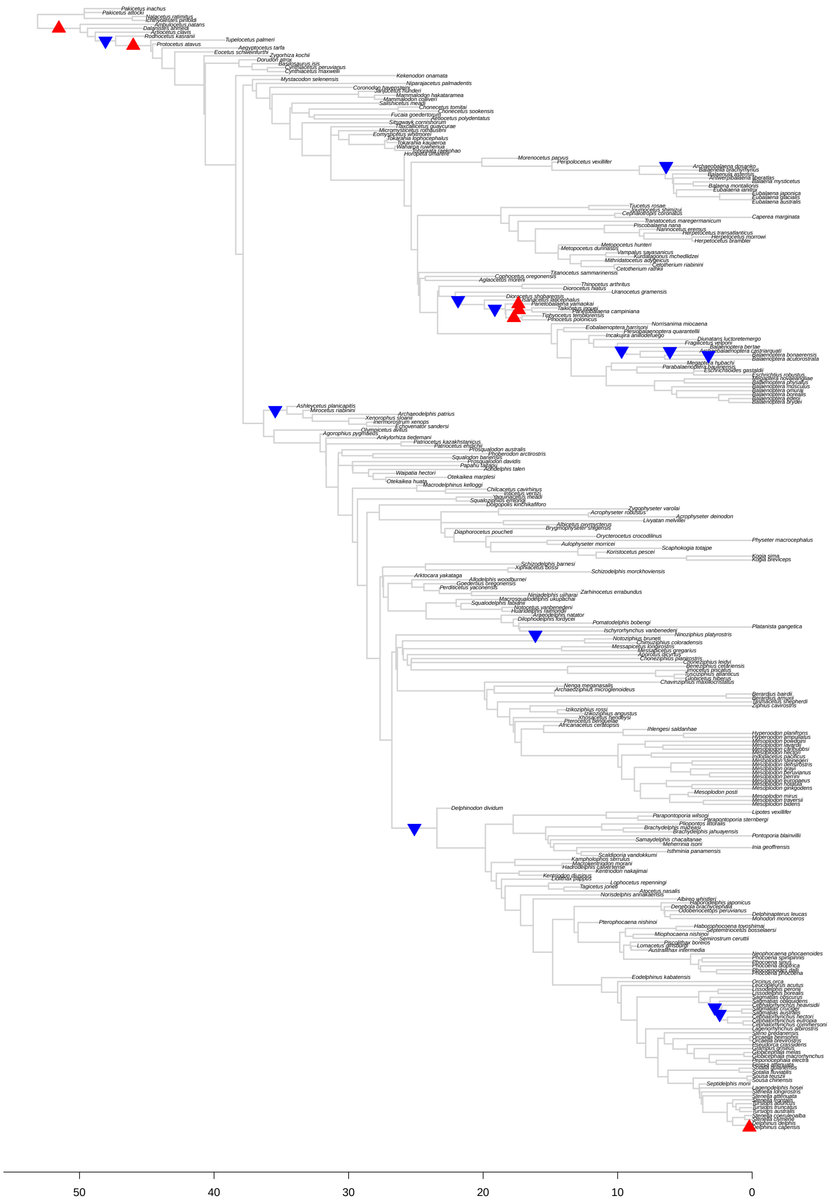

Figure S34: Results for *bayou* fit for the *Full* tree, excluding taxa on zero-length branches, setting the average number of shifts in the prior distribution ($\lambda$) to 15. The triangles represent the position and direction of the shifts with posterior probability higher than 0.1, with upward triangles (in red) indicating increases in $\theta$, and downward triangles (in blue) indicating decreases in $\theta$.

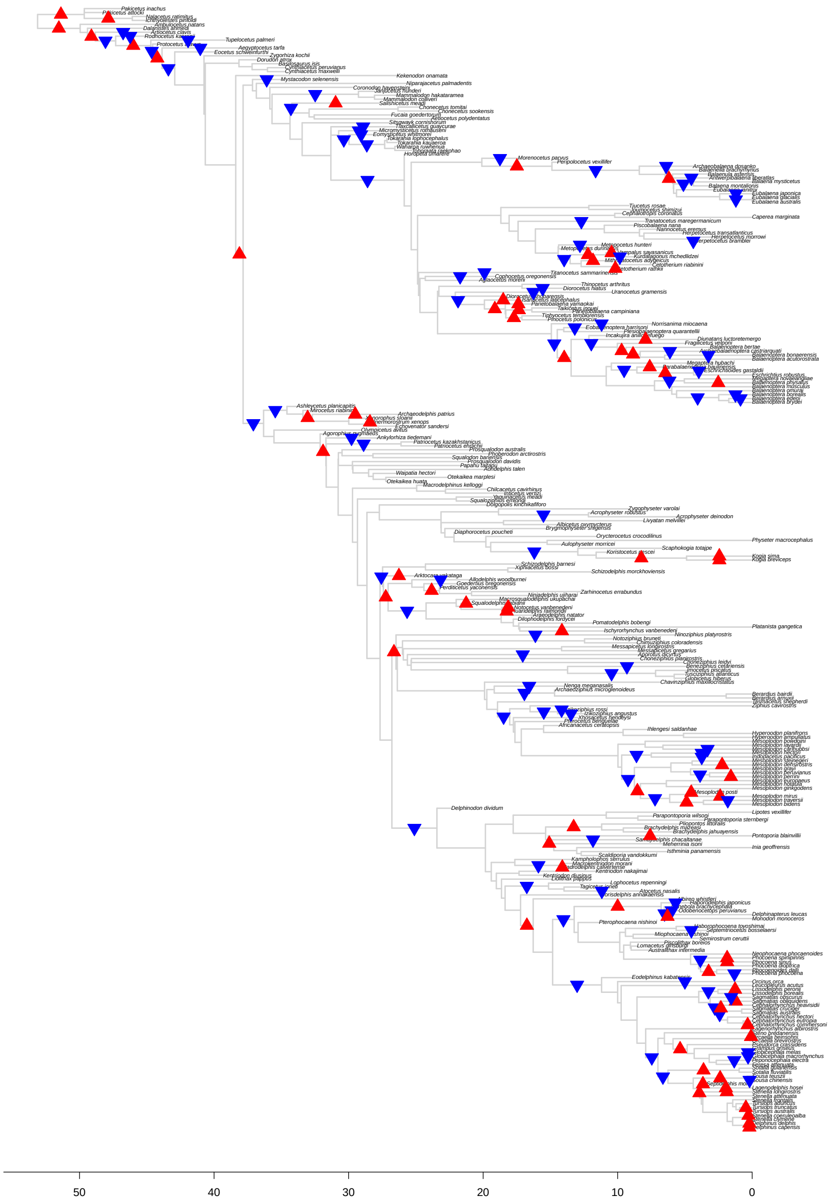

Figure S35: Results for *bayou* fit for the *Full* tree, excluding taxa on zero-length branches, setting the average number of shifts in the prior distribution ($\lambda$) to 50. The triangles represent the position and direction of the shifts with posterior probability higher than 0.1, with upward triangles (in red) indicating increases in $\theta$, and downward triangles (in blue) indicating decreases in $\theta$.

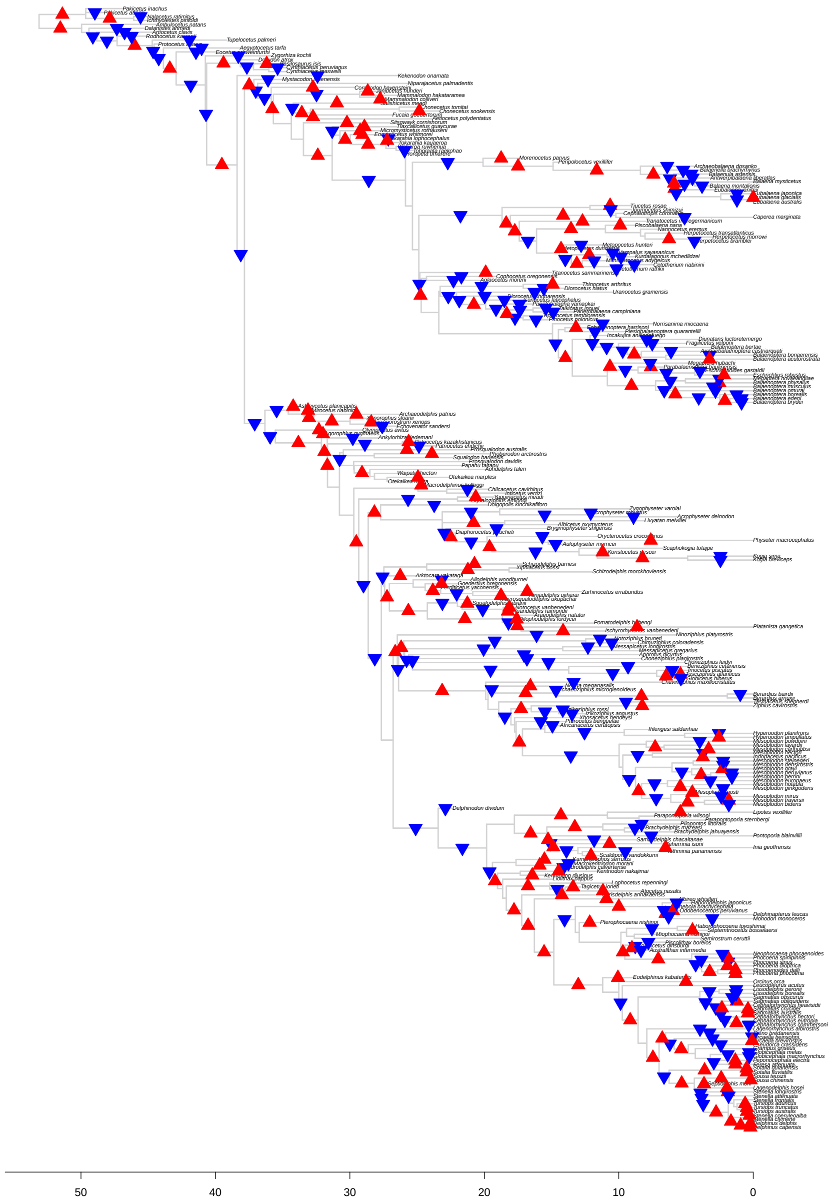

Figure S36: Results for *bayou* fit for the *Full* tree, excluding taxa on zero-length branches, setting the average number of shifts in the prior distribution ($\lambda$) to 250. The triangles represent the position and direction of the shifts with posterior probability higher than 0.1, with upward triangles (in red) indicating increases in $\theta$, and downward triangles (in blue) indicating decreases in $\theta$.

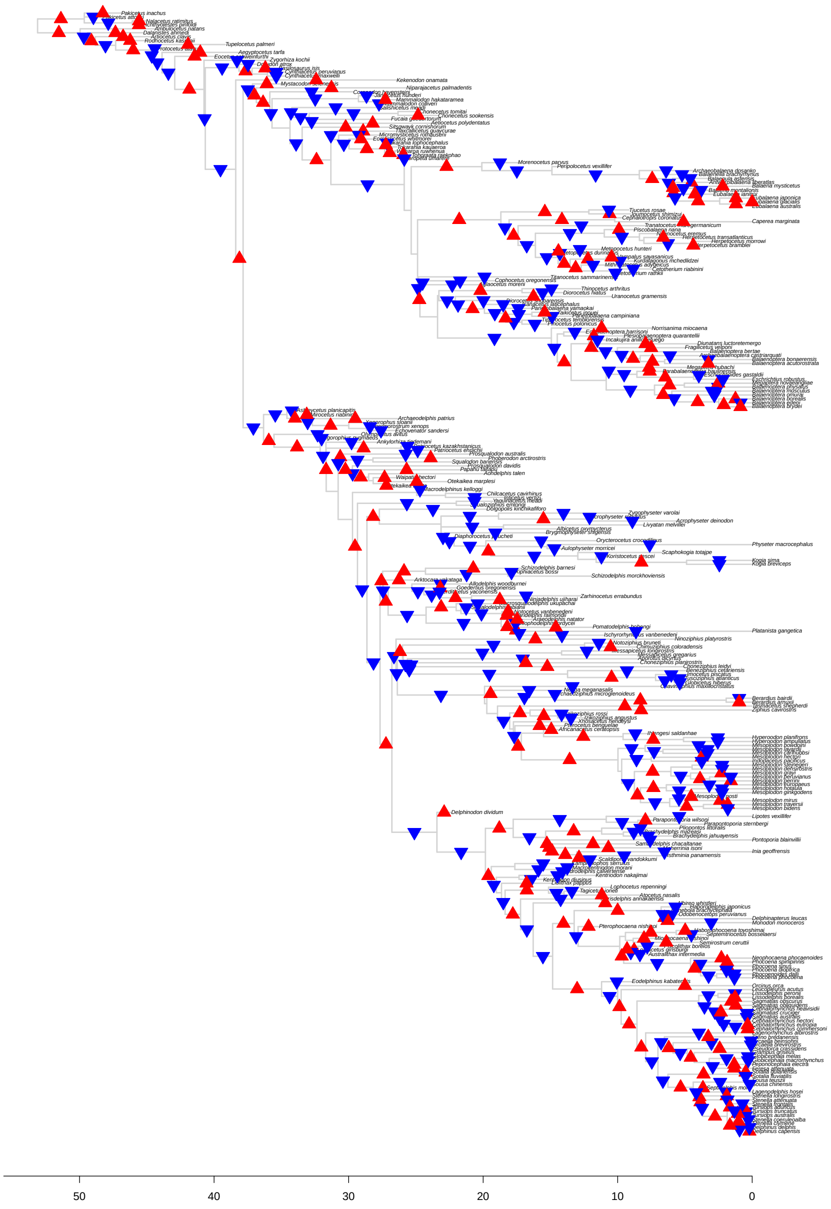

Figure S37: Results for *bayou* fit for the *Full* tree, excluding taxa on zero-length branches, setting the average number of shifts in the prior distribution ($\lambda$) to 500. The triangles represent the position and direction of the shifts with posterior probability higher than 0.1, with upward triangles (in red) indicating increases in $\theta$, and downward triangles (in blue) indicating decreases in $\theta$.

##### 3.5.2 *No Archaeoceti* tree

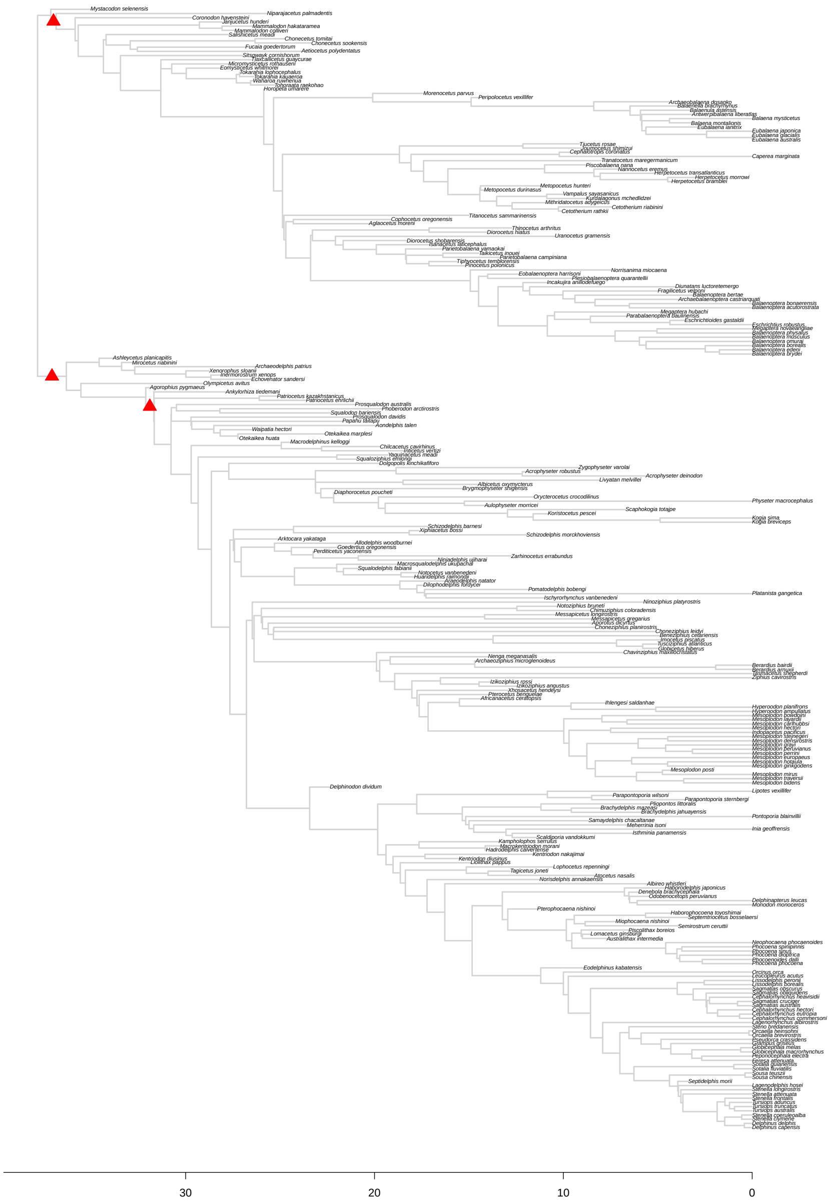

Figure S38: Results for *bayou* fit for the *No Archaeoceti* tree, excluding taxa on zero-length branches, setting the average number of shifts in the prior distribution ($\lambda$) to 5. The triangles represent the position and direction of the shifts with posterior probability higher than 0.1, with upward triangles (in red) indicating increases in $\theta$, and downward triangles (in blue) indicating decreases in $\theta$.

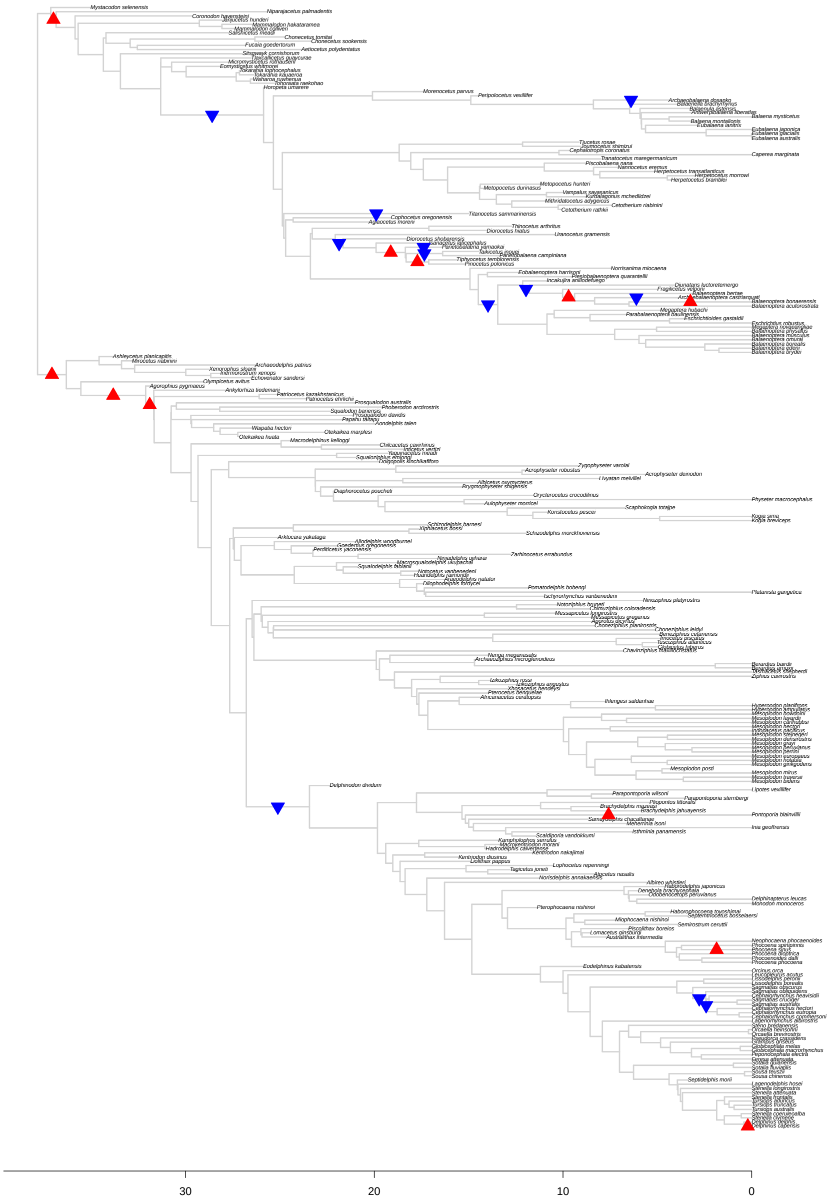

Figure S39: Results for *bayou* fit for the *No Archaeoceti* tree, excluding taxa on zero-length branches, setting the average number of shifts in the prior distribution ($\lambda$) to 15. The triangles represent the position and direction of the shifts with posterior probability higher than 0.1, with upward triangles (in red) indicating increases in $\theta$, and downward triangles (in blue) indicating decreases in $\theta$.

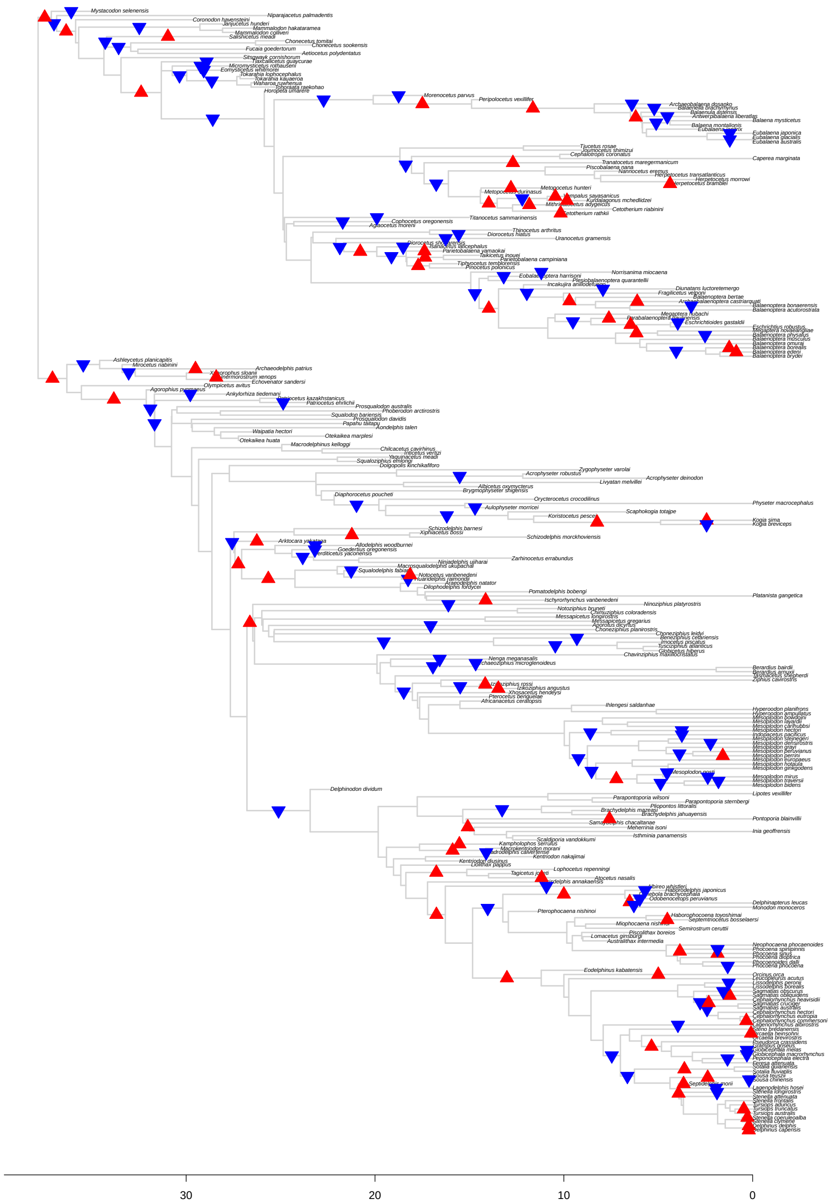

Figure S40: Results for *bayou* fit for the *No Archaeoceti* tree, excluding taxa on zero-length branches, setting the average number of shifts in the prior distribution ($\lambda$) to 50. The triangles represent the position and direction of the shifts with posterior probability higher than 0.1, with upward triangles (in red) indicating increases in $\theta$, and downward triangles (in blue) indicating decreases in $\theta$.

##### 3.5.3 *No Extant* tree

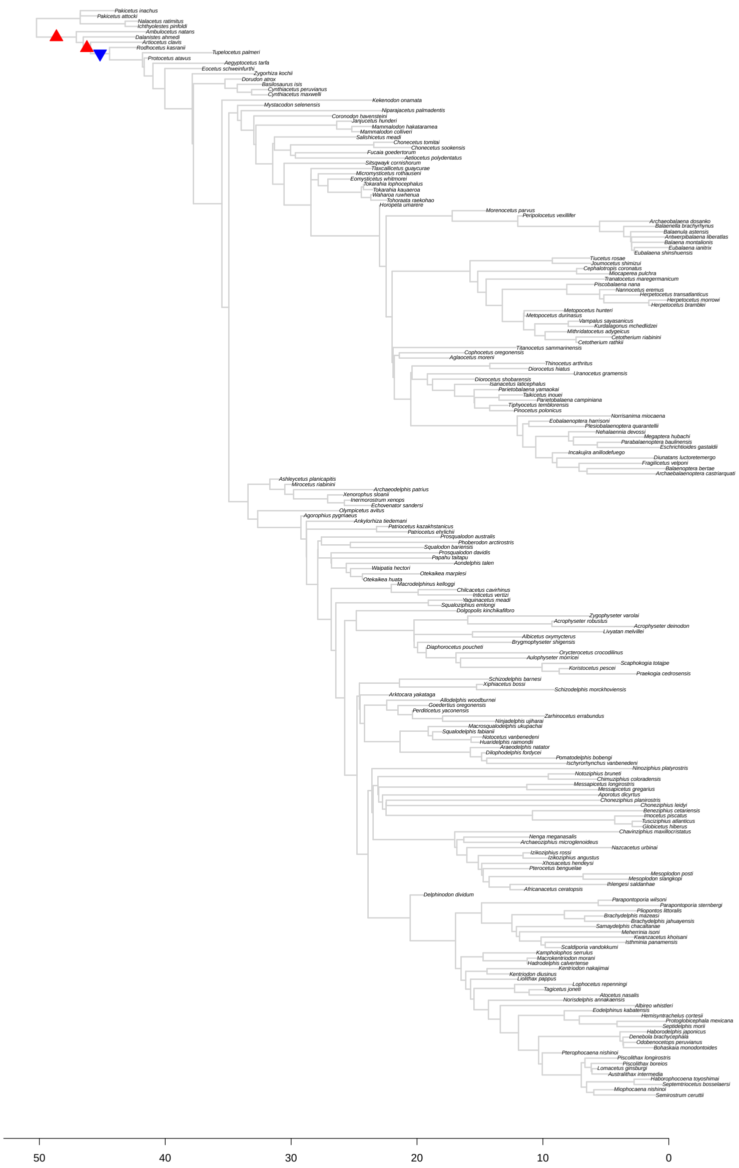

Figure S41: Results for *bayou* fit for the *No Extant* tree, excluding taxa on zero-length branches, setting the average number of shifts in the prior distribution ($\lambda$) to 5. The triangles represent the position and direction of the shifts with posterior probability higher than 0.1, with upward triangles (in red) indicating increases in $\theta$, and downward triangles (in blue) indicating decreases in $\theta$.

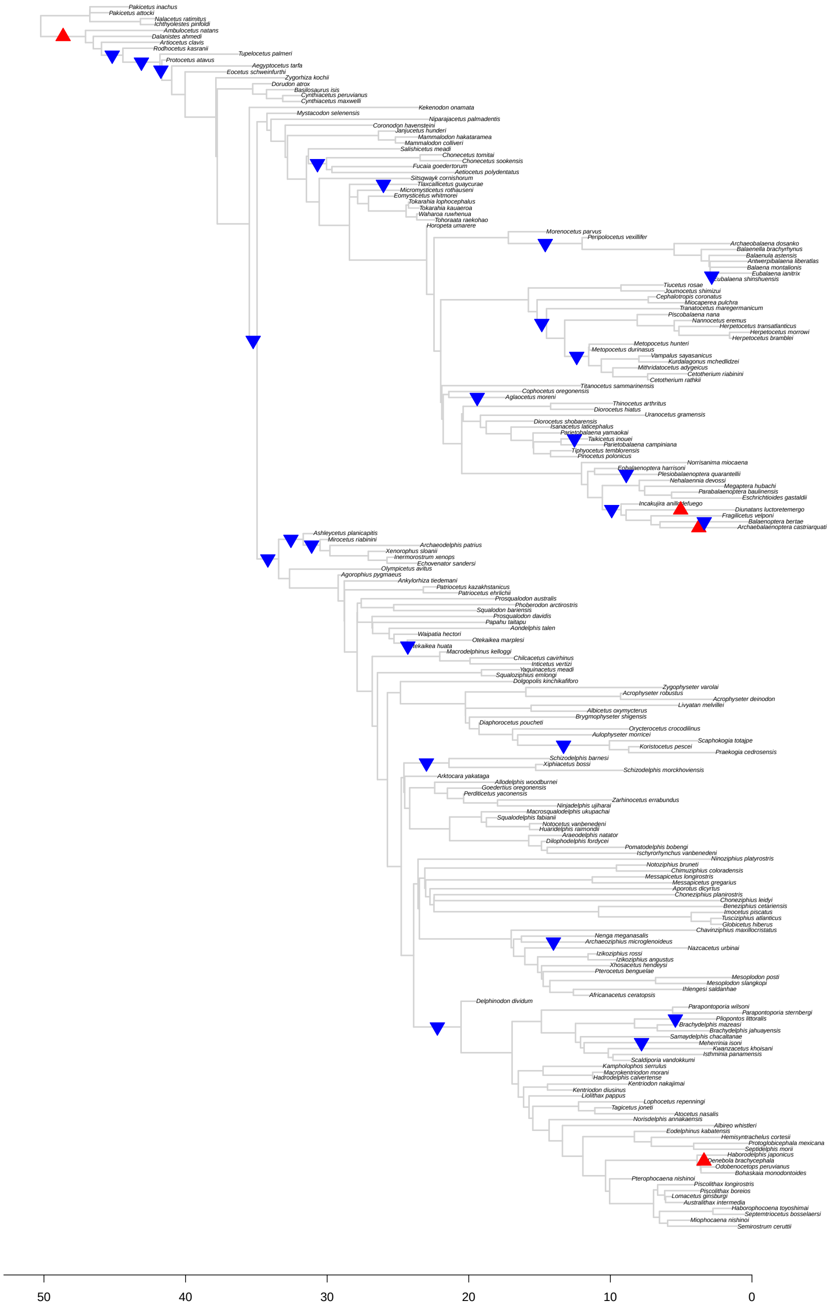

Figure S42: Results for *bayou* fit for the *No Extant* tree, excluding taxa on zero-length branches, setting the average number of shifts in the prior distribution ($\lambda$) to 15. The triangles represent the position and direction of the shifts with posterior probability higher than 0.1, with upward triangles (in red) indicating increases in $\theta$, and downward triangles (in blue) indicating decreases in $\theta$.

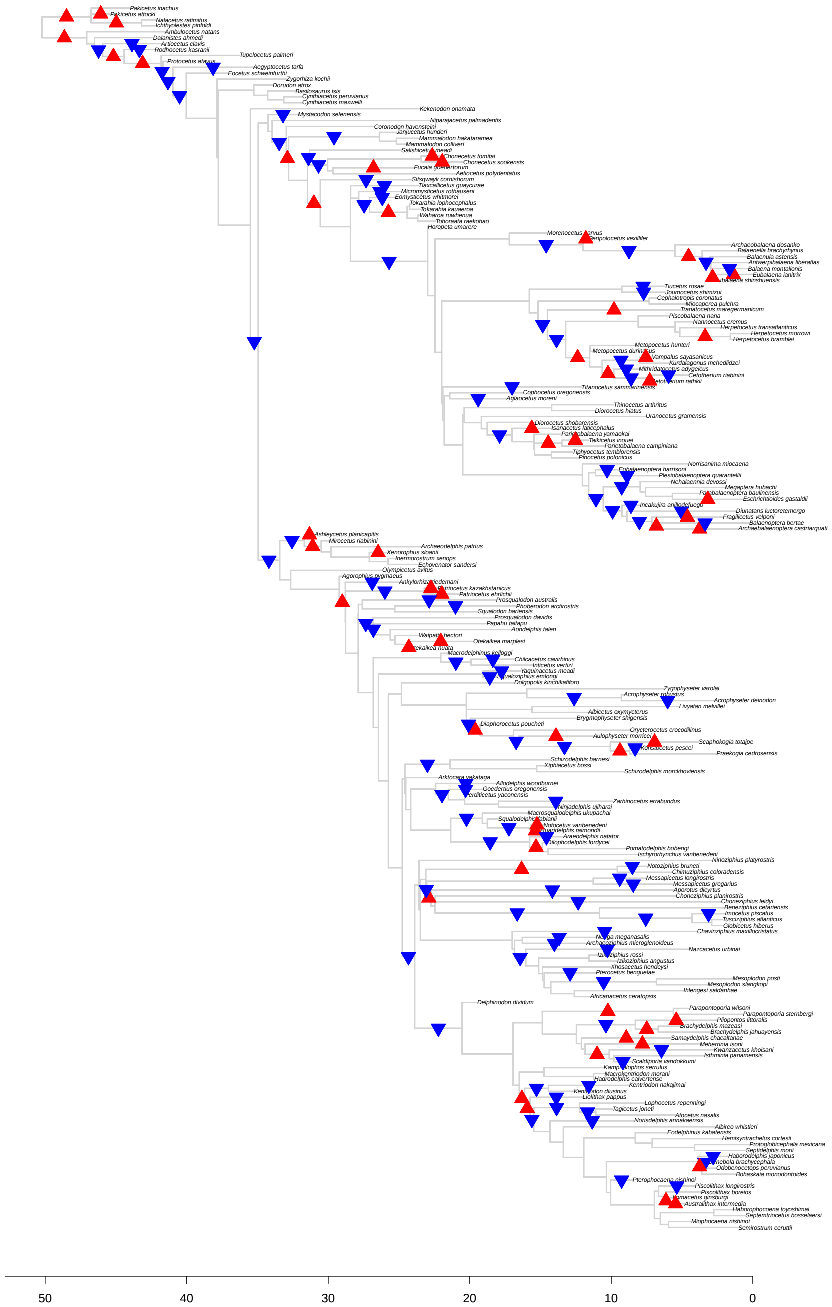

Figure S43: Results for *bayou* fit for the *No Extant* tree, excluding taxa on zero-length branches, setting the average number of shifts in the prior distribution ($\lambda$) to 50. The triangles represent the position and direction of the shifts with posterior probability higher than 0.1, with upward triangles (in red) indicating increases in $\theta$, and downward triangles (in blue) indicating decreases in $\theta$.

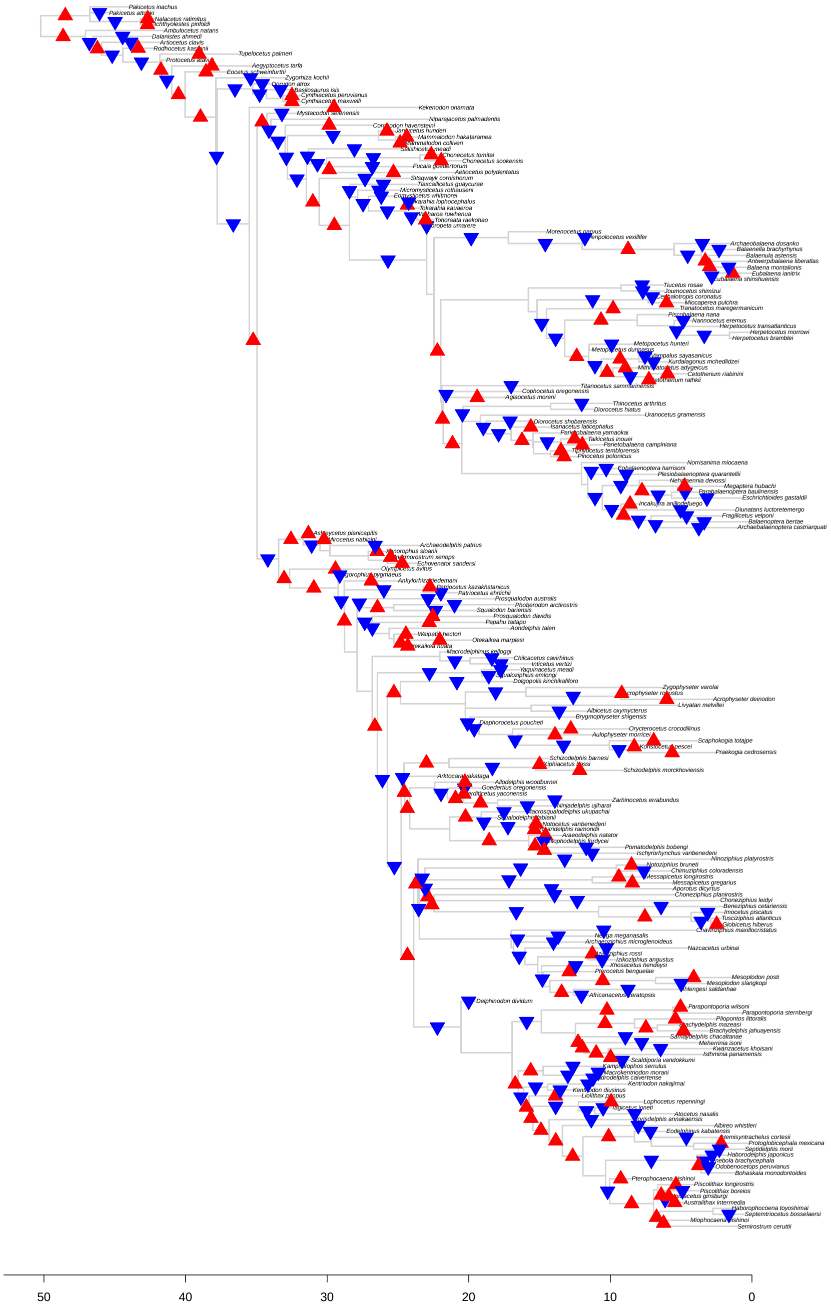

Figure S44: Results for *bayou* fit for the *No Extant* tree, excluding taxa on zero-length branches, setting the average number of shifts in the prior distribution ($\lambda$) to 250. The triangles represent the position and direction of the shifts with posterior probability higher than 0.1, with upward triangles (in red) indicating increases in $\theta$, and downward triangles (in blue) indicating decreases in $\theta$.

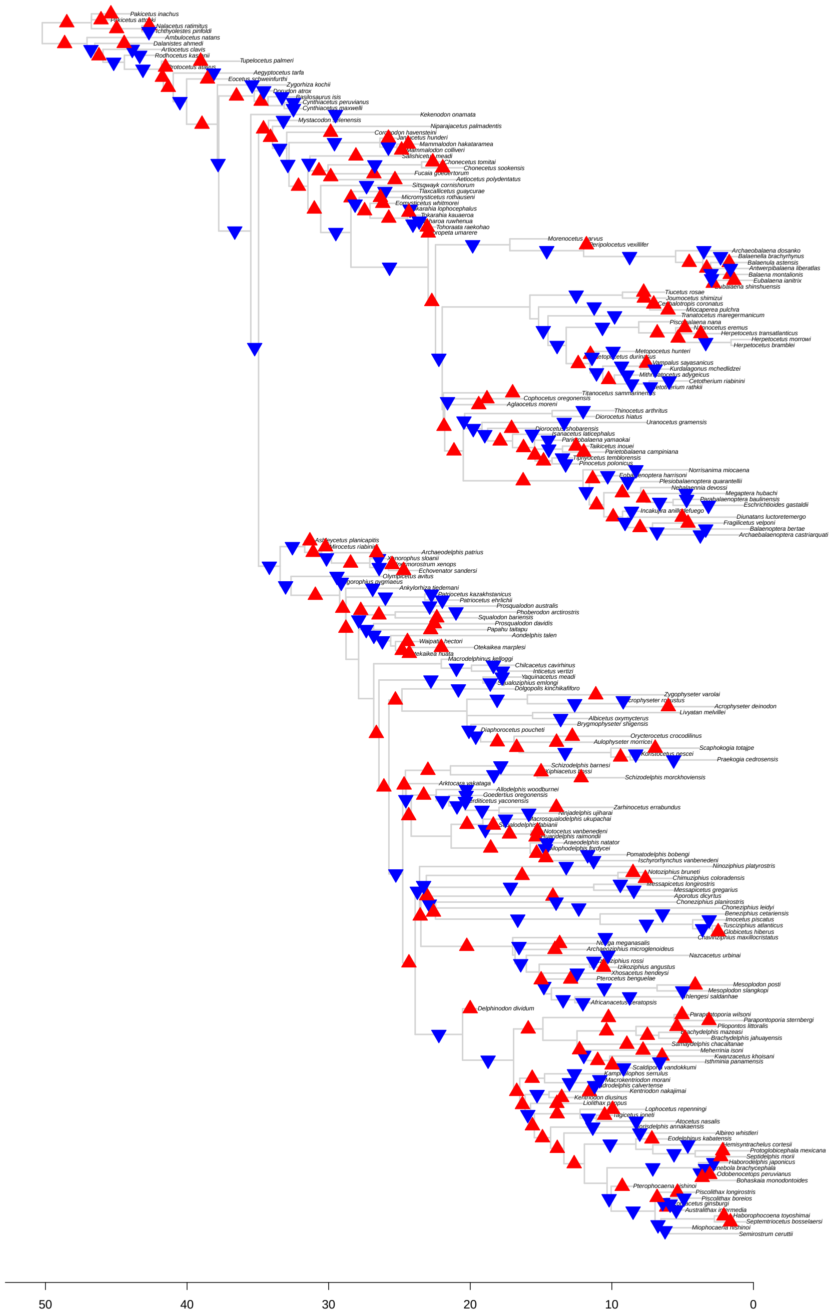

Figure S45: Results for *bayou* fit for the *No Extant* tree, excluding taxa on zero-length branches, setting the average number of shifts in the prior distribution ($\lambda$) to 500. The triangles represent the position and direction of the shifts with posterior probability higher than 0.1, with upward triangles (in red) indicating increases in $\theta$, and downward triangles (in blue) indicating decreases in $\theta$.

##### 3.5.4 *No Imput* tree

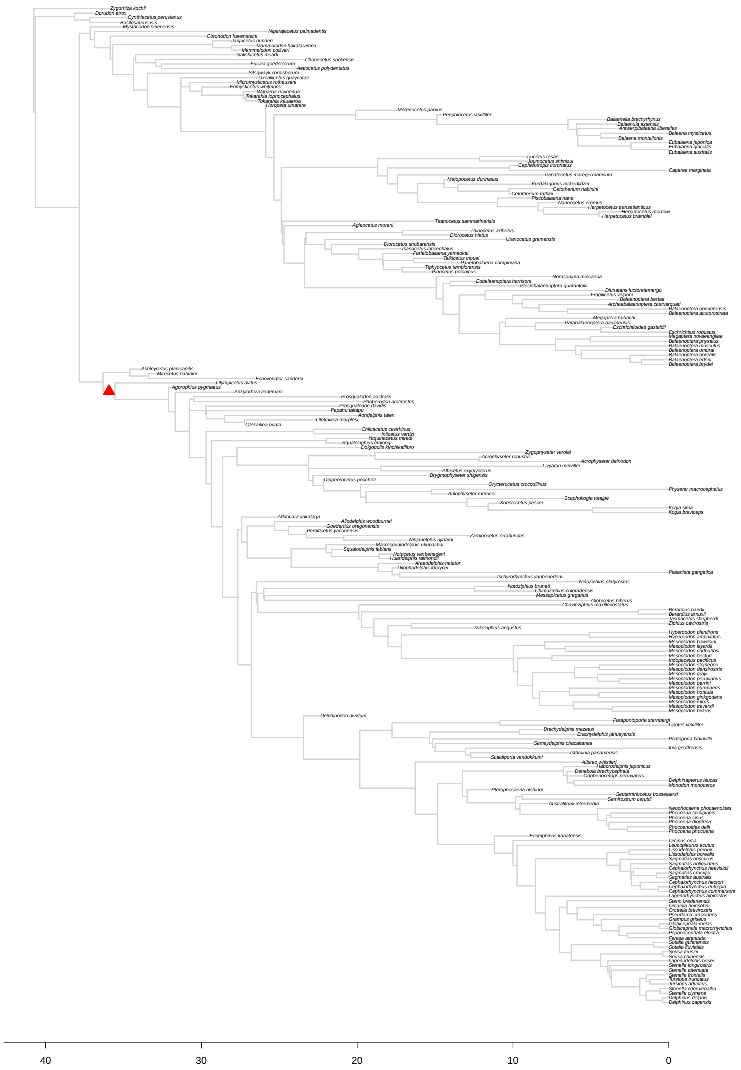

Figure S46: Results for *bayou* fit for the *No Imput* tree, excluding taxa on zero-length branches, setting the average number of shifts in the prior distribution ($\lambda$) to 5. The triangles represent the position and direction of the shifts with posterior probability higher than 0.1, with upward triangles (in red) indicating increases in $\theta$, and downward triangles (in blue) indicating decreases in $\theta$.

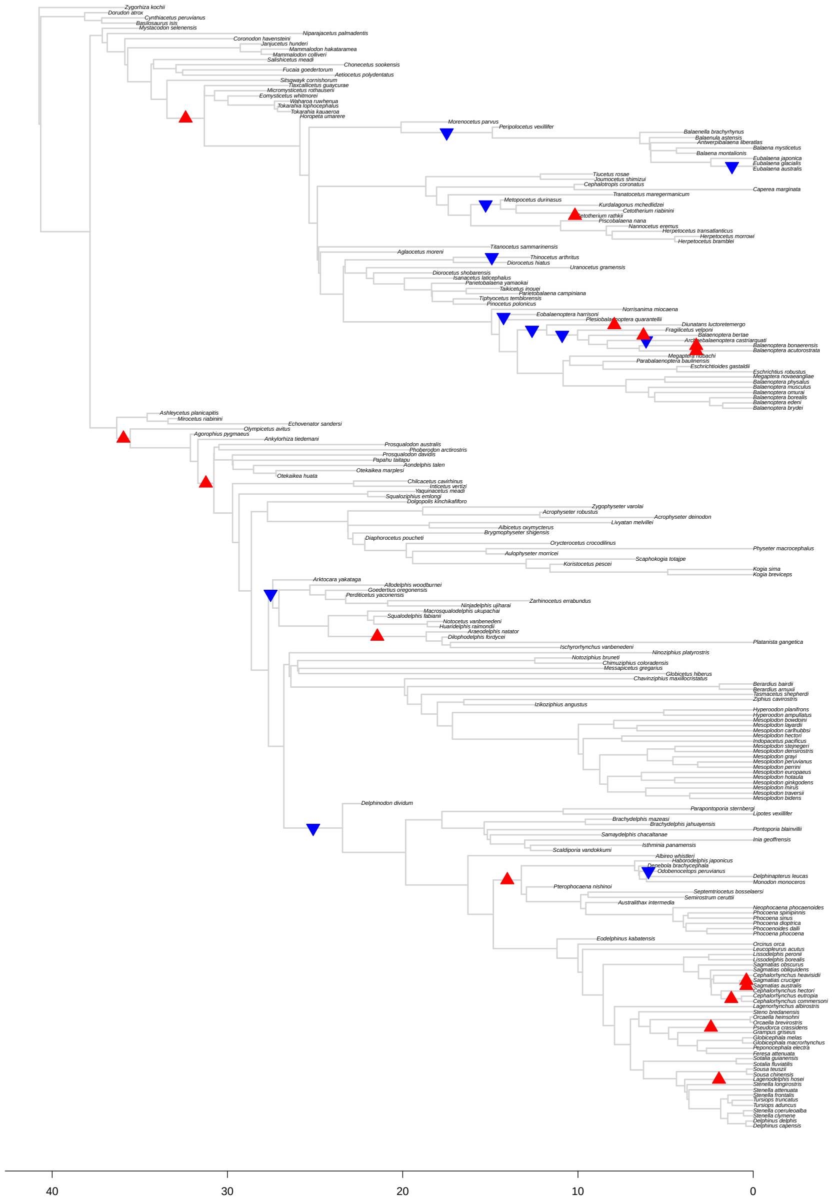

Figure S47: Results for *bayou* fit for the *No Imput* tree, excluding taxa on zero-length branches, setting the average number of shifts in the prior distribution ($\lambda$) to 15. The triangles represent the position and direction of the shifts with posterior probability higher than 0.1, with upward triangles (in red) indicating increases in $\theta$, and downward triangles (in blue) indicating decreases in $\theta$.

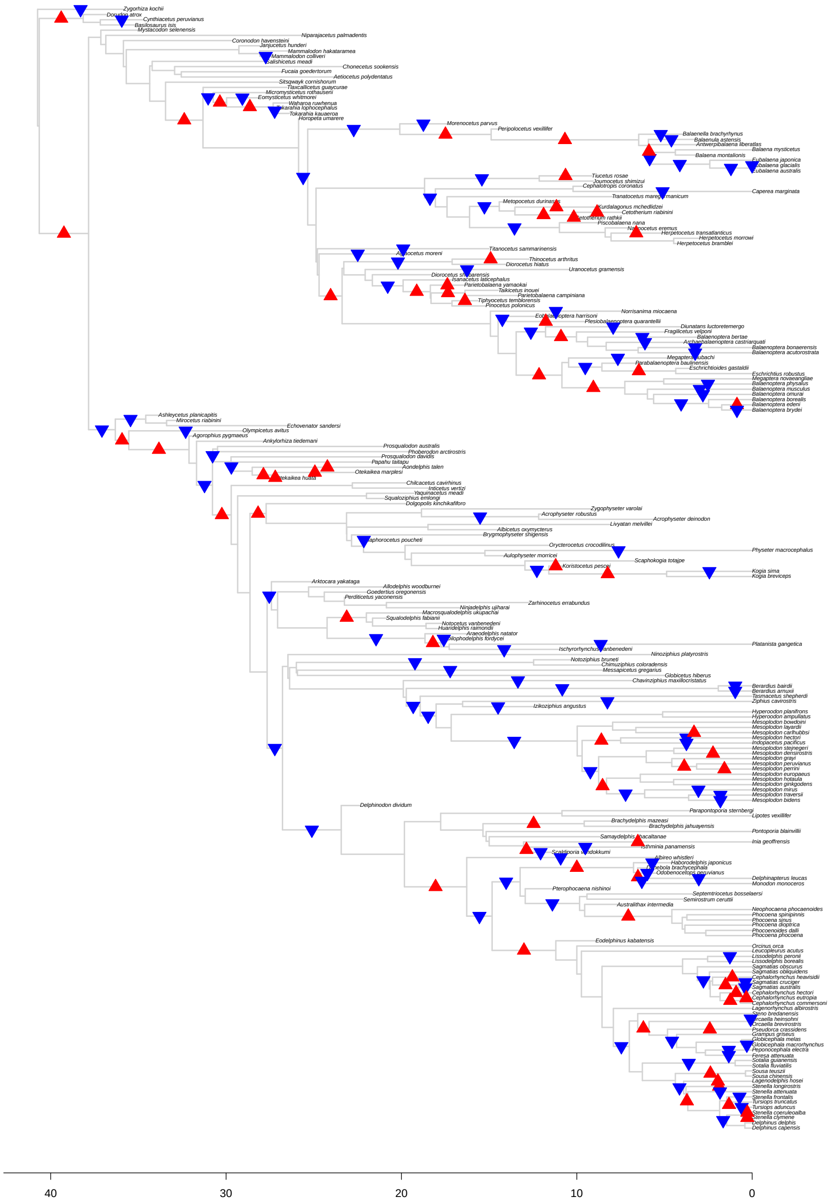

Figure S48: Results for *bayou* fit for the *No Imput* tree, excluding taxa on zero-length branches, setting the average number of shifts in the prior distribution ($\lambda$) to 50. The triangles represent the position and direction of the shifts with posterior probability higher than 0.1, with upward triangles (in red) indicating increases in $\theta$, and downward triangles (in blue) indicating decreases in $\theta$.

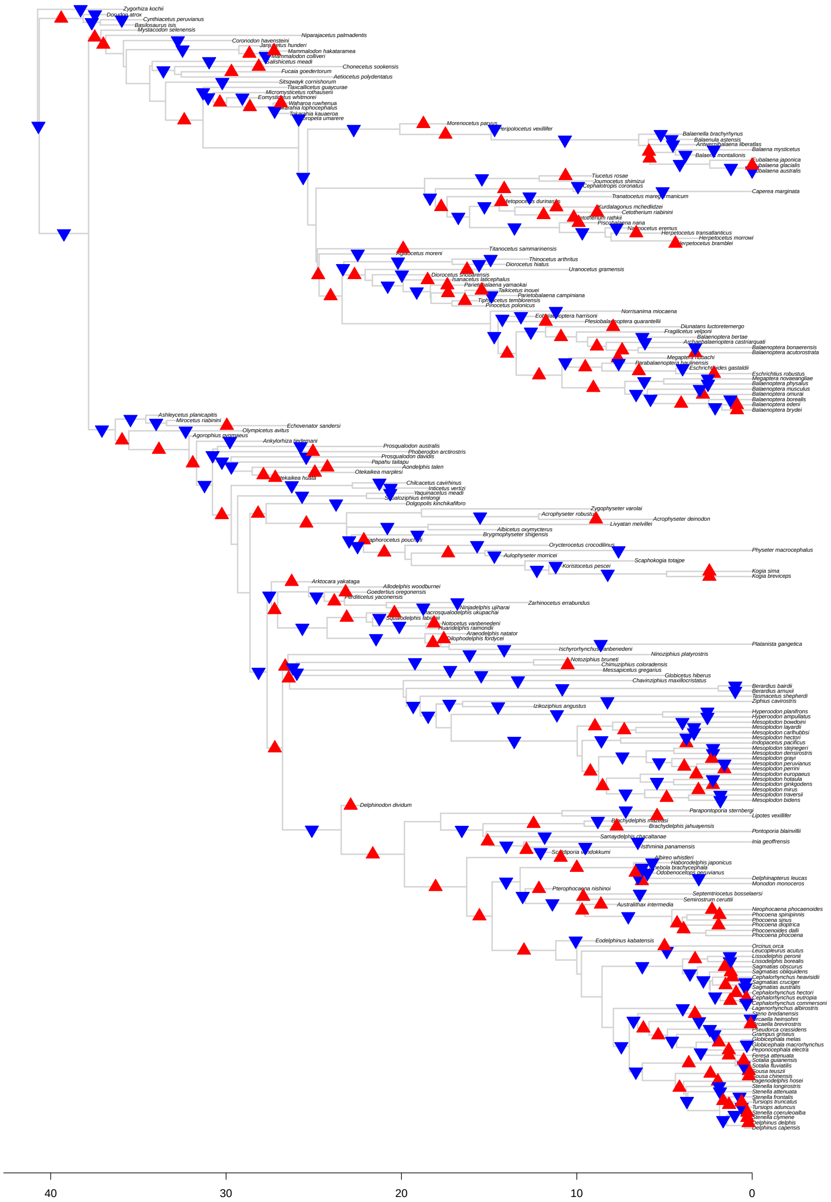

Figure S49: Results for *bayou* fit for the *No Imput* tree, excluding taxa on zero-length branches, setting the average number of shifts in the prior distribution ($\lambda$) to 250. The triangles represent the position and direction of the shifts with posterior probability higher than 0.1, with upward triangles (in red) indicating increases in $\theta$, and downward triangles (in blue) indicating decreases in $\theta$.

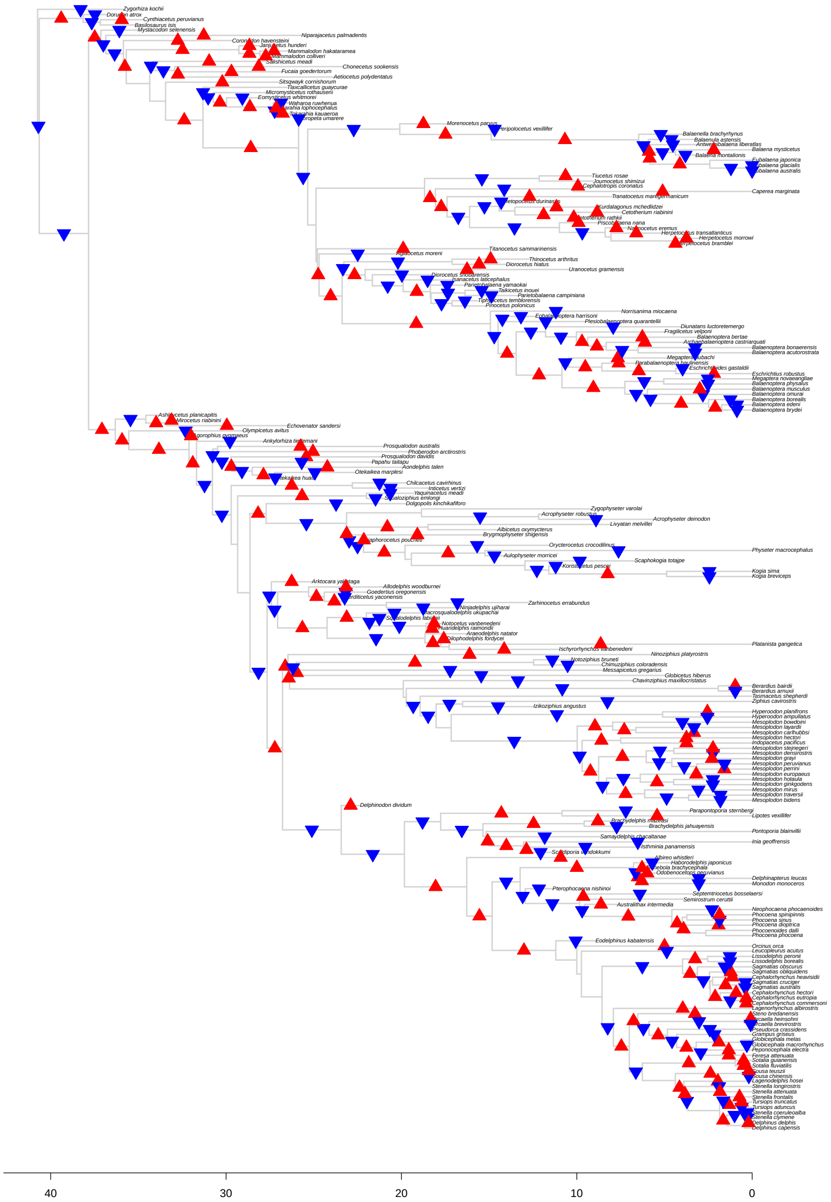

Figure S50: Results for *bayou* fit for the *No Imput* tree, excluding taxa on zero-length branches, setting the average number of shifts in the prior distribution ($\lambda$) to 500. The triangles represent the position and direction of the shifts with posterior probability higher than 0.1, with upward triangles (in red) indicating increases in $\theta$, and downward triangles (in blue) indicating decreases in $\theta$.

##### 3.5.5 *Baleen* tree

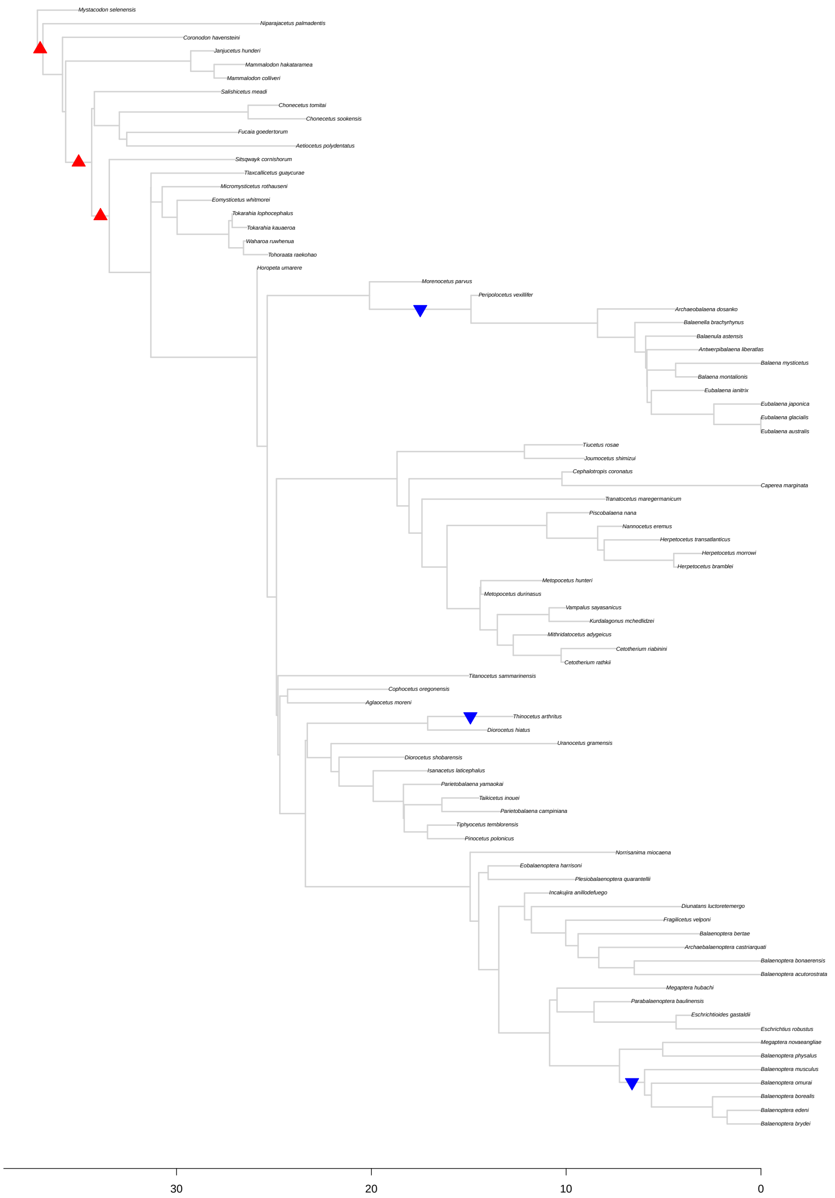

Figure S51: Results for *bayou* fit for the *Baleen* tree, excluding taxa on zero-length branches, setting the average number of shifts in the prior distribution ($\lambda$) to 5. The triangles represent the position and direction of the shifts with posterior probability higher than 0.1, with upward triangles (in red) indicating increases in $\theta$, and downward triangles (in blue) indicating decreases in $\theta$.

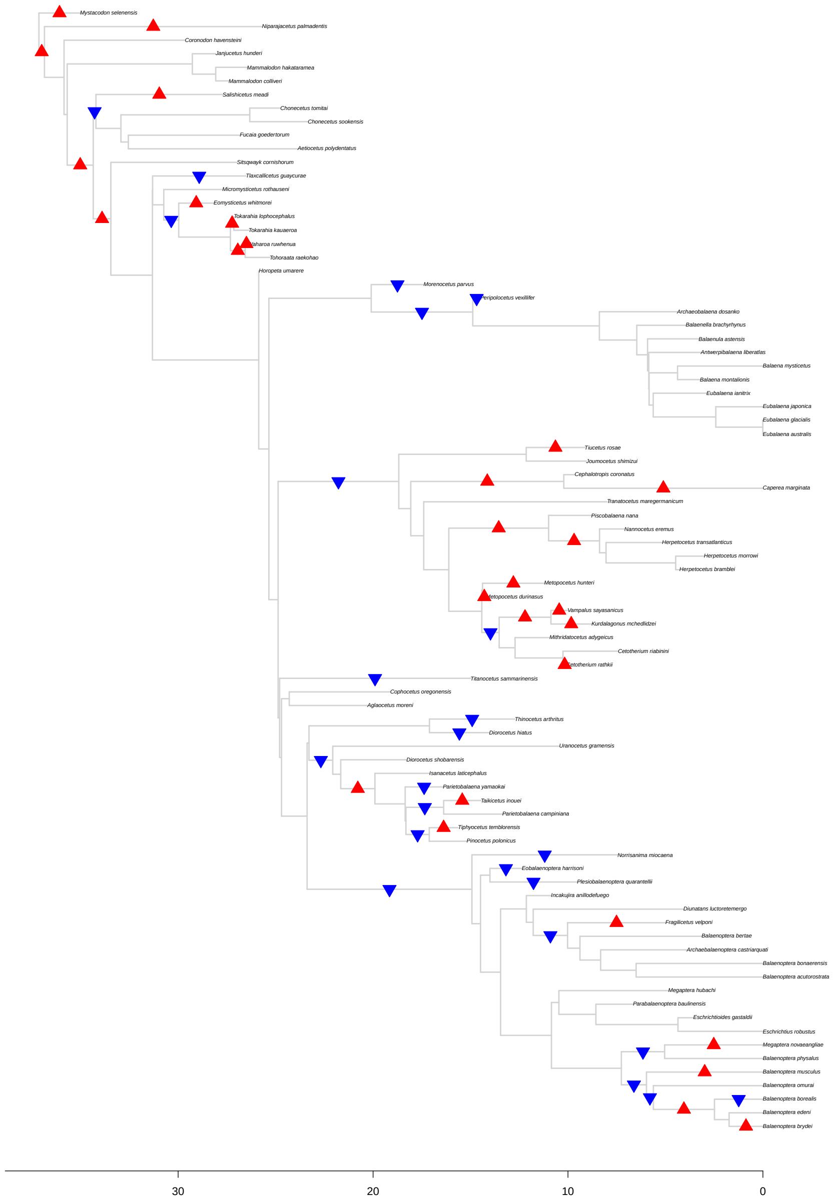

Figure S52: Results for *bayou* fit for the *Baleen* tree, excluding taxa on zero-length branches, setting the average number of shifts in the prior distribution ($\lambda$) to 15. The triangles represent the position and direction of the shifts with posterior probability higher than 0.1, with upward triangles (in red) indicating increases in $\theta$, and downward triangles (in blue) indicating decreases in $\theta$.

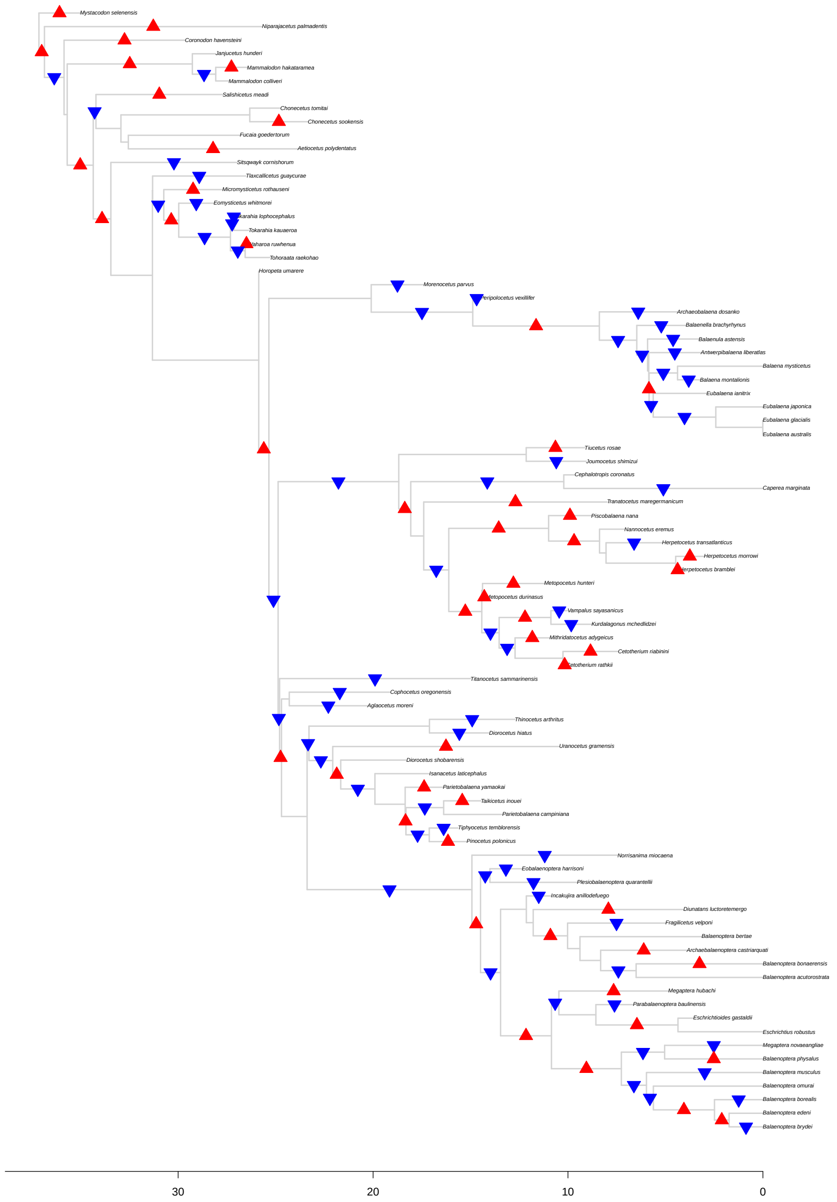

Figure S53: Results for *bayou* fit for the *Baleen* tree, excluding taxa on zero-length branches, setting the average number of shifts in the prior distribution ($\lambda$) to 50. The triangles represent the position and direction of the shifts with posterior probability higher than 0.1, with upward triangles (in red) indicating increases in $\theta$, and downward triangles (in blue) indicating decreases in $\theta$.

##### 3.5.6 *Toothed* tree

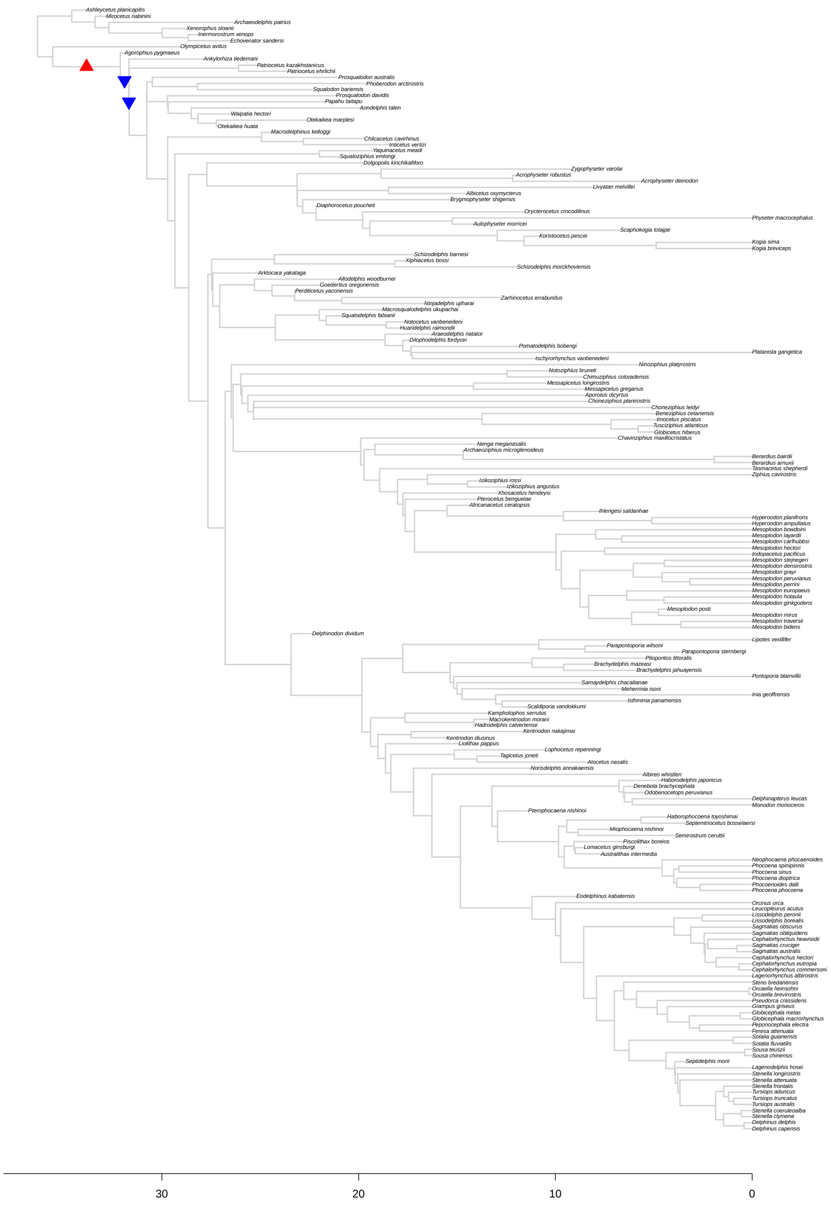

Figure S54: Results for *bayou* fit for the *Toothed* tree, excluding taxa on zero-length branches, setting the average number of shifts in the prior distribution ($\lambda$) to 5. The triangles represent the position and direction of the shifts with posterior probability higher than 0.1, with upward triangles (in red) indicating increases in $\theta$, and downward triangles (in blue) indicating decreases in $\theta$.

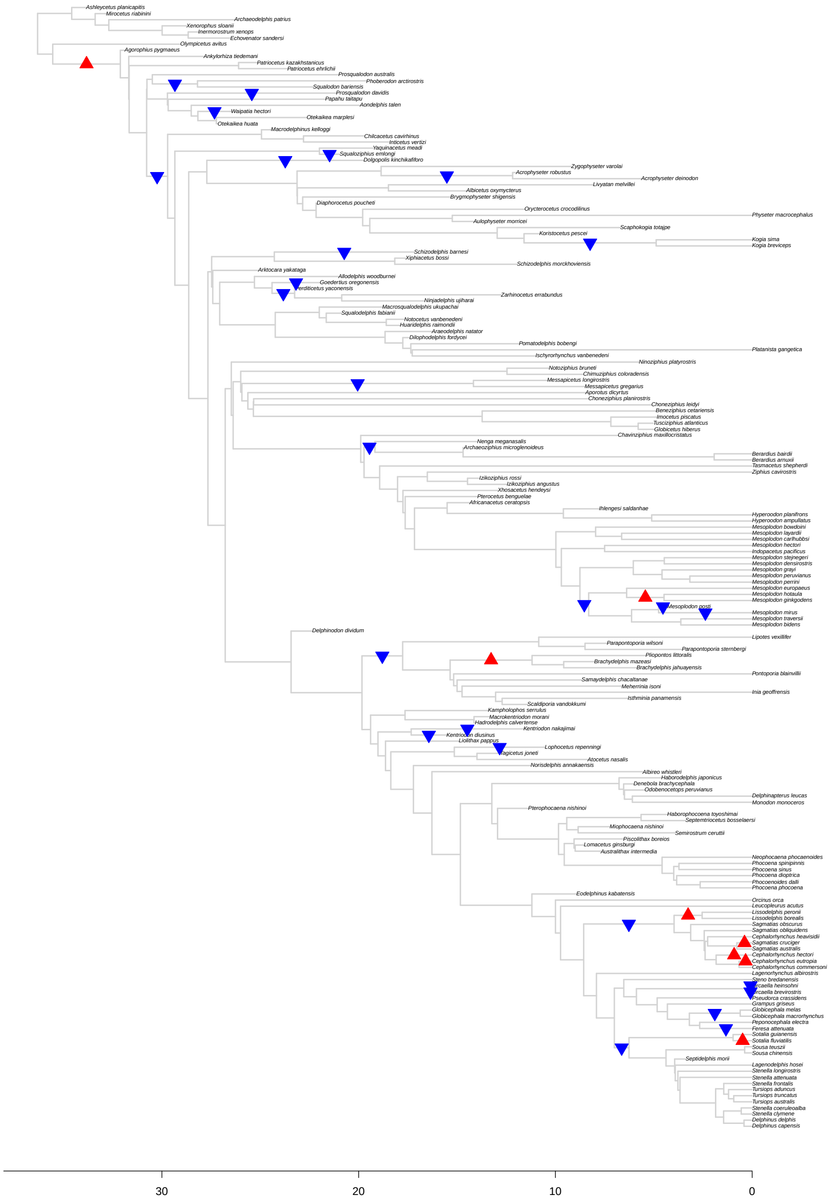

Figure S55: Results for *bayou* fit for the *Toothed* tree, excluding taxa on zero-length branches, setting the average number of shifts in the prior distribution ($\lambda$) to 15. The triangles represent the position and direction of the shifts with posterior probability higher than 0.1, with upward triangles (in red) indicating increases in $\theta$, and downward triangles (in blue) indicating decreases in $\theta$.

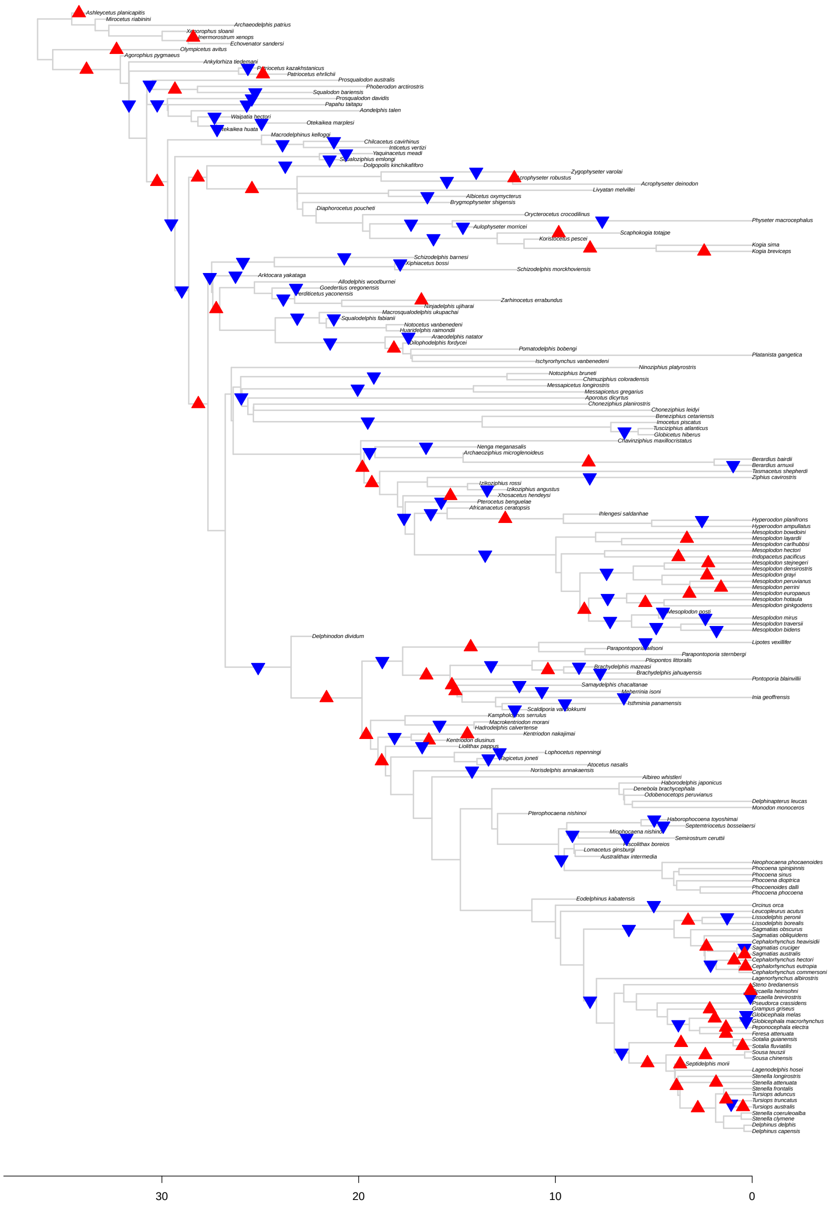

Figure S56: Results for *bayou* fit for the *Toothed* tree, excluding taxa on zero-length branches, setting the average number of shifts in the prior distribution ($\lambda$) to 50. The triangles represent the position and direction of the shifts with posterior probability higher than 0.1, with upward triangles (in red) indicating increases in $\theta$, and downward triangles (in blue) indicating decreases in $\theta$.
